# Biases in demographic modelling affect our understanding of recent divergence

**DOI:** 10.1101/2020.06.03.128298

**Authors:** Paolo Momigliano, Ann-Britt Florin, Juha Merilä

## Abstract

Testing among competing demographic models of divergence has become an important component of evolutionary research in model and non-model organisms. However, the effect of unaccounted demographic events on model choice and parameter estimation remains largely unexplored. Using extensive simulations, we demonstrate that under realistic divergence scenarios, failure to account for population size (*N*_*e*_) changes in daughter and ancestral populations leads to strong biases in divergence time estimates as well as model choice. We illustrate these issues reconstructing the recent demographic history of North Sea and Baltic Sea turbots (*Schopthalmus maximus*) by testing 16 Isolation with Migration (IM) and 16 Secondary Contact (SC) scenarios, modelling changes in *N*_*e*_ as well as the effects of linked selection and barrier loci. Failure to account for changes in *N*_*e*_ resulted in selecting SC models with long periods of isolation and divergence times preceding the formation of the Baltic Sea. In contrast, models accounting for *N*_*e*_ changes suggest recent (<6 kya) divergence with constant gene flow. We further show how interpreting genomic landscapes of differentiation can help discerning among competing models. For example, in the turbots data islands of differentiation show signatures of recent selective sweeps, rather than old divergence resisting secondary introgression. The results have broad implications for the study of population divergence by high-lighting the potential effects of unmodeleld changes in *N*_*e*_ on demographic inference. Tested models should aim at representing realistic divergence scenarios for the target taxa, and extreme caution should always be exercised when interpreting results of demographic modelling.

## Introduction

Since Alfred Wallace noted that “*Every species has come into existence coincident both in space and time with a pre–existing closely allied species*” (Wallace, 1855), understanding the processes by which new species arise (speciation) has been one of the major quests of evolutionary biology. In the case of sexual organisms, speciation can be defined as the evolution of reproductive isolation among populations, leading to distinct gene pools (Bolnick and Fitzpatrick, 2007). The study of population pairs where reproductive isolation is incomplete can therefore provide insight into the processes leading to the evolution of different species. According to the genic view of speciation (Wu, 2001), in the early stages of divergence reproductive isolation is a byproduct of differential adaptations and/or genetic incompatibilities, and therefore restricted to regions of the genome under exogenous and endogenous selection (often referred to as “barrier loci”). Such process results in heterogeneous differentiation across the genome, with barrier loci appearing as areas (“islands”) of higher differentiation due to selection, while unimpeded gene-flow homogenizes the rest of the genome (Nosil, 2012; Nosil and Feder, 2012; Ravinet *et al*., 2017; Roux *et al*., 2016). Partial reproductive isolation could arise via the gradual erosion of gene flow (primary divergence), for example because of multifarious selection across environments (Nosil, 2008; Nosil and Feder, 2012; Nosil *et al*., 2009). Alternatively, successive stages of strict isolation and secondary contact (secondary divergence) may facilitate the evolution of reproductive barriers among populations, whether they are due to ecological selection or to the evolution of genetic incompatibilities (e.g. Rougemont and Bernatchez, 2018; Rougeux *et al*., 2017; Roux *et al*., 2013, 2014). Both processes can lead to similar genomic landscapes of differentiation, as following secondary contact gene flow can erode genetic differentiation across the genome (with the exception of barrier loci and regions around them) to the point at which any signature of the initial stage of strict isolation is lost (Ravinet *et al*., 2017). Therefore, distinguishing whether divergence initiated in the presence or absence of gene flow, while important to understand how reproductive isolation arises, is not a trivial task.

Recent events of primary divergence and secondary introgression among ancient lineages are however expected to generate distinctive genomic landscapes surrounding barrier loci. Recent selection on a rare or novel mutations is likely to temporally reduce genetic diversity (π) surrounding barrier loci in the population experiencing selection (Smith and Haigh, 1974), revealing signatures typical of selective sweeps (e.g. Tavares *et al*., 2018). Absolute divergence among populations (*d*_*xy*_) will initially remain low, as in the early stages of divergence *d*_*xy*_ in regions surrounding a barrier locus is expected to reflect ancestral genetic diversity (see discussion in Ravinet *et al*., 2017). Instead, if barrier loci are of ancient origin, increased genetic differentiation (*F*_*ST*_) around barrier loci is likely to be driven by an increase in *d*_*xy*_ rather than a decrease in π (Cruickshank and Hahn, 2014). Indeed, there are several examples where islands of divergence that originated during long allopatric phases show both elevated *F*_*ST*_ and elevated *d*_*xy*_ with respect to the rest of genome, where the original signatures of divergence have been eroded by unimpeded gene flow (e.g. Duranton *et al*., 2018, 2020; Gagnaire *et al*., 2018; Nelson and Cresko, 2018). Unfortunately, the interpretation of genomic landscapes of differentiation is not always strait-forward (reviewed in: Ravinet *et al*., 2017), and it requires data (highly contiguous genome assembly) that are still lacking for most non-model organisms.

Demographic modelling provides a framework to reconstruct how gene flow has changed through the evolutionary history of diverging populations. Within the past two decades, several computational methods have been developed to reconstruct demographic history from genomic data. Such approaches usually rely on comparing summary statistics obtained from empirical data to simulations performed under competing divergence scenarios, of which the most commonly tested ones include isolation with continuous migration (IM), secondary contact (SC), strict isolation (SI) and ancient migration (AM) (Roux *et al*., 2013). Approximate Bayesian computation (ABC) approaches (Beaumont *et al*., 2002; Excoffier *et al*., 2013), as well as methods based on the diffusion approximation of the joint site frequency spectrum (jAFS) (*dadi* Gutenkunst *et al*., 2009) or its direct computation using a model of ordinary differential equations (*moments* Jouganous *et al*., 2017), have been broadly applied to test among competing demographic models of divergence. The models usually assume that an ancestral population of size *N*_*REF*_ gives rise to two populations of size *N*_1_ and *N*_2_ respectively at a time of split *T*_*S*_, after which several migration scenarios are contrasted. ABC, *dadi* and *moments* allow users to define complex demographic scenarios, explicitly modelling the effect of barrier loci—modelled as heterogeneous migration rates across the genome (Roux *et al*., 2013; Tine *et al*., 2014)—as well as the effect of linked selection-modelled as heterogeneous effective population size (*N*_*e*_) across the genome (Rougemont *et al*., 2017; Roux *et al*., 2016). Failing to account for heterogeneity in linked selection and migration rates may lead to biases in model choice and parameter estimation (Ewing and Jensen, 2016; Pouyet *et al*., 2018; Roux *et al*., 2014, 2016).

Such models have been used to test among competing gene flow scenarios across a broad range of divergence times, from a few thousands to millions of generations. For example, demographic modelling has been extensively used to test whether sympatric and parapatric lineages within environments that were shaped during the last glacial cycle arose via rapid ecologically driven divergence or are the result of postglacial secondary contact between more ancient lineages (e.g. Jacobs *et al*., 2020; Le Moan *et al*., 2016, 2019; Rougeux *et al*., 2017, 2019; Tine *et al*., 2014; Van Belleghem *et al*., 2018). The same approach has been extensively used to infer demographic models of divergence among incipient species that diverged hundreds of thousand to millions of generations ago (e.g. Bourgeois *et al*., 2019; Roux *et al*., 2013, 2014, 2016; Stuglik and Babik, 2016). Most of these studies concluded that contemporary heterogenous gene flow is a result of recent secondary contact (e.g. Gagnaire *et al*., 2018; Le Moan *et al*., 2016; Rougemont and Bernatchez, 2018; Rougemont *et al*., 2017; Rougeux *et al*., 2019; Roux *et al*., 2013, 2014, 2016; Tine *et al*., 2014) providing support to the hypothesis that initial allopatric phases of differentiation play a central role in the evolution of reproductive isolation (Roux *et al*., 2014), and hence primary divergence with gene flow due to ecological selection is rarer than suggested by some authors (Nosil, 2008). However, while these models can provide important insight into demographic history, they also show significant limitations. If the models tested are not close enough to the real divergence scenario, both model choice (for example, the choice between a SC and IM model) and parameter estimation may be affected. While great effort has been placed recentely to overcome potential biases due to barrier loci and linked selection (Bhaskar and Song, 2014; Gagnaire *et al*., 2018; Le Moan *et al*., 2016; Rougemont and Bernatchez, 2018; Rougemont *et al*., 2017; Roux *et al*., 2013, 2016; Tine *et al*., 2014), it remains unclear how unmodelled demographic events, such as size changes in both ancestral and daughter populations, may affect model choice and parameter estimation.

Most recent studies of non-model organisms, where prior knowledge of past demographic events is limited, assume that divergence starts from an ancestral population at mutation-drift equilibrium, with an instantaneous split into two populations of constant size (as in Roux *et al*., 2013). If, for example, there was an unmodelled size change in the ancestral population (such as a population expansion or contraction), we can expect an overestimation of divergence time, since the models allows changes in *N*_*e*_ only at time *T*_*S*_, pushing estimates of *T*_*S*_ towards the time of ancestral population size change. Similarly, unmodelled bottlenecks followed by exponential growth in one of the daughter populations may bias estimates of *T*_*S*_, as small populations experience faster genetic drift (potentially leading to overestimate recent divergence). Both bottlenecks (Luikart *et al*., 1998) and SC (Alcala *et al*., 2016) are expected to generate an excess of middle frequency variants, and hence an unmodelled bottleneck could bias model choice towards SC. It is less clear how changes in *N*_*e*_ in ancestral populations affect model choice, as to the best of our knowledge no one has addressed this question. This is a matter of concern, as few studies explicitly model growth in daughter populations, and very few studies of non-model organisms have tested for changes in ancestral population size. A brief search in Web of Science for published studies using demographic modelling in non-model organisms (using combinations of keywords “Isolation with Migration”, “Secondary Contact”, “dadi”, “abc”, “fastsimcoal”) suggests that less than one fifth of papers published between 2016 and 2020 years accounted for changes in *N*_*e*_ in the ancestral population or in at least one of the daughter populations.

Here, we used both simulations and empirical data to demonstrate that unmodelled demographic events in both ancestral and daughter populations can strongly affect both model choice and parameter estimation. Using coalescent simulation of IM and SC scenarios under the Wright-Fisher neutral model, we demonstrate that failure to account for changes in *N*_*e*_ in ancestral and daughter populations leads to extreme biases in estimates of *T*_*S*_ and to a strong bias towards the choice of SC models. We then reconstruct the demographic history of the Atlantic and Baltic Sea populations of the turbot *Schoptalmus maximus*, which a recent study (Le Moan *et al*., 2019) suggested have diverged at the start of the last glacial maximum (>50 kya) and experienced secondary contact following the end of the last glaciation. We argue that these inferences were likely biased because of the failure to account for a past demographic expansion (which leads to overestimate *T*_*S*_) and to model a bottleneck during the invasion of the Baltic Sea (leading to the erroneous choice of a SC model). Furthermore, we show that genomic patterns of differentiation are also consistent with a scenario of very recent divergence with gene flow. We discuss the potential implications of our findings for inferring the demographic history of non-model organisms in general.

## Results

### Analyses of simulated data

For all simulated data (Fig. 1A&B), we optimized parameters for the simple IM and SC models and tested, using Akaike weights (*W*_*AIC*_ see Material and Methods) the support for the correct gene flow scenario (IM). For the recent divergence scenarios, we tested (and optimized parameters for) all basic demographic models (Fig.1 C), and used *W*_*AIC*_ to test the support for the correct gene flow scenario (SC or IM depending on simulation). Details of all models used for demographic inference are given in Supplementary Table 1.

**FIG. 1.**
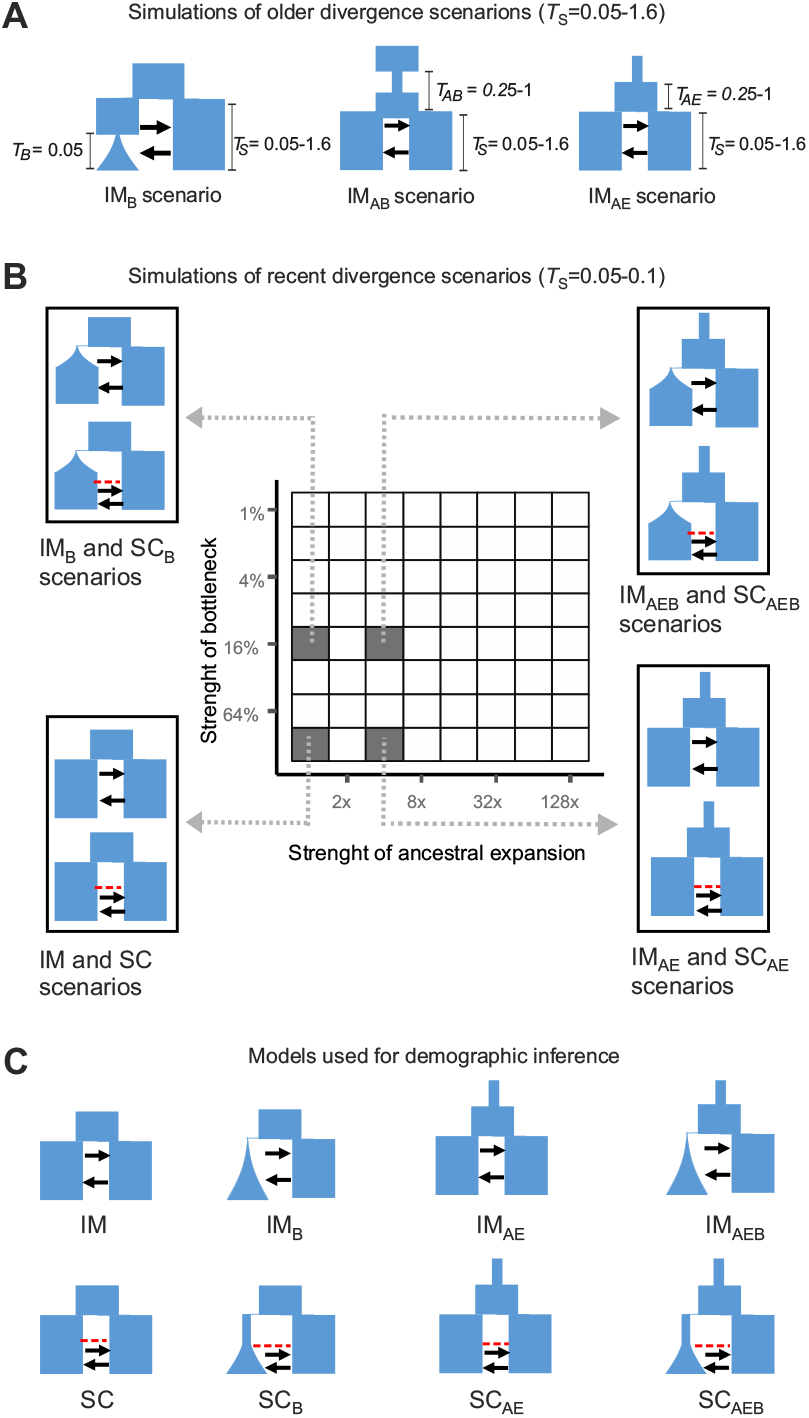
Simulation scenarios and models used for demographic inference. Times are given in units of 4*N*_*REF*_ generations. **A**: We simulated data under an IM scenario with divergence times from 0.05 to 1.6. In IM_B_ scenarios one population experiences a contraction to 1-64% of its previous size at time *T*_*B*_ followed by exponential growth. The IM_AB_ scenario reflects an ancestral population size contraction followed by expansion. In the IM_AE_ scenario, the ancestral population undergoes a population expansion. For both AB and AE scenarios, we tested a range of times (0.25, 05, 1) and strengths of the ancestral contraction (AB= 1/4, 1/16, 1/64 of previous size) and expansion (AE=4×, 16×, 64× the ancestral *N*_*e*_). **B**: simulations of recent divergence scenarios, with a *T*_*S*_ of 0.05 for IM scenarios and 0.1 for SC scenarios. Data were simulated under the IM and SC scenarios with 64 possible combinations of ancestral expansion and bottlenecks. Simulations were ran both with symmetric and asymmetric migration, for a total of 256 demographic scenarios. **C**: demographic models used for inference with *moments* and *dadi*, i.e. the basic IM and SC models as well as modifications that include ancestral population size changes as well as bottlenecks followed by growth in one of the daughter populations. Migration rate is asymmetric (i.e. different *m*_12_ and *m*_21_ parameters) in all inference models. For the analyses of the empirical data, we used the models graphically represented in panel **C**, and modifications of these models accounting for heterogeneity in migration rates (2M), *N*_*e*_ (2N models) across the genome, as well as modifications accounting for both (2N2M models).

### Biases in IM and SC models in older divergence scenarios

An unmodelled recent bottleneck always led to severe biases in model choice (favoring SC model, as shown by low *W*_*AIC*_) and estimates of *T*_*S*_, even when the simulated bottleneck was mild (Fig. 2 C). Estimates of *T*_*S*_ tended to reflect the strength of the bottleneck rather than divergence time (Fig. 2 B), suggesting the effects of a recent bottleneck on the joing allele frequency spectrum (jAFS) obscure signatures of divergence. This led to a severe overestimation of *T*_*S*_ in recent divergence scenarios (simulation *T*_*S*_ = 0.05) and a severe underestimation of *T*_*S*_ in older divergence scenarios (simulation *T*_*S*_ *>* 0.4). The estimated time of strict isolation as a proportion of divergence time (*T*_*SI*_*/T*_*S*_) is a positive function of the strength of the unmodelled bottleneck (Fig. 2 D).

**FIG. 2.**
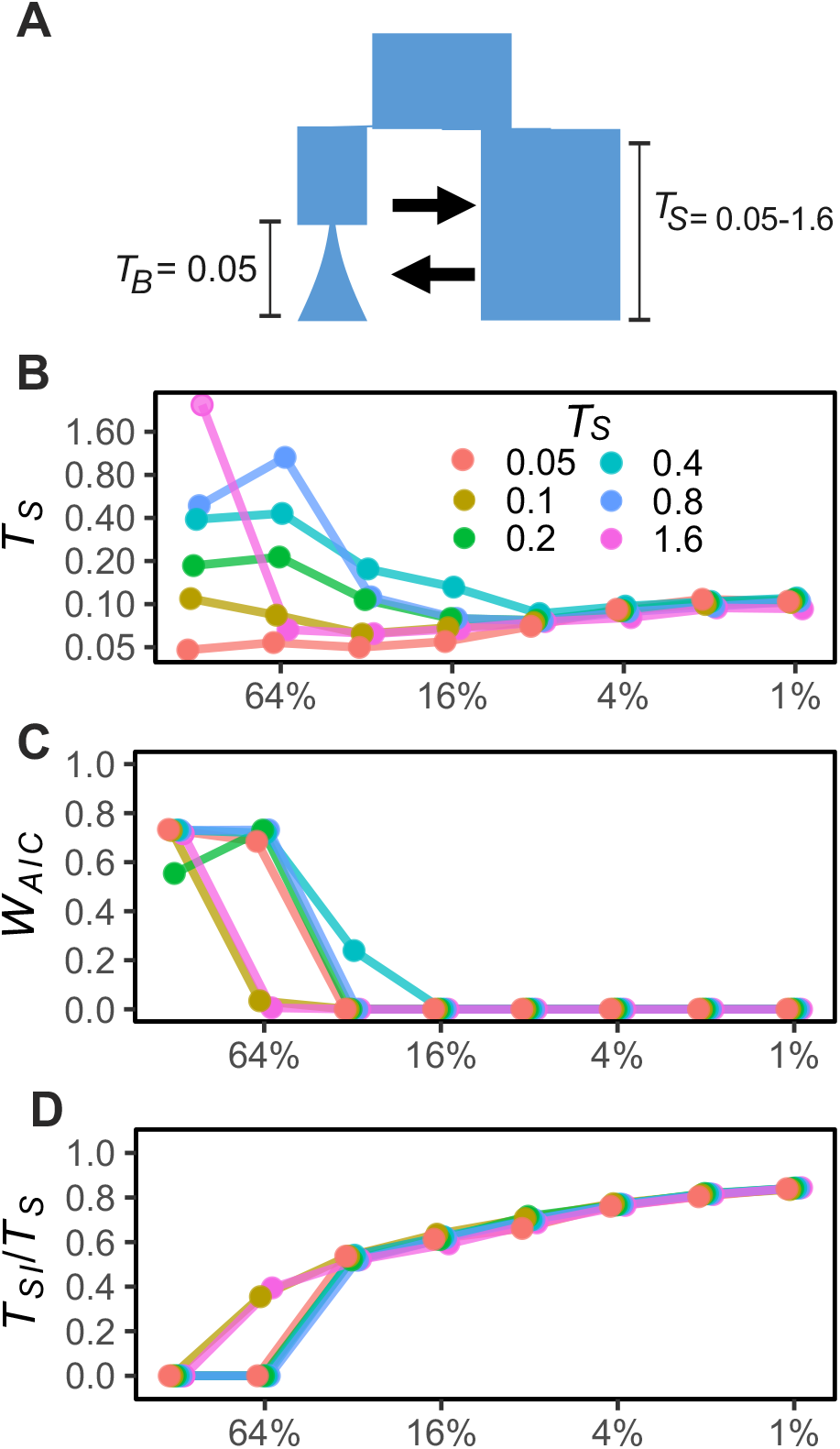
Effects of unmodelled bottlenecks in a daughter population on model choice and parameter estimates. Model choice and parameter estimates for the simple IM and SC models for simulations with *T*_*S*_ = 0.05−1.6 and recent bottlenecks of different strengths at time *T*_*B*_ = 0.05. Panel **A** shows the simulation model, panel **B** show estimated time of divergence *T*_*S*_, panel **C** shows Akaike weights (*W*_*AIC*_) for the correct model (IM) and panel **D** shows the inferred time of strict isolation (*T*_*SI*_) as a proportion of total divergence time.

Non equilibrium states in the ancestral population also led to biases in both model choice and parameter estimation (Fig. 3), but the severity of these biases were affected by both *T*_*S*_ and the time before *T*_*S*_ at with the last ancestral change in *N*_*e*_ occurred (*T*_*AE*_ for AE models and *T*_*AB*_*/*2 for B models). When *T*_*S*_ was relatively shallow (*T*_*S*_ < 0.4) unaccounted changes in *N*_*e*_ in the ancestral population led to overestimate *T*_*S*_. This bias was most severe when the last ancestral change in *N*_*e*_ (*T*_*AE*_ in AE model, and *T*_*AB*_*/*2 in AB models) occurred 0.5×4*N*_*REF*_ generations before *T*_*S*_. Biases towards the choice of SC models were most severe at intermediate *T*_*S*_, while the effects of not accounting for non-equilibrium in the ancestral population becomes irrelevant when *T*_*S*_ *>* 0.8 (Fig. 3).

**FIG. 3.**
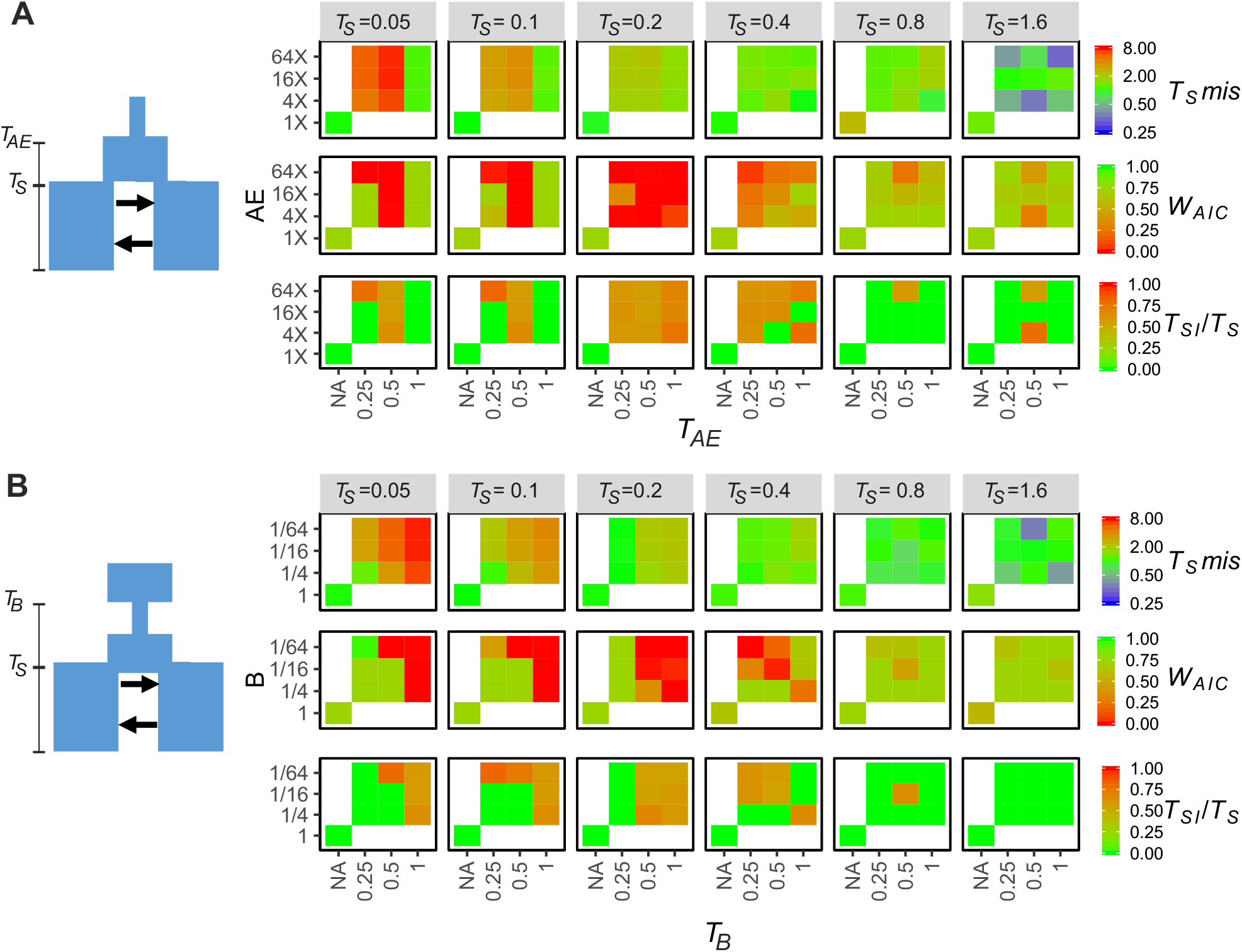
Effect of unmodelled demographic changes in the ancestral population on model choice and parameter estimates. In this figure we report biases in model choice and parameter estimation of simple IM and SC models for IM simulations with *T*_*S*_ = 0.05 − 1.6 including ancestral demographic expansion (**A**) and an ancestral bottleneck followed by expansion (**B**). On the left of panels **A & B** is a graphical representation of the simulation model. In panel **A** the y axis represents the ancestral population size after the expansion as a multiplier of its size preceding the expansion, and the x axis represents the time before *T*_*S*_ at which the ancestral expansion happened (*T*_*AE*_). In panel **B** the y axis represents the size of the ancestral population after contraction as fraction of the population size preceding contraction and the x axis represent the time before *T*_*S*_ at which the ancestral bottleneck happened (*T*_*B*_). Times are given in units of 4*N*_*REF*_ generations. For **A & B** and for each simulated *T*_*S*_ we present the misestimation of *T*_*S*_ (*T*_*S*_ *mis* = *T*_*S*_ *estimated/T*_*S*_ *simulated*), the weight of evidence (*W*_*AIC*_) for the correct (IM) gene flow scenario as well as the estimated time of strict isolation as a proportion of total diverging time (*T*_*SI*_ */T*_*S*_). In each graph, the bottom-left square represents estimates for simulations with no ancestral changes in *N*_*e*_.

### Biases in IM and SC models in recent divergence scenarios

For the simulations of recent divergence scenarios, we conducted analyses based on simulations of 1 million or 100 000 loci. These simulations gave similar results (compare Fig. 4 and Supplementary Fig. 2 with Supplementary Figs. 16 & 17 respectively), with the exception that there was more stochastic variation when analyzing the smaller data sets. Hence, we report results based on the larger data sets in the main body of the manuscript, while the results for the smaller numbers of loci are presented in the supplementary materials. Models generally converged very well with the optimization scheme utilized (see examples in Supplementary Figs. 3-10).

**FIG. 4.**
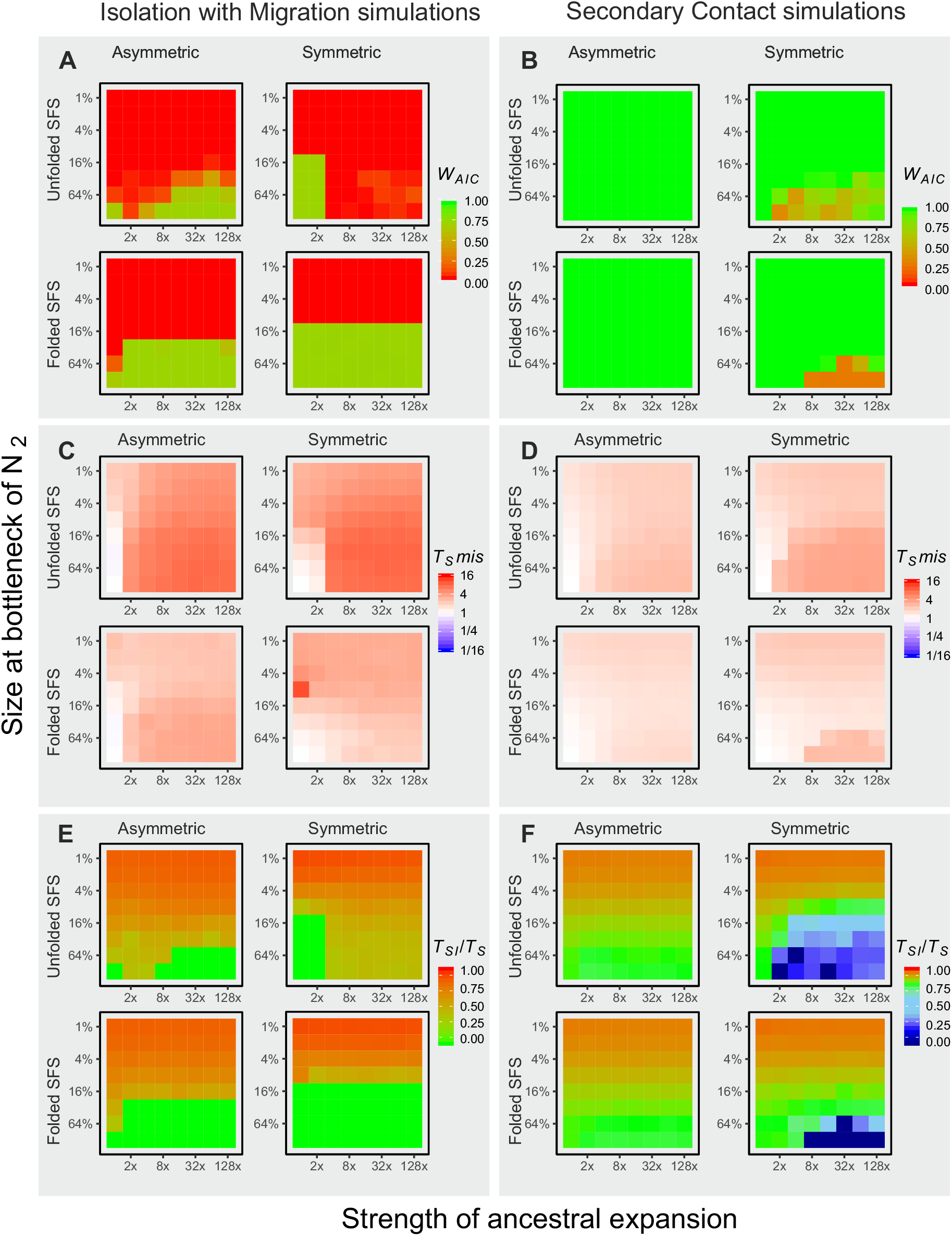
Model choice and parameter misestimation for the simple IM and SC models for all simulations of the larger data sets (1 million loci). Left panels (**A, C, E**) show results from simulations with constant migration (IM), right panels (**B, D, F**) show results from simulation with a period of strict isolation (SC). Within each panel, results are shown for simulations with symmetric and asymmetric migration, and for inference using the folded or unfolded jAFS. Within each panel, each graph represent the values for all 64 simulations as per Fig. 1C. Panel **A** and **B** show weight of evidence for the correct gene flow scenario (0-1). Panel **C** and **D** show misestimation of the parameter *T*_*S*_ (*T*_*S*_ estimate /*T*_*S*_ of simulation). Panels **E** and **F** show the estimated proportion of the divergence time for which the model inferred strict isolation (green represents the correct time, i.e. 0 for IM model and 0.75 for SC models).

Parameter estimation and model choice from basic IM and SC models were severely biased when the demographic scenario of the simulations deviated from the models (Fig. 4). While for SC simulations the underlying gene flow scenario was almost always identified (Fig. 4 B), when the true divergence scenario of the simulations had continuous gene flow even moderate unmodelled bottlenecks resulted in strong support for SC (Fig. 4 A). Stronger bottlenecks were associated with an overestimation of divergence times, and the bias was much more severe for simulations with constant gene flow (IM; Fig. 4 C&D, Supplementary Fig. 11 A&B). There was also a clear relationship between the length of the estimated period of strict isolation (SI) in IM simulations and the strength of the unmodelled bottleneck (Fig. 4 E). Hence, in recent demographic scenarios unmodelled bottlenecks generally led to choice of SC models with long periods of strict isolation even when the simulation scenario had constant gene flow. Unmodelled size changes in the ancestral population led to severe overestimates of divergence time in IM scenarios (Fig. 4 C) and to a lesser degree in SC scenarios (Fig. 4 D). The stronger the unmodelled ancestral expansion, the more estimates of *T*_*S*_ approached *T*_*AE*_ (which is 10×*T*_*S*_ under IM scenarios, 5× *T*_*S*_ under SC scenarios, see methods; Fig. 4 C&D, Supplementary Fig. 11 A&B). Unmodelled ancestral expansions also biased model choice towards SC when the true scenario had constant migration, but only when the unfolded jAFS was used for demographic inference. However, when an unmodelled ancestral expansion led to incorrectly choose the SC model, the proportion of time of SI inferred was low (< 50% of the total divergence time) unless the simulation included also a strong bottleneck (Supplementary Fig. 12 A&B). Interestingly, in SC simulations without strong bottlenecks, an unmodelled ancestral expansion led in some cases to a bias towards IM models (Fig. 4 B), and in general to an underestimate of the proportion of SI (Supplementary Fig. 12 A&B). Not surprisingly, unmodelled ancestral expansions led also to strong overestimates of ancestral population size *N*_*ANC*_ (Supplementary Fig. 13).

All the above reported biases were more severe when the unfolded jAFS was used for demographic inference. Particularly, *T*_*S*_ misestimation was roughly twice as high when inference was carried out using the unfolded jAFS compared to the folded jAFS (Fig. 4 C&D, Supplementary Fig. 11 A&B). Similarly, bias in model choice towards SC models was stronger when using the unfolded jAFS (Fig. 4A and Fig. 4, Supplementary Fig. 11). It should be noted, however, that estimates of contemporary *N*_*e*_ and migration rates were always fairly accurate (see Supplementary Figs. 14 & 15). Furthermore, in general these biases were less severe when the smaller data set was used for inference (Supplementary Fig. 16). These biases were not unique to the simulation engine we used, as the analyses of a subset of the data using *dadi* gave nearly identical results (Supplementary Fig. 18). Gross model mis-specifications were, however, identifiable by inspecting the jAFS residuals produced by *dadi* and *moments* (Supplementary Fig. 19)

### Biases in 8-model comparisons in recent divergence scenarios

Testing demographic scenarios that more closely approximate the real demographic history of simulated populations resulted in much less severe biases in model choice. Weight of evidence for the correct gene flow scenario was much stronger, with all simulations of SC scenarios being correctly identified and IM simulations being sometimes misidentified as SC only when population two experienced very severe bottlenecks followed by exponential growth (Supplementary Fig. 2 A). However, when SC scenarios were incorrectly chosen, the length of the inferred periods of strict isolation was usually very short (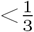 of total divergence), suggesting that IM and SC models were converging towards the same demographic history (Supplementary Fig. 2 C, Supplementary Fig. 12 C&D). While ancestral expansions and bottlenecks still caused a systematic bias towards overestimating divergence times, this bias was negligible (Supplementary Fig. 2B, Supplementary Figs. 11 C&D).

### Analyses of empirical data

#### Population genetics analyses

A PCA based on genotype likelihoods from 9063 biallelic SNPs with MAF > 0.02 shows a clear partitioning along the 1^*st*^ PC between the North Sea and Baltic Sea individuals (Fig. 5 A&C), with several individuals from the transition zone (locations VD and OS in Fig. 5 C) showing intermediate genotypes. FastSTRUCTURE analyses (Fig. 5 B) shows very similar results, with two clear genetic clusters (North Sea and Baltic Sea) and individuals from VD and OS (the transition Zone) exhibiting high proportion of admixture. Based on these results, individuals from admixed populations (VD and OS) where excluded from further analyses, with the exception of genome scans for selection using PCAngsd (since this analysis does not assume discrete populations).

**FIG. 5.**
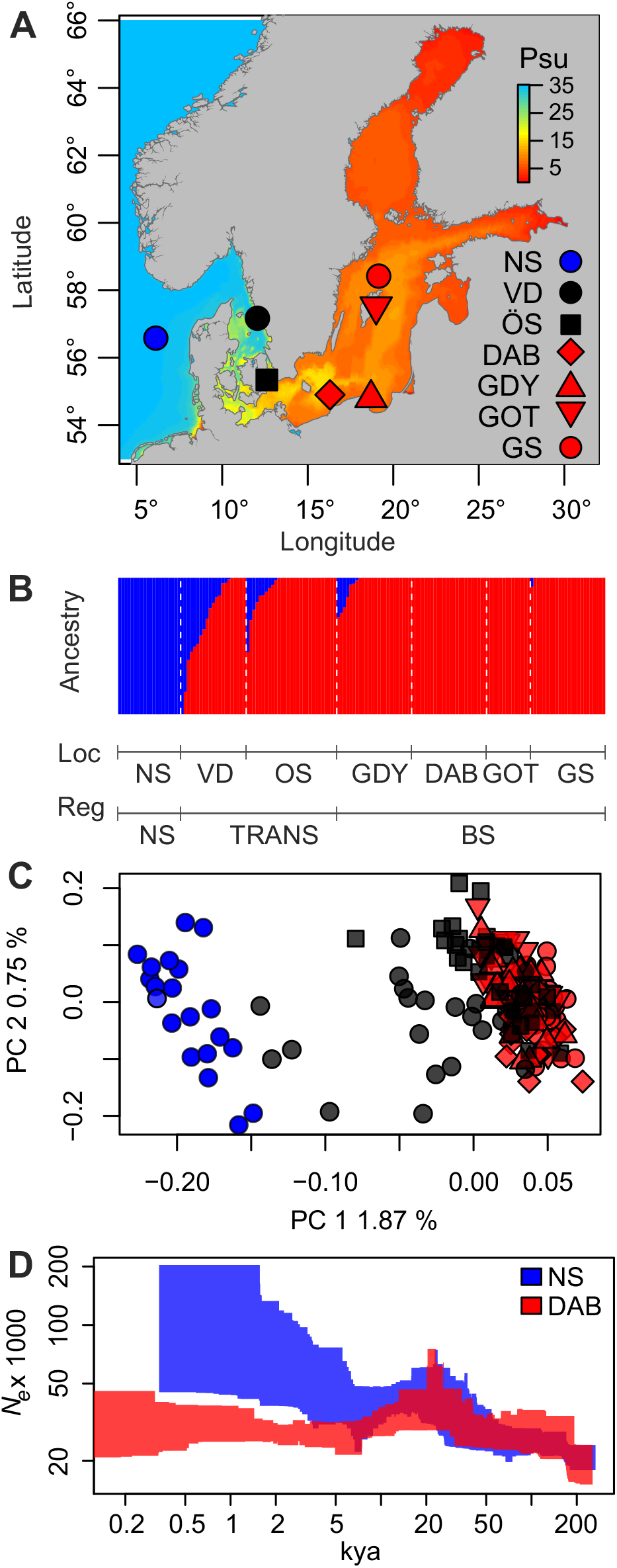
Sampling sites, genetic structure and historical change in *N*_*e*_ of *Schophtalmus maximus* populations in the North Sea (blue) the transition zone (black) and Baltic Sea (red). **A** Sampling locations showing modelled bottom salinity of the Baltic Sea. **B** Individual ancestry reconstructed from fastSTRUCTURE. **C** PCA performed from genotype likelihoods of 9063 bi-allelic SNPs, color and population codes as per **A. C** Changes in *N*_*e*_ across time inferred for a representative population from the North Sea (NS) and the Baltic Sea (GOT). Polygons represent 95% confidence intervals.

#### Genomic landscape of differentiation

Average differentiation between the North Sea and Baltic Sea populations was weak (mean *F*_*ST*_ = 0.017), however several SNPs across the genome showed marked *F*_*ST*_, notably SNPs located around the center of chromosome 1 and in chromosome 13 (Fig. 6A). Genome scans, performed using the extended model of FastPCA (Galinsky *et al*., 2016) implemented by PCAngsd and a Hidden Markov model (HMM) approach to detect genomic islands (Hofer *et al*., 2012; Marques *et al*., 2016), identified 32 outlier loci with a false discovery rate <0.01, located in 15 distinct genomic regions (Fig. 6B). Most of these 15 regions included one or two outlier loci, and were located at the very end of chromosome arms (Supplementary Fig. 20), were the chance of false positives is highest. However, two genomic islands, located in chromosome 1 and 13, had several outlier SNPs (10 and 13, respectively) spanning a distance of over 1Mb and showing extreme levels of differentiation (Fig. 6 A, B, C & D).

**FIG. 6.**
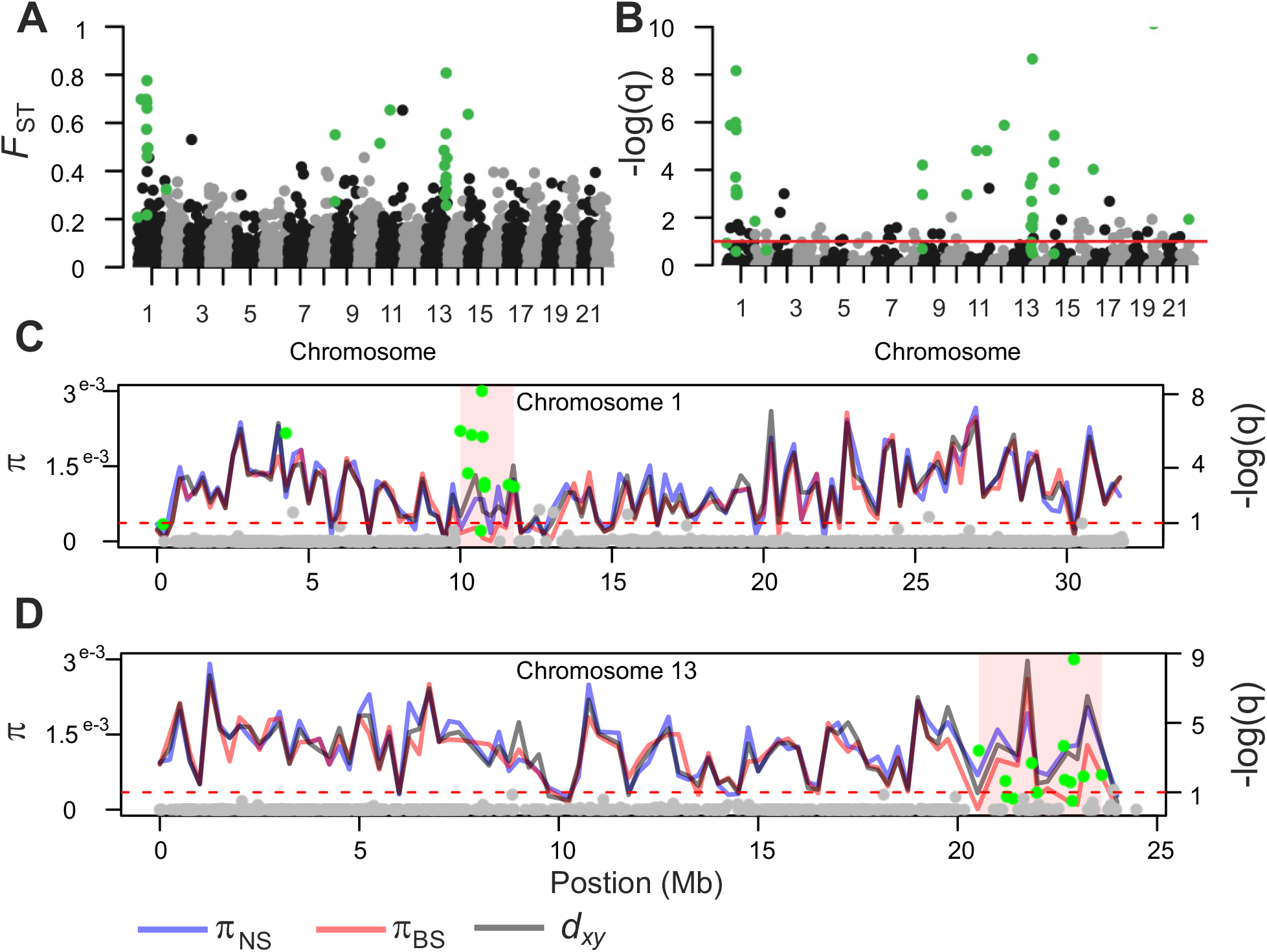
Genomic landscape of differentiation between North Sea and Baltic Sea turbots. **A** per site *F*_*ST*_ from hard-called SNPs across the genome, SNPs significant in the HHM test are shown in green. **B** q-values (FDR) on a negative log scales from fastPCA genome scan for selection. The red line marks a FDR of 0.1, and green circles above this line denote SNPs which are significant based on a FDR threshold of 0.1 as well as based on results from the HHM test. **C** patterns of genetic diversity across Chromosome 1 calculated in non-overlapping windows of 250 kb. The red line represents π in the Baltic Sea, the blue line π in the North Sea, and the black line represents *d*_*xy*_. Dots along the chromosomes represent q-values (FDR) on a negative log scales from fastPCA genome scan for selection for each individual SNP, the dotted line demarks the 0.1 FDR and green dots represents significant outliers according to the HHM test. **D** same as **C** but for Chromosome 13

Levels of genetic diversity (π) and absolute divergence (*d*_*xy*_) were highly correlated across the genome (Fig. 6 C & D, Supplementary Fig. 20). *d*_*xy*_ was highly correlated with π in the North Sea (*R*^2^ = 0.85, p < 1^*e*−16^), which is the expectation since in early stages of divergence *d*_*xy*_ is expected to approximate π in the ancestral population, which is likely still represented by the North Sea. The correlation between π in the North Sea and Baltic Sea population breaks down within the two genomic islands of differentiation in chromosome 1 and 13, where increased allelic differentiation (*F*_*ST*_) is driven by a dramatic decrease in genetic diversity in the Baltic Sea, rather than an increase in absolute divergence (Fig. 6 C & D). Such genomic landscapes are classic signatures of recent selective sweeps acting on novel or rare variants, resulting in a transient loss of genetic diversity surrounding the site of selection.

#### Demographic modelling

Stairway plots revealed that Baltic Sea and North Sea turbots shared a similar demographic history until 10-20 kya, with evidence of a population expansion between 20 and 100 kya (Fig. 5D). The Representative population for the Baltic Sea (GOT) show signs of population contraction followed by growth following the end of the last glaciation, and lower contemporary *N*_*e*_ compared to NS. We also tested three simple 1-population models using *moments* on all individuals from the NS and BS separately: a Standard Neutral Model (SNM, assuming constant population size at equilibrium), a 2 Epochs model (2EP, assuming a sudden population change at time T1) and a 3 Epochs model including a sudden demographic change at time *T* 1 followed by a bottleneck at time *T*_2_ and exponential growth (2EP_B_). The results give strong support for population size changes, rejecting the SNM neutral mode for both populations. In the North Sea the 2EP model had stronger support than the 2EP_B_ model (*W*_*AIC*_ of 0.88 and 0.12, respectively) while for the Baltic Sea population both models had similar support (0.57 and 0.43, respectively). One-population models provide guidance in selecting realistic demographic scenarios to test in more complex models but should otherwise be interpreted with extreme caution as they ignore the effects of gene flow. Here results from both stairway plots and one-population models performed in *moments* suggest that both an ancestral population expansion and a bottleneck followed by growth in the Baltic Sea population should be formally tested in two-population models.

The folded jAFS used for testing two-population models included 16270 biallelic SNPs. We tested the eight basic scenarios presented in Fig. 1 C (IM, SC, IM_B_, SC_B_, IM_AE_, SC_AE_, IM_AEB_ and SC_AEB_) as well as modifications of these four basic models accounting for heterogeneous migration rate (2M models) and *N*_*e*_ (2N models) across the genome as well as a combination of the two (2M2N models). Model choice and parameter estimates in two-population models were strongly affected by the inclusion or exclusion of ancestral expansions and bottlenecks (Fig. 7). Scenarios including ancestral population expansions (AE) and bottlenecks (B) had lower AIC than simple IM and SC models, with AEB models performing best (Fig. 7 A). Within basic IM and SC models, SC models fitted the data significantly better showing much lower AIC than IM models. However, as more complex demographic scenario (AE and B models) were compared, the difference in AIC between IM and SC models became smaller, with IM models showing the lowest AIC in AEB scenarios. Interestingly a very similar pattern was observed in our simulations. In Fig. 7 E we show the AIC of IM and SC models (and their AE, B, and AEB variations) for a simulation with an ancestral expansion (*N*_*AE*_ = 2×*N*_*ANC*_) and bottleneck (where *N*_2_ at time of split is 4% of current *N*_2_). Models accounting for heterogenous migration rates (2M) and *N*_*e*_ (2N) across the genome fitted the data better (Fig. 7A), but the inclusion or exclusion of these parameters in the models did not change the relative support to IM and SC models.

**FIG. 7.**
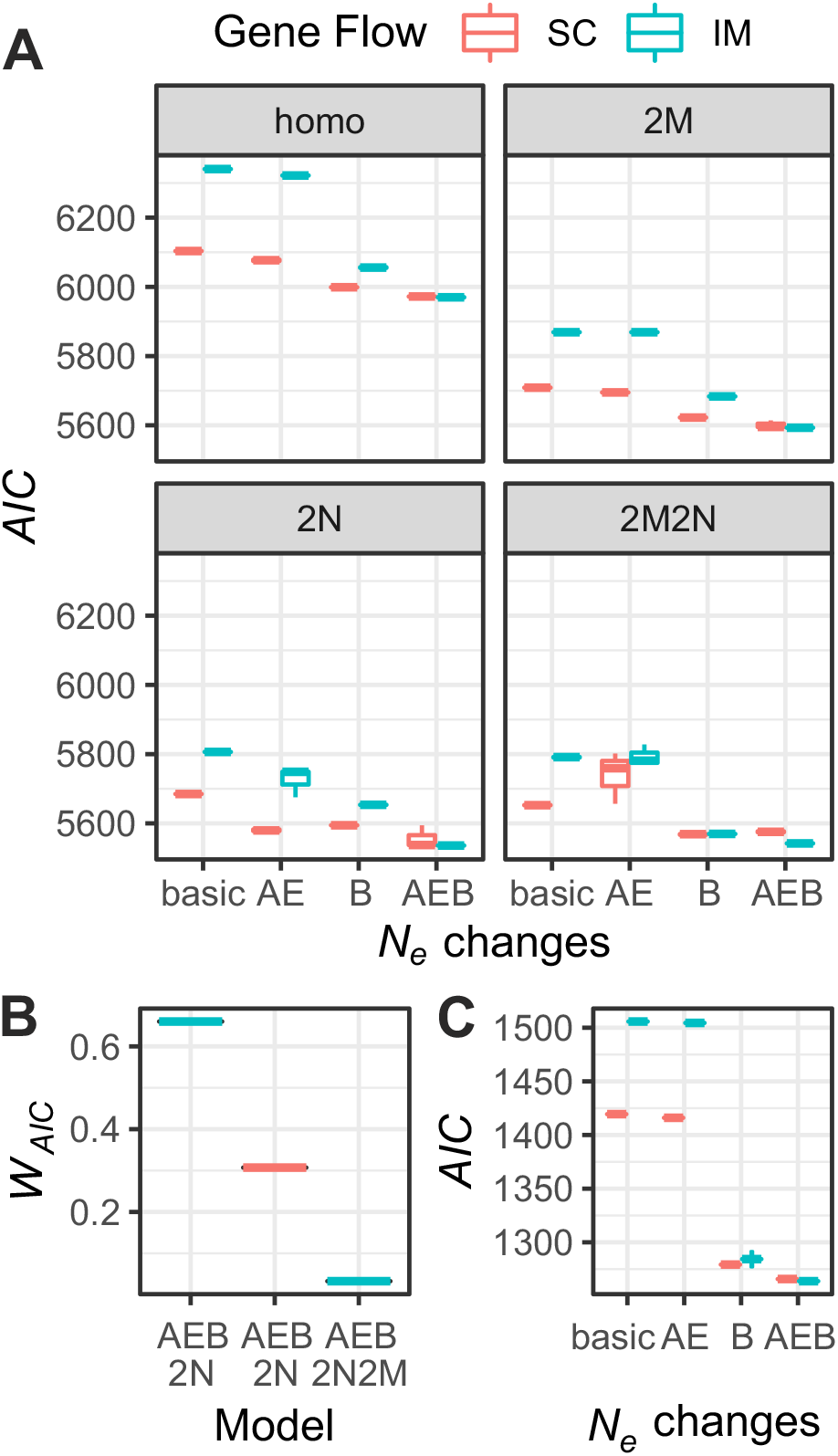
Results from the 32 models optimized for the *S. maximus* data (**A & B**) and data simulated with similar parameters (**C**). **A** shows AIC from the best 3 replicates of SC (red boxplots) and IM (blue boxplots) models for any combination of ancestral expansion/bottleneck (AE, B and AEB models). Top right panel (Homo): homogenous gene flow and *N*_*e*_. Panel 2M: heterogeneous migration rates across the genome. Panel 2N: heterogenous *N*_*e*_ across the genome. **B** *W*_*AIC*_ of the three best models. **C** Results from simulations representing a scenario similar to the best inferred model (IM scenario with *N*_*AE*_ = 2× *N*_*ANC*_ and a bottleneck in population two at time *T*_*S*_ to 4% of the current *N*_*e*_)

The two best fitting models were IM_AEB2N_ and SC_AEB2N_, with *W*_*AIC*_ of 0.66 and 0.31, respectively (Fig. 7C). However, these two models converged to approximately the same scenario (Fig. 8). Both models suggest an ancestral population expansion approximately 30-100 ky before divergence to about 1.7 times the ancestral size, a colonization of the Baltic Sea < 5.5 kya, and a strong reduction in *N*_*e*_ at the time of colonization followed by growth in the Baltic Sea population. The estimate of the period of strict isolation (*T*_*SI*_) from the SC model is very small (0.6 ky, Fig. 8 D). Formal testing among these two competing, nested models using a Likelihood Ratio Test (LRT), and adjusting the D statistics to account for possible effects of linkage (Coffman *et al*., 2015), gives no statistical support for the SC_AEB2N_ model (*D*_*adj*_=0.1293, *p*= 0.3596). LRT tests, on the the hand, provide support for the inclusion of a bottleneck in the model (LRT test between IM_2N_ and IM_B2N_: *D*_*adj*_ = 20.8501,*p*< 1^−4^) and of an ancestral expansion (LRT test between IM_2N_ and IM_AE2N_: *D*_*adj*_ = 7.4156,*p* = 0.0155). Unscaled parameters for the two best models and their standard deviation estimated using the Fisher Information Matrix and the Godambe Information Matrix (Coffman *et al*., 2015) are given in Supplementary Table 3.

**FIG. 8.**
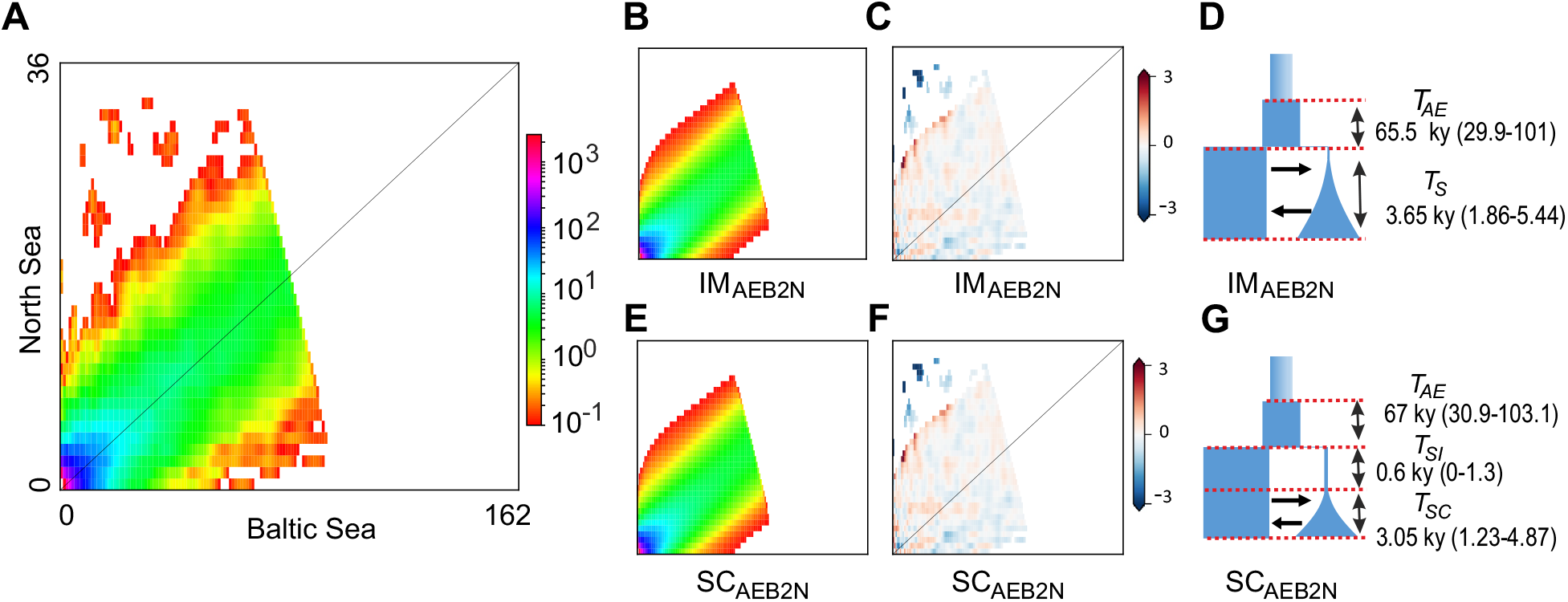
Inferred demographic history and model fit from the two models with highest support. Observed jAFS (**A**), modelled jAFS (**B & E**), model residuals (**C & F**) and parameters (**D & G**) for the two best models. **B** modelled jAFS, **C** residuals and **D** graphical representation of the Isolation with migration model including a bottleneck in daughter population, an ancestral change in *N*_*e*_ and heterogenous *N*_*e*_ across the genome (IM_AEB2N_ model) as well as estimated time parameters with 95% Confidence Intervals. **E** modelled jAFS, **F** residuals and **G** graphical representation of the secondary contact model including a bottleneck in daughter population, an ancestral change in *N*_*e*_ and heterogenous *N*_*e*_ across the genome (SC_AEB2N_ model) as well as estimated time parameters with 95% Confidence Intervals estimated from the Godambe Information matrix.

## Discussion

Reconstructing the demographic history of diverging populations is of central importance to understanding the role of gene flow and periods of strict isolation in shaping the process of speciation. Here we demonstrate that when the demographic history of the simulated taxa deviates from the tested scenarios, model choice and parameter estimation can be severely biased. Specifically, unmodelled changes in both ancestral and daughter populations led to biases in estimates of divergence times and to favor scenarios that include periods of strict isolation. These biases can be minimized by comparing more realistic models, but a small systematic bias towards the choice of secondary contact models always remained. Using data from turbot populations from the Baltic Sea and North Sea, we further demonstrate using an empirical case study that not accounting for potential demographic changes in both ancestral and daughter populations can lead to overestimate divergence times and conclud that these populations diverged during a long allopatric phase (Le Moan *et al*., 2019) whereas our results indicate a very recent divergence with constant gene flow.

### Lessons from simulations

Testing basic models of divergence, such as AM, SC, IM and SI, relies on the assumption that model choice is largely robust to unmodelled demographic events. Surprisingly, until now this assumption was never formally tested, though it has been shown that in the original IM program (Hey and Nielsen, 2004) departures from assumptions can lead to strong biases in parameter estimation (Becquet and Przeworski, 2009). Using extensive simulations, we demonstrate that under a very broad range of divergence scenarios this assumption does not hold. Recent bottlenecks followed by growth in a daughter population always lead to strong support for SC models, regardless of *T*_*S*_. However, not accounting for a recent bottleneck can lead to overestimate or underestimate *T*_*S*_ depending on whether divergence is recent or older. This is because estimates of *T*_*S*_ tend to reflect the strength of the recent bottleneck, rather than divergence time. Another pattern emerging from our simulations is that failure to model a change in *N*_*e*_ in the ancestral population can lead to both biases in model choice and to overestimate divergence time, but this effect depends both on how recent divergence is and on how much time has passed between *T*_*AE*_ or *T*_*AB*_ and *T*_*S*_. Biases caused by unmodelled demographic changes in the ancestral population are most severe for recent divergence scenarios (i.e. when *T*_*S*_ « *T*_*AE*_ or *T*_*AB*_) and their effect on demographic inference fades when *T*_*S*_ ≈ 1. Similarly, the effect of a change in *N*_*e*_ in the ancestral population on parameter estimation and model choice becomes irrelevant when the last change in *N*_*e*_ occurred ≈ 4*N*_*e*_ generations before *T*_*S*_. This is theoretically expected, as it takes approximately 4*N*_*e*_ generations for a population to reach mutation-drift equilibrium (Kimura and Ohta, 1969; Lande, 1980).

The most commonly tested demographic models assume an ancestral population at mutation-drift equilibrium and a change in *N*_*e*_ is permitted only at the time *T*_*S*_. Therefore, an unmodelled ancestral expansion or contraction could push estimates of *T*_*S*_ back to the time of the ancestral change in *N*_*e*_. When *T*_*S*_ is small, demographic models that did not account for ancestral expansion tended to overestimate *T*_*S*_ by up to a factor of ten (as *T*_*AE*_ +*T*_*S*_ = 0.5, i.e. 10×*T*_*S*_). In our SC coalescent simulations *T*_*S*_ = 0.1 and *T*_*AE*_ +*T*_*S*_ = 0.5, and as expected, the overestimation of *T*_*S*_ was up to a factor of five. In IM simulations, unmodelled ancestral expansions also led to a bias towards SC models.

In our simulations of recent divergence scenarios under the SC model, an ancestral population expansion in some cases led to a slight bias towards choosing IM models. This is most likely because the effect of not modelling an ancestral expansion is to push *T*_*S*_ back in time, which under the scenario of long strict isolation will result in much longer divergence times. Roux *et al*. (2016) also demonstrated that when the period of strict isolation preceding SC is a small proportion (<60 %) of the total divergence time, distinguishing between IM and SC can be very difficult. When the true divergence scenario is a basic SC model, we obsrved that IM models that included very abrupt population size changes performed better than IM models that did not (but always worse than SC models). For example, IM_B_ fitted better than IM models, and IM_AEB_ models had even stronger support (Supplementary Figs 2-9).

All these observations taken together suggest that extreme caution should be exercised when choosing among competing divergence scenarios using methods based on the jAFS or its summary statistics. It is known that, if one allows competing models to be arbitrarily complex, there is an infinite number of demographic histories that can produce the same AFS (Myers *et al*., 2008). In reality, when comparing more biologically realistic models, a unique function producing the expected AFS is identifiable (Bhaskar and Song, 2014; Rosen *et al*., 2018). However, we demonstrate that when the models compared do not match closely the demographic history of the simulated populations, several demographic parameters in the model can yield better fits but lead to the wrong biological conclusion. Herein lies the major issue in interpreting results from demographic modelling: models are always extreme simplification of complex biological processes, and we often do not know what complexities can be safely excluded. Several studies (Ewing and Jensen, 2016; Pouyet *et al*., 2018; Roux *et al*., 2014, 2016) clearly showed that the effects of barrier loci and linked selection should be accounted for and here we demonstrate that historical changes in *N*_*e*_ cannot be ignored.

The systematic bias towards choice of SC models when the real scenario generating the data had continuous gene flow, and the general overestimation of the proportion of strict isolation for SC models, suggest that the use of simple IM and SC models to differentiate between primary and secondary divergence may often lead to the wrong conclusion. These findings have important repercussions on how we interpret demographic analyses of recent divergence scenarios; changes in *N*_*e*_ in ancestral populations have almost inevitably happened during past glacial cycles, and bottlenecks followed by population expansions are a classic signature of colonization of novel environments (e.g. Feng *et al*., 2020; Hewitt, 2008; Liu *et al*., 2016). Furthermore, as testing on a smaller number of simulations demonstrated, this bias is not unique to the main method employed in this manuscript (*moments*), but also applies to another very widely utilize approach to estimate demographic parameters based on the jAFS of multiple populations, *dadi* (Gutenkunst *et al*., 2009, Supplementay Fig. 18). It is unclear at this stage what the effect of unmodelled changes in *N*_*e*_ in ancestral and daughter populations would be on model choice and parameter estimation under an ABC framework. Most likely this will depend upon the choice of summary statistics. Most summary statistics commonly used (Watterson’s Θ, *π*, Tajima’s D, *F* _ST_ and *d*_*xy*_) are summaries of the AFS and therefore are also expected to be affected. However, statistics such as the decay of linkage disequilibrium are not, and could perhaps be less sensitive to these biases (e.g. Jay *et al*., 2019). It is interesting to notice that when reconstructing the demographic history of model organisms, it is common to test more realistic demographic scenarios modelling past demographic changes in ancestral and daughter populations as well as bottlenecks followed by population growth (eg: Garud *et al*., 2015; Gravel *et al*., 2011; Gutenkunst *et al*., 2009; Jouganous *et al*., 2017). However, these realistic demographic scenarios are more seldom considered when testing IM and SC models in non-model species. This possibly is in part due to the assumption that the data are inadequate to deal with such model complexity, and in part due to the assumption that model choice and parameter estimation is robust to such unmodelled demographic events. Here we showed that neither of these assumptions is correct. It should also be noted that a novel unsupervised approach for inferring demographic histories that performs jointly model structure and parameter optimization (GADMA Noskova *et al*.,2020) has the potential to alleviate the biases we describe here.

It is not our intention to suggest that most studies that compared IM and SC models and found strong support for SC have likely chosen the wrong scenario. Several very recent studies have indeed modelled the potential effects of bottlenecks (e.g. Christe *et al*., 2017; Hartmann *et al*., 2020; Montano *et al*., 2015; Rougemont *et al*., 2020; Rougeux *et al*., 2017, 2019), and formal model testing is often only one of several lines of evidence suggesting secondary contact (e.g. Le Moan *et al*., 2016; Rougemont *et al*., 2017; Roux *et al*., 2014; Tine *et al*., 2014). For example, a correlation between *F*_*ST*_ and *d*_*xy*_ (i.e. elevated divergence in genomic islands of differentiation) provides further evidence for SC (e.g. Cruickshank and Hahn, 2014; Duranton *et al*., 2018, 2020; Gagnaire *et al*., 2018; Nelson and Cresko, 2018). Nevertheless, the results clearly show that realistic demographic scenarios can generate strong false support for SC models with long periods of strict isolation. This has clear implication for studying incipient speciation and recent population divergence, for example as a result of range expansions and colonization of novel habitats following the end of the last glaciation.

It must also be considered that we explored a limited number of unmodelled demographic events. Spatial genetic structure, recent range expansions and admixture from ghost populations are all common demographic events, and these scenarios can also lead to biases in demographic inference (e.g. Delser *et al*., 2019). Similarly, while it is relatively simple to account (albeit in a coarse way) for linked background selection by modelling heterogeneity of *N*_*e*_ across the genome, it is more difficult to model the possible effects of a reduction in *N*_*e*_ in genomic regions of a single population (i.e. the expectation for linked positive selection). Since selective sweeps have the same local effect of a bottleneck (and can indeed lead to false inferences of changes in *N*_*e*_, e.g. Schrider *et al*., 2016), it is reasonable to assume that the presence of large recent selective sweeps may also lead to a bias towards SC models.

### Lessons from the turbot’s demographic history

A clear example of when the biases we describe in this study are especially problematic is given by the study of the origin of the Baltic Sea marine biodiversity. The Baltic Sea is a large body of brackish water which became connected to the North Sea about 8 kya. Its marine fauna has probably more than one origin, with evidence of populations and species in the Baltic Sea being the result of both primary and secondary divergence (reviewed in Johannesson *et al*., 2020). The evolutionary origin of some specific taxa, such as the Baltic Sea populations of Pleuronectiformes, remains controversial (Jokinen *et al*., 2019; Le Moan *et al*., 2019; Momigliano *et al*., 2017, 2018). In our empirical study our model fit suggests that *S. maximus* from the Baltic Sea originated from a very recent invasion (< 6 kya) from the North Sea and diverged with continuous gene flow. One-population models suggest that North Sea and Baltic Sea *S. maximus* share the same demographic history, with an ancestral population expansion that occurred 35-102 kya, until approximately 5 kya, roughly the divergence time estimated by our two-population models. After this, both stairway plots and two-population models show support for a founder event coincident with the time at which the Baltic Sea had the highest salinity in its history (Gustafsson and Westman, 2002). At this time, as noted by Momigliano *et al*. (2017), there would have been broad opportunity for marine fish to colonize the Baltic Sea. Failure to account for these realistic demographic events would have resulted in a very strong (*W*_*AIC*_ *>* 0.999) support for SC models and estimates of divergence times that predate the origin of the Baltic Sea.

Le Moan *et al*. (2019) reconstructed the demographic history of North Sea-Baltic Sea population pairs for five flatfish species (including *S. maximus*) using similar data and approaches as in this study, but assuming an ancestral population at equilibrium and no bottleneck followed by population growth associated with the invasion of the Baltic Sea. The authors found strong support for SC in four out of the five populations studied, with estimates of divergence time for each of these population pairs predating the origin of the Baltic Sea by at least a factor of five (Le Moan *et al*., 2019). Since their estimates of timing of secondary contact was different for the four species, the authors concluded that the population-pairs diverged in strict isolation in several unidentified marine refugia, and colonized the Baltic Sea at different times following the end of the last glaciation (Le Moan *et al*., 2019). As there is no other evidence for such scenario apart from testing of IM and SC demographic models, it is possible that Le Moan et al (Le Moan *et al*., 2019) results are a product of the biases we described here. We also note that when we do not assume an ancestral population at equilibrium and model potential bottlenecks, IM and SC models converge towards the same demographic scenario (Fig. 8) giving strong support for postglacial divergence between flatfish populations in the North Sea and the Baltic Sea. Furthermore, the likleihood ratio test provided no support for the secondary contact model. Interestingly, Momigliano *et al*. (2017) used ABC to model the divergence of the flounder species pair (*Platichthys flesus* and *P. solemdali*) in the Baltic Sea taking into account both potential changes in *N*_*e*_ in ancestral and daughter populations (but with no formal testing of IM and SC scenarios), finding support for postglacial colonization of these two flatfish species. Improved modelling in this study has led to support for a more biologically plausible history of *S. maximus*’s invasion of the Baltic Sea but we wish to caution that it is entirely possible that new data and/or better modelling approaches will provide in the future support for an alternative evolutionary scenario. We also wish to caution against over-interpreting scaled parameters, since their value is dependent on the mutation rate, the exact value of which is not know.

The genomic landscape of differentiation between North Sea and Baltic Sea turbot populations also points towards shallow divergence and recent selection in the Baltic Sea. Differentiation across the genome was generally low, and genomic islands of high allelic differentiation (*F*_*ST*_) were driven by strongly reduced *π* in the Baltic Sea rather than by a local increase in *d*_*xy*_, i.e. the classic signature of a recent selective sweep. Indeed within these genomic islands, *d*_*xy*_ approximates *π* in the North Sea; assuming the North Sea still represents ancestral population diversity, this suggests that net divergence among the selected haplotypes is close to 0. As noted by Cruickshank and Hahn (Cruickshank and Hahn, 2014), genomic islands of differentiation that evolved in allopatry and resist introgression (the expected pattern under a model of SC and heterogeneous gene flow) are expected to instead show increased levels of absolute divergence.

## Conclusion

In conclusion, using very extensive simulations as well as empirical data on turbots we demonstrate that testing IM and SC models can be difficult when the demographic history of the studied taxa deviates from the tested scenarios. We conclude that statistical support for SC or IM in model testing can often be an artifact of unmodelled demographic events, and that estimates of *T*_*S*_ can often reflect recent changes in *N*_*e*_, rather than divergence time. Given the centrality of formal testing between competing divergence scenarios in current research on local adaptation and speciation, these biases should not be ignored. Testing one-population models can provide guidance in identifying what demographic scenarios need to be incorporated in formal model testing, and testing more realistic demographic scenarios is paramount for avoiding at least the most severe biases described in this manuscript. However, even when testing more realistic divergence models, extreme caution should be exercised when interpreting results.

## Material and methods

### Analyses of simulated data

#### Simulations of older divergence scenarios

We tested the effect of recent bottlenecks as well as ancestral expansions and contractions on model choice and parameter estimation within divergence scenarios with *T*_*S*_ ranging from 4000 to 128 000 generations. To do this, we first generated several simulations under an Isolation with Migration model using the software *ms* (Hudson, 2002). All simulated scenarios within this study share a few common parameters. The simplest models represent a scenario where an ancestral population of size *N*_*ANC*_ splits at time *T*_*S*_ in two populations of size *N*_1_ (which is always fixed at 20 000 individuals, and is used as the reference population size *N*_*REF*_) and *N*_2_ (*N*_2_ = 0.25×*N*_1_, i.e. 5000 individuals). The migration rate *m* is, unless explicitly mentioned, symmetrical and set to four. The migration rate is given in units of *M* = 4*N*_*REF*_ *m*, where *M* is the fraction of each population which is made up of migrants at a given generation. We explored the effects of unmodelled demographic events across six divergence times, with *T*_*S*_ values raging from 0.05 to 1.6 in units of 4*N*_*REF*_ generations (i.e. 4000 to 128 000 generations) in log2 steps (i.e. *T*_*S*_ = 0.05, 0.1, 0.2, 0.4, 0.8, 1.6).

For each *T*_*S*_ we generated simulations including a bottleneck in population 2 (Fig. 1A). The time of the bottleneck remains constant at 0.05 (*T*_*B*_), as we aim to represent the effect of a recent bottleneck associated with fluctuations in *N*_*e*_ within the last glacial cycle. We simulated bottlenecks of different strengths, so that *N*_2_ at times *T*_*B*_ ranges from 1% to 64% in log2 steps of current *N*_2_, and following the bottleneck population 2 experiences an exponential growth that starts at time *T*_*B*_ and continues until present. This resulted in six simulations without bottlenecks (one for each *T*_*S*_) as well as 48 simulations including a bottleneck in population 2 (six *T*_*S*_ and eight strengths of bottleneck).

We further investigated the effects of changes in effective population size in the ancestral population, i.e. an ancestral expansion (AE) and an ancestral bottleneck (AB) (Fig. 1A). We included nine scenarios of population expansion at time *T*_*AE*_ + *T*_*S*_, and ancestral expansions were modelled as different values of ancestral population size *N*_*ANC*_ (to 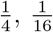 and 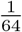 of *N*_*REF*_). We modelled every possible combination of six values of *T*_*S*_ (as above), three values for *T*_*AE*_ (*T*_*AE*_ = 0.025, 0.5, 1) and three strengths of ancestral expansion for a total of 54 independent simulations.

We simulated, in the same way, scenarios where the ancestral population underwent first a demographic contraction followed by expansion (Fig. 1 A). The time of the demographic contraction is set to *T*_*AB*_ + *T*_*S*_, where *T*_*AB*_ represent the number of generations (in units of 4*N*_*REF*_) before *T*_*S*_ at which the demographic contraction takes place. At time *T*_*S*_ +*T*_*AB*_*/*2 the ancestral population returns to its original size, which is equal to *N*_*REF*_. As per the AE scenario, we modelled every possible combination of six values of *T*_*S*_ (as above), three values fro *T*_*AB*_ (*T*_*AB*_ = 0.025, 0.5, 1) and three strengths of ancestral contraction for a total of 54 independent simulations.

For each of the simulations above, we simulated sampling of 20 individuals per populations and of 1 million unlinked loci of a length of 36 bp, which, when using a standard germ-line mutation rate (µ) of 1×10^−8^ and a *N*_*REF*_ of 20 000 individuals, results in roughly 80-120 thousand unlinked SNPs (depending on the specific model). We then used the unfolded jAFS from the simulated data for demographic inference.

#### Simulations of recent divergence scenarios

We furthermore tested more extensively the effects of unmodelled demographic events on model choice and parameter estimation on recent divergence scenarios (when *T*_*S*_ =0.05 and 0.1, i.e. 4000-8000 generations). These simulated scenarios are particularly relevant to the empirical case study we present later (the divergence of turbot populations in the North Sea and Baltic Sea). Here we simulated data under four gene-flow scenarios: SC and IM, each with symmetric (*M* = 4) and asymmetric gene flow (*m*_12_=4 and *m*_21_=16 where *M*_*ij*_ = 4*N*_*REF*_ *m*_*ij*_ and *M*_*ij*_ is the proportion of individuals in population *i* which is made up of migrants from population *j*). Modelling gene flow as the proportion of migrants in a given population (rather than the proportion of individuals migrating from a population), while not always realistic, has the advantage to keep gene flow constant even while *N*_*e*_ fluctuates among populations exchanging genes. For each of these gene flow scenarios, we modelled 64 combinations of demographic events in a fully orthogonal design (Fig. 1B). The simplest models represent IM and SC scenarios where an ancestral population of size *N*_*REF*_ splits at time *T*_*S*_ in two populations of size *N*_1_ and *N*_2_ (which have the same values as given above). We then included seven scenarios of population expansion at time *T*_*AE*_ (*T*_*SC*_ + *T*_*AE*_ = 0.5, in units of 4×*N*_*REF*_ generations, i.e. 40 000 generations), ranging from 2×*N*_*ANC*_ to 128×*N*_*ANC*_ on log2 steps. Since our reference population size (*N*_*REF*_) for parameter scaling is always *N*_1_, ancestral expansions were modelled as different values of *N*_*ANC*_ (to 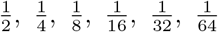 and 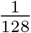 of *N*_1_). Furthermore, we modelled seven scenarios of bottlenecks followed by exponential growth in population two. In these scenarios *N*_2_ at time *T*_*S*_ can be 1%, 2%, 4%, 8%, 16%, 32%, and 64% of contemporary *N*_2_, reflecting a range of strong to very mild reduction in *N*_*e*_ at time of divergence. Following the bottleneck, exponential growth took place during a period lasting 2000 generations following *T*_*S*_, after which the *N*_*e*_ of population two reached *N*_2_. This kind of scenario is meant to reflect the invasion of a novel habitat, for example a new environment that became available after the end of the last glaciation. In IM models, the time of divergence is set at 4000 generations ago (*T*_*S*_ = 0.05×4*N*_*REF*_). In SC models, the time of divergence is set at 8000 generations ago (*T*_*S*_ = 0.1×4*N*_*REF*_), while secondary contact is established at time *T*_*SC*_ (*T*_*SC*_ = 0.25×*T*_*S*_), i.e. 2000 generations ago).

Fully orthogonal combinations of all the demographic scenarios outlined above (SC and IM, symmetric and asymmetric gene flow, ancestral expansions and bottlenecks) resulted in a total of 256 simulated scenarios. Firstly, for each of the 256 scenarios we simulated sampling of 20 individuals per populations and of 1 million unlinked loci of a length of 36 bp (as for the simulations above). We then estimated the folded and unfolded jAFS from the simulated data. These data sets represent standard data-sets when working with high quality whole genome data, assuming both scenarios whereby the genome of a closely related species is and is not available to polarize the jAFS. Secondly, for each of the 256 scenarios we simulated sampling of 10 individuals per populations and of 100 000 unlinked loci of a length of 36 bp, resulting in roughly 8-12 thousand unlinked SNPs (depending on the specific model) and estimated again both the unfolded and the folded jAFS. These data sets represent a standard, small scale 2b-RAD data set for a non-model species (i.e. the kind of data that most people working of non-model organisms can easily access). This led to a total of 512 coalescent simulations of recent divergence scenarios.

#### Demographic modelling of simulated data

Demographic modelling of simulated data was carried out using the software package *moments* (Jouganous *et al*., 2017), which is a development of the *dadi* (Gutenkunst *et al*., 2009) method for demographic inference from genetic data based on diffusion approximation of the allele frequency spectrum. *Moments* introduces a new simulation engine based on the direct computation of the jAFS using a model of ordinary differential equations for the evolution of allele frequencies that is closely related to the diffusion approximation used in *dadi* but avoids some of its limitations (Jouganous *et al*., 2017). Firstly, for all simulated scenarios, we tested whether a simple isolation with migration (IM) or a secondary contact (SC) model fitted the data best (IM and SC models in Fig. 1C). The models consist of an ancestral population of size *N*_*ANC*_ that splits into two populations of sizes *N*_1_ and *N*_2_ at time *T*_*S*_. In the IM model there is continuous asymmetric migration. In the SC model there is a period of strict isolation starting at time *T*_*S*_ followed by a period of secondary contact with asymmetric migration starting at time *T*_*SC*_.

For the 256 simulations of recent divergence scenarios (where biases were found to be most severe), we also tested models that accounted for population size changes in the ancestral population (AE models) and bottlenecks followed by growth in population two (B models), as well as both demographic changes (AEB) models (Fig. 1C). Therefore, for both IM and SC scenarios we had four alternative models: a basic scenario assuming an ancestral population at equilibrium and instantaneous size changes at time *T*_*S*_, and the three above mentioned combinations of demographic changes in the ancestral population and in population two (AE, B and AEB models). It is to be noted that models including bottlenecks intentionally do not match exactly the coalescent simulations (Fig. 1B), but rather mirror how growth is modelled in previously utilized models in speciation research (e.g. Rougeux *et al*., 2017). In IM_B_ models exponential growth starts at time *T*_*S*_ and continues until the present while in SC_B_ models exponential growth starts at time *T*_*SC*_ and continues until the present (Fig. 1 B).

Models were optimized for three rounds following an approach similar to Portik et al (Portik *et al*., 2017). In the first round, a Broyden-Fletcher-Goldfarb-Shanno (BFGS) algorithm optimization (function “optimize.log”; max-iter=10) was run for 10 sets of threefold randomly perturbed parameters. In the second round, the parameters from the replicate with the highest likelihood from each model were used as a starting point and the same optimization algorithm was used on 10 sets of twofold randomly perturbed parameters, increasing max-iter to 20. We repeated the same procedure for round 3, but using onefold perturbed parameters and setting max-iter to 30. We ran the entire optimization procedure 10 times to check for convergence among independent optimizations. We selected the replicate with the highest likelihood for each of the two-populations models and calculated the Akaike Information Criterion as −2(*log* −*likelihood*)+2*K* where *K* is the number of model parameters, and the Δ*AIC*_*i*_ = *AIC*_*i*_ −*AIC*_*min*_. We then calculated Akaike weight of evidence (*W*_*AIC*_) as outlined in Rougeux *et al*. (2017). The equation is outlined below, and *R* represents the total number of models compared.

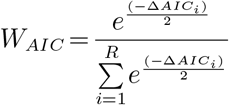

Firstly, we looked at the effect of unaccounted demographic events when our model choice was restricted to the basic IM and SC models (i.e. excluding models that account for demographic size changes in the ancestral population and a bottleneck and growth in population two). We evaluated not only if the IM and SC models were correctly identified, but also whether not accounting for ancestral expansion and bottlenecks affected parameter estimation.

Second, for all scenarios of recent divergence (i.e. the ones more relevant to our empirical study), we looked at whether including models with ancestral population expansions and bottlenecks improved model choice and parameter estimation. When comparing the eight full models, we calculated the *W*_*AIC*_ for each gene flow scenario (IM vs SC) as the sum of *W*_*AIC*_ for all IM and SC models. This comparison along family of models was carried out since at times the correct gene flow scenario was identified, but one of the demographic changes modelled was not (i.e. this process maximized our chance to recover the simulated gene flow scenario). For the simulations of ancestral expansions (AE) and ancestral bottlenecks (AB), we used only the unfolded jAFS. Since in our empirical study we use the folded jAFS, for all 512 simulations of recent divergence scenarios, model optimization and parameter estimation were carried out using both the folded and unfolded jAFS, to test whether demographic inference using the unpolarized jAFS is less or more susceptible to biases in model choice and parameter estimation.

#### Comparison between methods

To determine whether the biases reported in this study were specific to *moments* or reflected a general issue in inferring demographic histories from the jAFS, we repeated a portion of the analyses using the diffusion approximation approach implemented in *dadi*. We performed model choice and parameter estimation for the 126 simulations of recent divergence scenarios under the IM model (Fig. 1B), with symmetric and asymmetric gene flow and using only the simulations of the larger datasets (1 million loci). We used exactly the same optimization strategy, but only ran 3 independent optimization routines, which were sufficient to get convergence for independent runs of the simple IM and SC models.

### Analysis of empirical data

#### Sampling

We obtained a total of 172 samples of *Scophtalmus maximus* from seven locations and three biogeographic regions (Supplementary Table 2 and Fig. 5A): the North Sea (1 location, N=20), the transition zone separating the North Sea from the Baltic Sea (2 sampling locations, Vendelsö: N=27 and Öresund: N=35), and the Baltic Sea (4 populations, Dabki: N=24, Gdynia: N=23, Gotland: N=20 and Gotska Sandön: N=24). Individuals from the transition zone and the Baltic Sea are a subsample of the individuals analyzed by Florin and Höglund (2007), while samples from the North Sea were originally collected by Nielsen *et al*. (2004).

#### Library preparation

We built 2b-RAD libraries following the approach described by Wang *et al*. (2012), but with degenerate adaptors to allow identification of PCR duplicates. The protocol is described in detail by Momigliano *et al*. (2018). In short, DNA was extracted using a modified salting out protocol, and about 200 ng of DNA was digested with the type II b enzyme BcgI (New England Biolabs). This enzyme cuts both upstream and downstream of the 6 bp recognition site, creating fragments of a length of exactly 36 bp with 2 bp overhangs. Adaptors, one of which included degenerate bases, were ligated and the fragments amplified via 10 cycles of PCR as described in Momigliano *et al*. (2018). Fragments of the expected size were isolated using a BluePippin machine (Sage Science). Libraries were sequenced on Illumina machines (NextSeq500 and Hiseq 4000) to achieve a mean coverage of approximately 20x.

#### Bioinformatics and basic population genetic analyses

Raw reads were demultiplexed and PCR duplicates were removed as per Momigliano *et al*. (2018), then mapped to the latest version of *S. maximus* reference genome (Figueras *et al*., 2016, Assembly ASM318616v1, GenBank accession: GCA003186165) using Bowtie2 (Langmead and Salzberg, 2012). SAM files were converted to BAM files and indexed using SAMTOOLS (Li *et al*., 2009). A genotype likelihood file in beagle format was produced in the software ANGSD (Korneliussen *et al*., 2014), using the following filters: no more than 20% missing data, retaining only biallelic loci, removing bases with mapping quality below 30 and Phred quality scores below 20. See Supplementary Fig. 1 for summary statistics. A principal component analysis (PCA) based on genotype likelihoods was performed using the software PCAngds (Meisner and Albrechtsen, 2018) using only variants with a minor allele frequency above 0.02. The folded AFS for each population as well as the jAFS were produced in ANGSD. ANGSD calculates folded jAFSs where the minor allele is computed separately for each AFS while *moments* expects minor alleles to be estimate for the jAFS. Thus, for the jAFS for NS and BS, we produced the unfolded jAFS in ANGSD in the form of a *dadi* data dictionary. We then selected a random SNP within each locus and folded the jAFS in *moments*.

We produced also a VCF (Variant Call File) using the UnifiedGenotyper function from GATK v.3.8. Following UnifiedGenotyper, we removed individuals with an average read depth below seven. Then, we used four technical replicate pairs (i.e. four pairs of individuals for which we constructed and sequenced two independent libraries) to generate a list of SNPs for which we have high confidence (i.e. which show 100% matches between all replicate pairs). We used this list of SNPs to carry out Variant Quality Score Recalibration (VQSR), following GATK best practice (Dixon *et al*., 2015). Finally, we removed genotype calls with low sequencing depth (<7), removed indels, triallelic SNPs, SNPs with minor allele frequencies below 0.01 and with more than 10% missing data. This resulted in a final VCF containing 12678 sites genotyped for 154 individuals. This data-set was used to calculate Weir and Cockerham *F*_*ST*_ for each SNP. Furthermore, we thinned the data retaining, for each tag, the SNP with the highest minor allele frequency and used this data for inferring population structure using fastSTRUCTURE (Raj *et al*., 2014). Bioinformatic steps, scripts for analyses, and the jAFSs used are publicly available (see Data Availability section).

#### Inferring the genomic landscape of differentiation

We used several approaches to identify potential islands of differentiation across the genome between North Sea and Baltic Sea turbots. Firstly, we calculated from the called genotypes *F*_*ST*_ between the North Sea and Baltic (excluding individuals from the transition zone) using VCFTOOLS (Danecek *et al*., 2011). This is the only measure of differentiation we calculated from called genotypes. Secondly, we used the software package PCAngsd to run a selection scan using an extended model of FastPCA (Galinsky *et al*., 2016) working directly on genotype likelihoods, based on the input beagle file we used for the PCA. This approach identifies variants whose differentiation along a specific principal component (in our case the first PC) is greater than the null distribution under genetic drift. To account for multiple comparisons, we converted p-values to q-values (False Discovery Rate, FDR) following Benjamin and Hochberg (Benjamini and Hochberg, 1995). As a second approach to classify SNPs as outliers,we used a Hidden Markov model (HMM) approach to detect genomic islands, based on the uncorrected p-values from PCAngsd selection scan (Hofer *et al*., 2012; Marques *et al*., 2016). We counted as candidate outliers SNPs that show and FDR < 0.1 and that simultaneously were identified as outliers by the HMM test. If adjacent SNPs identified by both approaches lied within a distance of < 500kb, we identified as part of the same genomic island of differentiation.

We then obtained estimates of within population genetic diversity (π) and absolute divergence (*d*_*xy*_) across the genome. In order to maximize the usable data and account for differences in coverage among samples we performed all analyses in ANGSD directely from genotype likelihoods. Firstly, we generated windows of 250kb across the genome using BedTOOLS (Quinlan and Hall, 2010). Then, we calculated the unfolded AFS (for the North Sea and Baltic Sea Individuals) as well as the jAFS for each 250kb window across the genome with ANGSD, using only filters that do not distort the AFS (-uniqueOnly 1 -remove_bads 1 -minMapQ 20 -minQ 20 -C 50). We finally used custom R scripts to calculate π, and (*d*_*xy*_) for each window, retaining only windows for which the AFS was derived from at at least 1000 sequenced bases. All scripts to calculate the AFS in windows and derive summary statistics are available from GitHub (sea Data Availability section). Note that for diversity analyses we used the unfolded AFS even if we do not have an appropriate outgroup, assuming the reference allele as the ancestral state. However, this is not an issue since the summary statistics calculated are based on allele frequencies, which are symmetrical with respect to folding.

#### Demographic modelling of empirical data

The demographic history of the North Sea and Baltic Sea populations of *S. maximus* were reconstructed using several approaches based on the one-dimensional (for one-population models) and joint folded allele frequency spectra (henceforth 1d-AFS and jAFS, respectively). Since taking into account possible changes in effective population size may have important effects on the estimation of parameters such as migration rate and divergence times (Gravel *et al*., 2011), we first determined the demographic history of each population independently using the 1d-AFS, using both *moments* and Stairway plots (Liu and Fu, 2015). We then proceeded to compare 32 two-population models to determine the demographic history from the jAFS. We used a µ of 1×10^−8^ and a generation time of 3.5 years for scaling demographic events to make results directly comparable to Le Moan et al (Le Moan *et al*., 2019). All models tested in this study are available from GitHub (sea Data Availability section).

#### One-population models

Firstly, we estimated past demographic changes in the North Sea populations and from the Baltic Sea populations (i.e. excluding samples from the transition zone) using the multi-epoch model implemented in the software Stairway plot v2 (Liu and Fu, 2015). Stairway plots use composite likelihood estimations of Θ at different epochs, which is then scaled using the mutation rate. For Baltic Sea populations, we estimated past demographic changes from the 1d-SFS from each sampling location independently. Stairway plots were generated including singletons, using 2/3 of the sites for training and four numbers of random break points for each try (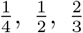, and 1 times the number of samples in each population). Since demographic histories were similar in all locations, and there was no evidence of population structure from other analyses, all subsequent analyses were performed using the jAFS estimated from pooling all samples from populations in the Baltic Sea.

Secondly, we compared three simple 1-population models using *moments*: a Standard Neutral Model (SNM, assuming constant population size at equilibrium), a 2 Epochs model (2EP, assuming a sudden population change at time T1) and a 3 Epochs model including a sudden demographic change at time *T*_*AE*_ followed by a bottleneck at time *T*_*B*_ followed by exponential growth (2EP_B_). The 1EP model represents a single demographic change and a scenario where a demographic expansion/contraction occurred either in the ancestral population from which the NS and BS populations are derived or in the NS and BS populations themselves. The 2EP_B_ model represent a scenario where, in addition to an ancestral expansion/contraction, there was a recent bottleneck followed by growth; this could be a realistic scenario for the Baltic Sea populations, which must have invaded the Baltic Sea following its connection to the North Sea within the past 8 ky.

#### Two-population models

Given the existence of a well-known hybrid-zone between the Baltic Sea and the North Sea (Nielsen *et al*., 2004), we tested the two main divergence scenarios that include contemporary migration: the isolation with migration model (IM), and the secondary contact (SC) models. All models consist of an ancestral population of size *N*_*ANC*_ that splits into two populations of size *N*_1_ and *N*_2_ at time *T*_*S*_. Migration is continuous and asymmetric in the IM model. The SC model includes a period of isolation starting at time *T*_*S*_ and a period of secondary contact when asymmetric migration starts at time *T*_*SC*_. For each of these basic models, we tested models that included heterogeneous migration rates across the genome (2M, i.e. islands resisting migration), and heterogeneous *N*_*e*_ across the genome (2N, a way to model linked selection) as described in Rougeux et al (Rougeux *et al*., 2017). We therefore had four possible variations of each of the basic model (for example: SC, SC_2M_, SC_2N_, SC_2M2N_), for a total of eight basic divergence models.

Both Stairway Plots and *moments* analyses of the empirical data suggest (see Results) a demographic expansion 20-100 kya, and that the BS may also have undergone a recent bottleneck. If such demographic changes had happened at the time of split between the two populations, this would be well captured by the eight models described above which take into account a single change in *N*_*e*_ from *N*_*ANC*_ to *N*_1_ and *N*_2_ at time *T*_*S*_. However, given that the timing and magnitude of the population expansion are very similar in all populations, another possibility is that the ancestral population underwent a demographic expansion prior to the split between the North Sea and the Baltic Sea. We incorporated this hypothesis by extending the eight models described above to include an ancestral population expansion (AE, ancestral expansion models), a recent bottleneck followed by population growth in the Baltic Sea (B, bottleneck models) or both (AEB models). In the AE models, the ancestral population undergoes a demographic change at time *T*_*AE*_, after which population size remain constant until time of split (*T*_*S*_). In the B models, the Baltic Sea population undergoes a bottleneck followed by population growth at time *T*_*S*_. This scenario mimics a possible invasion of the Baltic Sea from a small founder population. For both IM and SC models, all possible combinations of heterogeneous *N*_*e*_ (2N models), heterogeneous migration rates (2M models), ancestral expansion (AE models) and bottlenecks (B models) were tested, yielding 16 variations of the IM and SC models and a total of 32 models tested. It should be note that in 2N2M models, it is assumed that regions experiencing lower migration rates and regions experiencing lower effective population size do not overlap. This made convergence of the complex models easier, and it should not be problematic assuming the proportion of the genome experiencing lower migration rates and with lower *N*_*e*_ are relatively small (as is our case, see Supplementary Table 3).

#### Model optimization and model selection

With the exception of the SNM models, which has no free parameters, all one- and two-population models were optimized in five independent optimization routines, each consisting of five rounds of optimization using an approach similar to Portik et al (Portik *et al*., 2017). In the first round we generated 30 sets of threefold randomly perturbed parameters and for each ran a BFGS optimization setting max-iter to 30. In the second round, we chose for each model the replicate with the highest likelihood from the first round and generated 20 sets of threefold randomly perturbed parameters, followed by the same optimization strategy. We repeated the same procedure for round three, four and five, but using two-fold (round three) and one-fold (round four and five) perturbed parameters, respectively. In the final round, we also estimated 95% confidence intervals (CI) of the estimated parameters using the Fisher Information Matrix, as described by Coffman *et al*. (2015), and 95% CI of *T*_*S*_ and *T*_*SC*_ were estimated according to the rules of propagation of uncertainty. We selected the best replicate for each of the final models (three one-population models and 32 two-population models). We then calculated *W*_*AIC*_ as a relative measure of support for each model. Parameters in coalescent units were scaled based on estimates of *N*_*ANC*_ as outlined in *dadi* ‘s manual. *N*_*ANC*_ was calculated as Θ*/*(4*Lµ*), where *L* is the total sequence length from which SNPs used in demographic analyses originated and *µ* = 1×10^−8^. The total sequence length was calculated as *S* ×(*V*_*U*_ */V*_*T*_), where *S* is the number of sites (variant and invariant) retained in ANGSD to calculate the jAFS, *V*_*U*_ is the number of unlinked SNPs retained for demographic modelling and *V*_*T*_ is total number of variants in the jAFS before linked SNPs were removed. Time parameters were scaled assuming a generation time of 3.5 years to be directly comparable with a previous study on the same species (Le Moan *et al*., 2019).

The use of AIC to rank models and of the Fisher Information Matrix to estimate parameter uncertainty relies on the assumption that genetic data are independent. For RAD data, it is generally assumed that keeping one SNP per RAD locus is sufficient to satisfy this assumption. Nevertheless, we carried out further analyses to ensure that unaccounted linkage did not lead to *a)* biased estimates of parameter uncertainty and *b)* favoring more complex models. Firstly, we estimated parameter uncertainty for the two best models (that according to *W*_*AIC*_ had similar support) using block-bootstraps and the Godambe Information Matrix (GIM). Secondly, after the two best fitting models had been identified (in this case, IM_AEB2N_ and SC_AEB2N_) we used Likelihood Ratio Tests (LRT) to test for the support of specific parameters among nested models, controlling for Type I error by normalizing the difference in log-likelihoods as outilined in Coffman *et al*. (2015). We used LRT to test support for a SC scenario by comparing IM_AEB2N_ and SC_AEB2N_ models. Since the test failed to reject the simpler IM scenario, we further used LRT to test for support for a past bottleneck in the Baltic Sea population (comparing IM_2N_ and IM_B2N_ models) and for an ancestral expansion (comparing IM_2N_ and IM_AE2N_ models). The correct weighting of *χ*^2^ distributions for the LRT were calculated according to Ota *et al*. (2000).

## Aknowledgements

We thank Roger Butlin, Ryan Gutenkunst, Paul Blischak, Xin Huang, Petri Kemppainen, Bohao Fang, Pierre Nouhaud and Sean Stankowski for providing feedback on the manuscript. We thank Einar Nielsen for donating turbot samples from the North Sea. This study was funded by the Academy of Finland (grant 316294 to PM and 218343 to JM) and the Helsinki Institute of Life Science (HiLIFE).

## Author Contributions

PM conceived the study, performed laboratory work, analyzed the data and wrote the manuscript. JM contributed to study design, provided constructive feedback on the manuscript and funded sequencing. ABF contributed most of the samples and provided constructive feedback on the manuscript.

## Data Availability

All raw sequence data are publicly accessible via GenBank Sequence Read Archive (Bioproject ID PRJNA699405). Scripts and processed data are available at Zenodo data repository (doi: 10.5281/zenodo.4518372), and models for demographic inference are also available via GitHub (https://github.com/Nopaoli/ Demographic-Modelling).

## Notes

### Competing Interest Statement

The authors have declared no competing interest.

