## Supplementary Materials for "Biases in demographic modelling affect our understanding of recent divergence"

### Supplementary Information

**Supplementary Table 1**: Description of all models used to infer the demographic history from simulated and empirical data. When reconstructing the demographic history from empirical data, we also tested the same two-population demographic scenarios accounting for heterogeneous migration rates (2M), heterogeneous effective population size (2N) and both (2N2M).

| Graphic representation | Demographic scenarios | Number of populations | Modifiers | Number of free parameters |
|---|---|---|---|---|
| 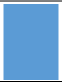 | SNM<br>Standard Neutral Model | 1 | None | 0 |
| 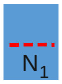 | 2EP<br>Two epochs model, one instantaneous change in $N_e$ | 1 | None | 2 |
| 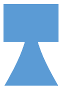 | 2EP$_B$<br>Two epochs model, one instantaneous change in $N_e$ followed by growth | 1 | None | 3 |
| 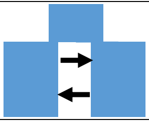 | IM<br>Isolation with Migration | 2 | 2M, 2N, 2M2N | Basic = 5<br>2N = 7<br>2M = 8<br>2M2N = 10 |
| 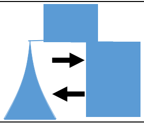 | IM$_B$<br>Isolation with Migration. Bottleneck and growth in one population | 2 | 2M, 2N, 2M2N | Basic = 6<br>2N = 8<br>2M = 9<br>2M2N = 11 |
| 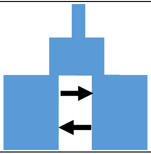 | IM$_{AE}$<br>Isolation with Migration. One change in $N_e$ in the ancestral population | 2 | 2M, 2N, 2M2N | Basic= 7<br>2N = 9<br>2M = 10<br>2M2N = 13 |
| 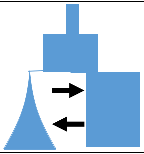 | IM$_{AEB}$<br>Isolation with Migration. One change in $N_e$ in the ancestral population, bottleneck and growth in a daughter population | 2 | 2M, 2N, 2M2N | Basic = 8<br>2N = 10<br>2M = 11<br>2M2N = 13 |
| 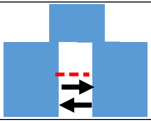 | SC<br>Secondary Contact | 2 | 2M, 2N, 2M2N | Basic = 6<br>2N = 8<br>2M = 9<br>2M2N = 11 |
| 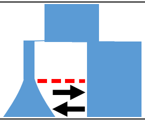 | SC<br>Secondary Contact. Bottleneck and growth in one population | 2 | 2M, 2N, 2M2N | Basic = 7<br>2N = 9<br>2M = 10<br>2M2N = 12 |
| 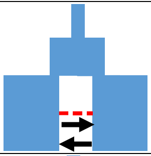 | SC$_{AE}$<br>Secondary Contact. One change in $N_e$ in the ancestral population | 2 | 2M, 2N, 2M2N | Basic = 8<br>2N = 10<br>2M = 11<br>2M2N = 13 |
| 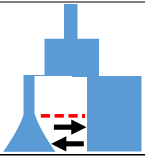 | SC$_{AEB}$<br>Secondary Contact. One change in $N_e$ in the ancestral population, bottleneck and growth in a daughter population | 2 | 2M, 2N, 2M2N | Basic = 9<br>2N = 11<br>2M = 12<br>2M2N = 14 |

**Supplementary Table 2**: Location names, population codes, sample size (N) and geographic coordinates of all sampling locations of *S. maxixmus* used in this study.

| Location | Population code | N | Latitude | Longitude | Reference |
|---|---|---|---|---|---|
| North Sea | NS | 20 | 56.58° N | 06.12° E | Nielsen et al. (2004) |
| Vendelsö | VD | 27 | 57.17° N | 12.06° E | Florin et al. (2007) |
| Öresund | OS | 35 | 55.34° N | 12.51° E | Florin et al. (2007) |
| Dabki | DAB | 24 | 54.45° N | 6.30° E | Florin et al. (2007) |
| Gdynia | GDY | 23 | 54.45° N | 8.30° E | Florin et al. (2007) |
| Gotland | GOT | 20 | 57.32° N | 18.57° E | Florin et al. (2007) |
| Gotska Station | GS | 24 | 58.22 °N | 19.15° E | Florin et al. (2007) |

**Supplementary Table 3**: Parameter estimates of the two best models for the empirical study, $\pm$sd inferred using the Fisher Information Matrix /FIM) and the Godambe Information Matrix (FIM), the latter in brackets. AIC= Akaike Information Criterion, $W_{AIC}$= weight of evidence, $\Theta = 4N_{REF}\mu$ , $N_{AE}$= $N_e$ after population expansion, $N_1$= contemporary $N_e$ in the North Sea, $N_2$= contemporary $N_e$ in the Baltic Sea, s=$N_e$ of the Baltic Sea at time of invasion as a proportion of $N_1$, $T_{AE}$= time of ancestral expansion, $T_S$ = time of population divergence, $T_{SC}$= time of secondary contact, $m_{12}$ = migration rates from the Baltic Sea to the North Sea, $m_{21}$= migration rates from the North Sea to the Baltic Sea, Bf= background factor, i.e. $N_e$ of the part of the genome with reduced recombination/higher background selection as a proportion of $N_e$ in the rest of the genome, Q= proportion of the genome with reduced $N_e$. Population sizes are given in units relative to $N_{REF}$. Times are given in units of $2N_{REF}$ generations, and need to be added backward in time (the real time of ancestral expansion for the $IM_{AEB2n}$ model is $T_{AE}$ + $T_S$). Migration rates are given in units $M_{ij} = 2N_{REFmij}$, where $M_{ij}$ is the proportion of individuals in population $i$ that is made up of migrants from population $j$ at a given generation. For FIM we used the pruned SNPs dataset (1 SNP per 2b-RAD locus), for estimating uncertainties with the GIM we used all SNPs. Therefore, we have two $\Theta$ estimates (the one for the data set including all SNPs is in brackets, with its standard deviation estimated from the GIM.)

| Model | *AIC* | $W_{AIC}$ | $\Theta$ | $N_{AE}$ | $N_1$ | $N_2$ | *s* | $T_{AE}$ | $T_S$ | $T_{SC}$ | $m_{12}$ | $m_{21}$ | *Bf* | *Q* |
|---|---|---|---|---|---|---|---|---|---|---|---|---|---|---|
| $IM_{AEB2N}$ | 5536 | 0.66 | 2398 ±104 (2682 ±430) | 1.68 ±0.069 (0.15) | 2.78 ±0.56 (1.07) | 2.77 ±0.73 (0.83) | 0.0356 ±0.006 (0.004) | 0.37 ±0.076 (0.065) | 0.02 ±0.003 (0.004) | NA | 4.66 ±2.35 (4.1) | 28.15 ±2.01 (4.04) | 0.1 ±0.013 (0.05) | 0.22 ±0.03 (0.06) |
| $SC_{AEB2N}$ | 5537.5 | 0.31 | 2383 ±105 (2666 ±409) | 1.675 ±0.07 (0.127) | 2.898 ±0.62 (1.07) | 2.75 ±0.7 (0.44) | 0.054 ±0.023 (0.007) | 0.38 ±0.08 (0.069) | 0.003 ±0.003 (0.002) | 0.017 ±0.0037 (0.0027) | 4.53 ±2.54 (3.22) | 28.15 ±2.26 (1.84) | 0.1 ±0.012 (0.049 | 0.22 ±0.03 (0.06) |

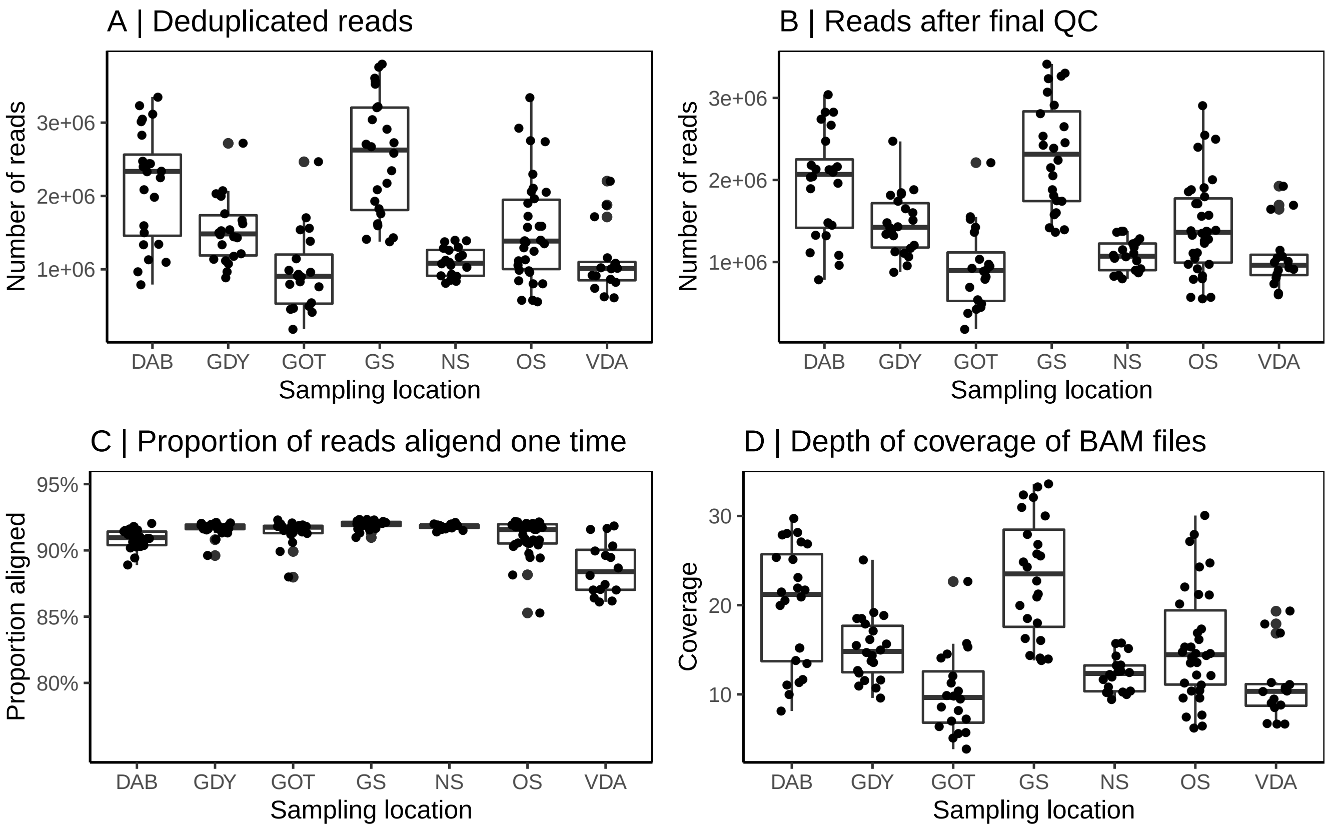


**Supplementary Fig. 1:** summary statistics from 2b-RAD data. Number of deduplicated reads (**A**) reads after quality check (**B**), proportion of reads which mapped one time to the reference genome (**C**) and mean per individual depth of coverage calculated from BAM files (**D**).

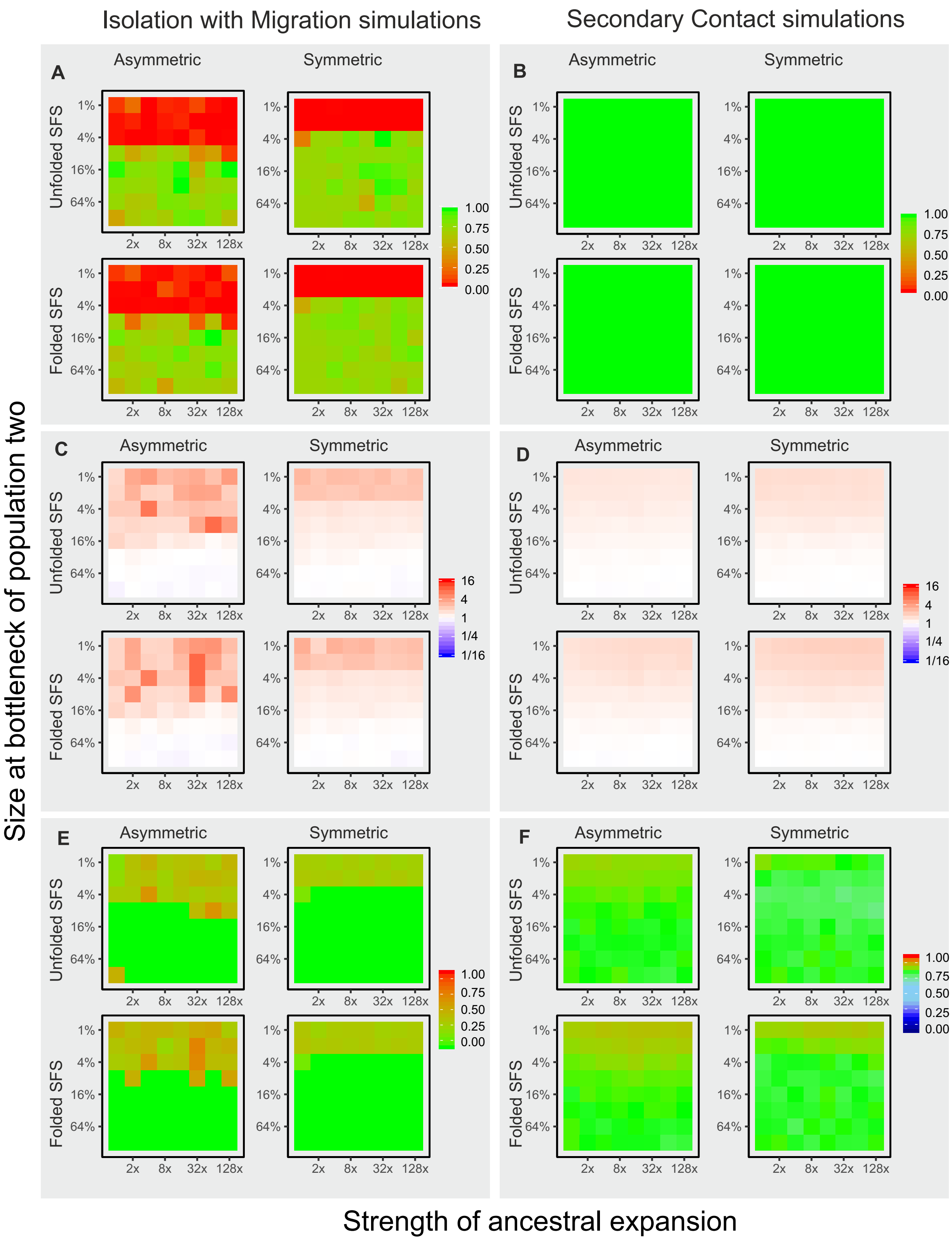


**Supplementary Fig. 2:** Model choice and parameter misestimation for all eight divergence models for all simulations of the larger data sets (1 million loci). Left panels **A, C & C** show results from simulations with constant migration (IM), right panels **B, D & F** show results from simulation with a period of strict isolation (SC). Within each panel, results are shown for simulations with symmetric and asymmetric migration, and for model estimates using the folded or unfolded jAFS. Within each panel, each graph represent the values for all 64 simulations as per Fig. 1. Panel **A** and **B** show weight of evidence for the correct gene flow scenario (0-1). **C & D** show misestemation of the parameter $T_S$ ($T_S$ estimate /$T_S$ of simulation). Panels **E** and **F** show the estimated proportion of the divergence time for which the model inferred strict isolation (green represent the correct time, i.e. 0 for IM model and 0.75 for SC models).

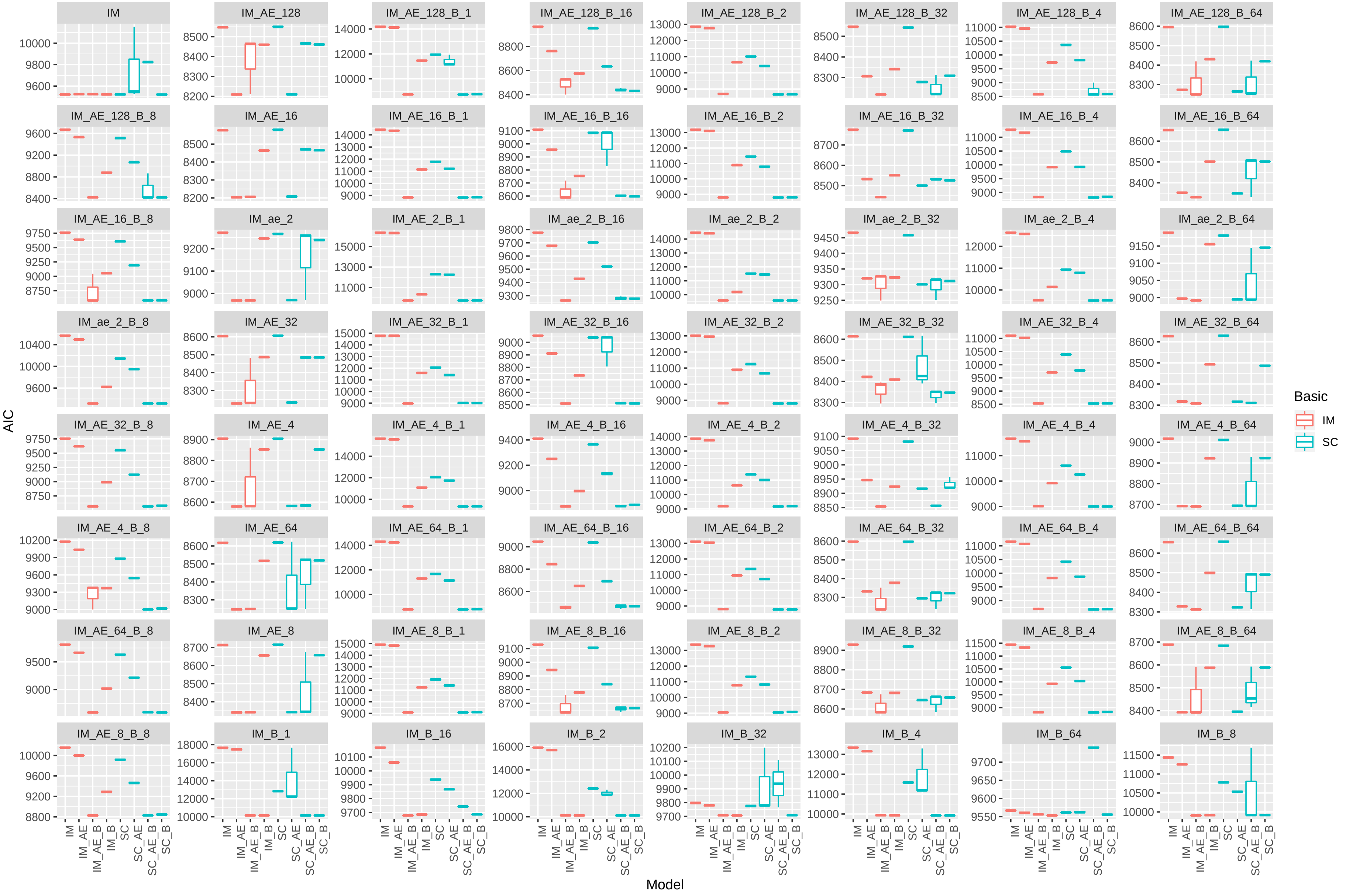


**Supplementary Fig. 3:** Boxplot showing AIC estimates for IM and SC models for the unfolded jAFS of 64 simulations (1 million loci) under the asymmetric IM model. The best three replicates for each model/simulation combination are plotted. Note the very narrow boxplots showing models have converged to the same likelihhods, with a few exception.

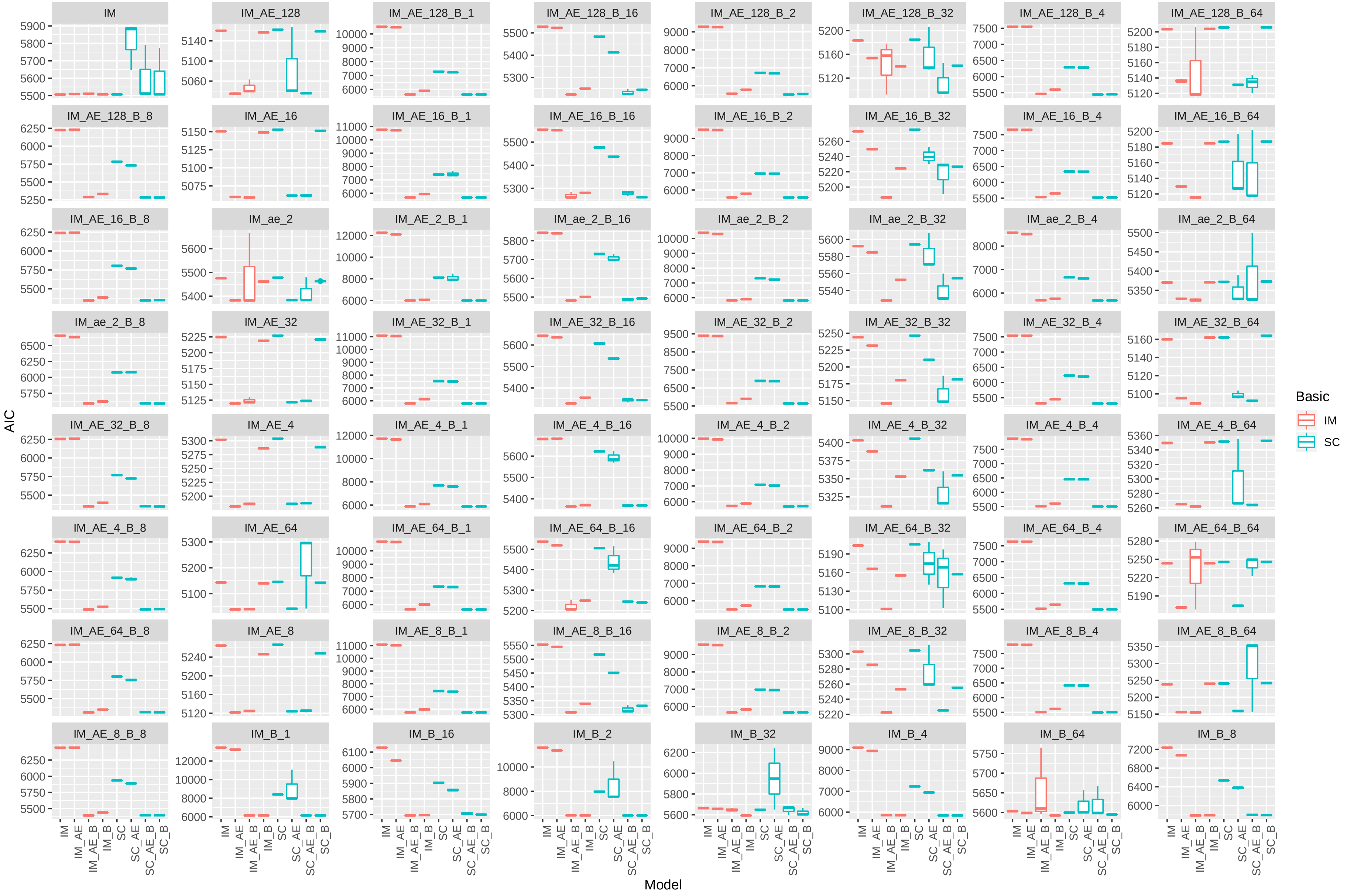


**Supplementary Fig. 4:** Boxplot showing AIC estimates for IM and SC models for the folded jAFS of 64 simulations (1 million loci) under the asymmetric IM model. The best three replicates for each model/simulation combination are plotted. Note the very narrow boxplots showing models have converged to the same likelihhods, with a few exception.

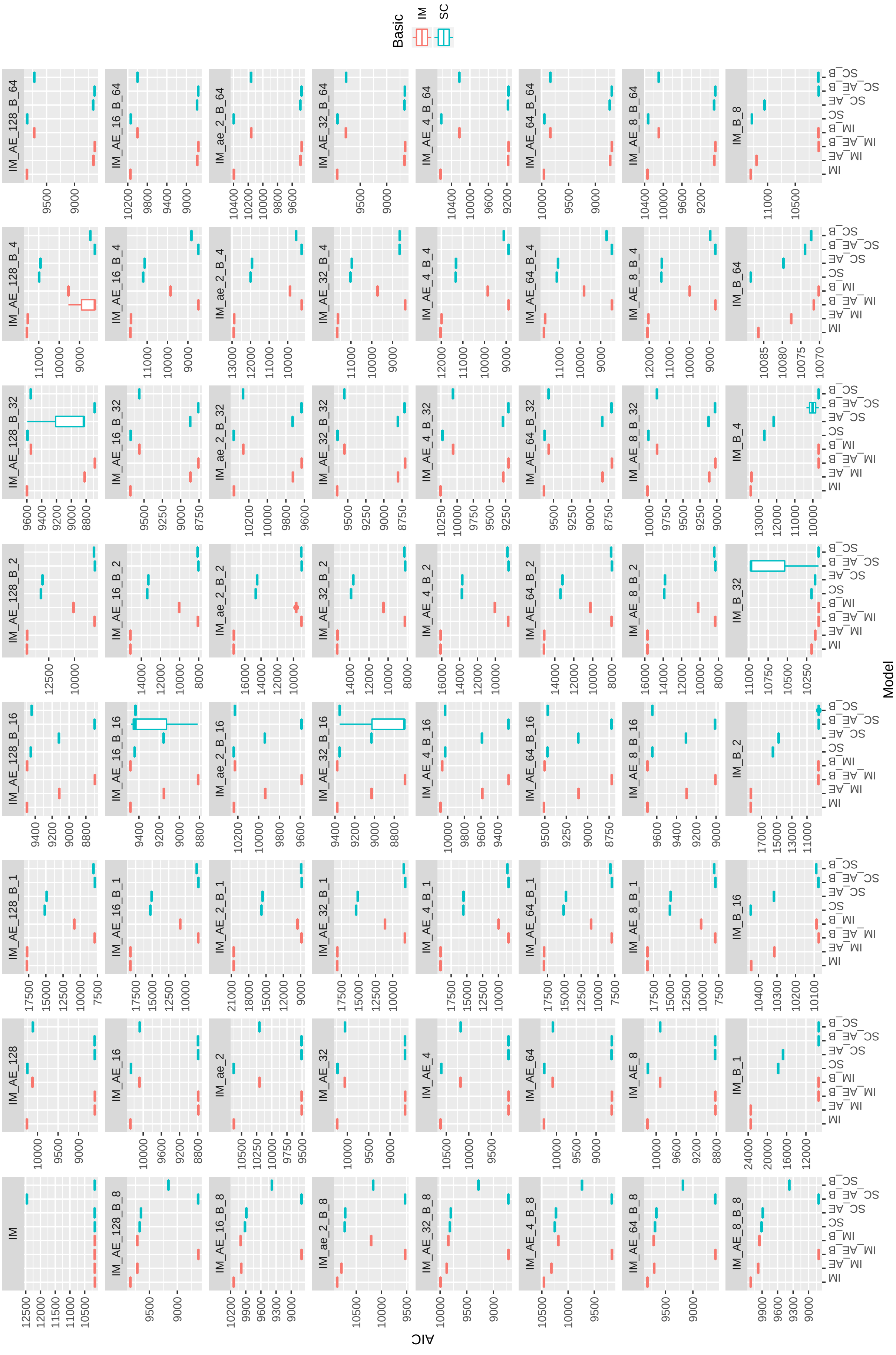


**Supplementary Fig. 5:** Boxplot showing AIC estimates for IM and SC models for the unfolded jAFS of 64 simulations (1 million loci) under the symmetric IM model. The best three replicates for each model/simulation combination are plotted. Note the very narrow boxplots showing models have converged to the same likelihoods, with a few exception.

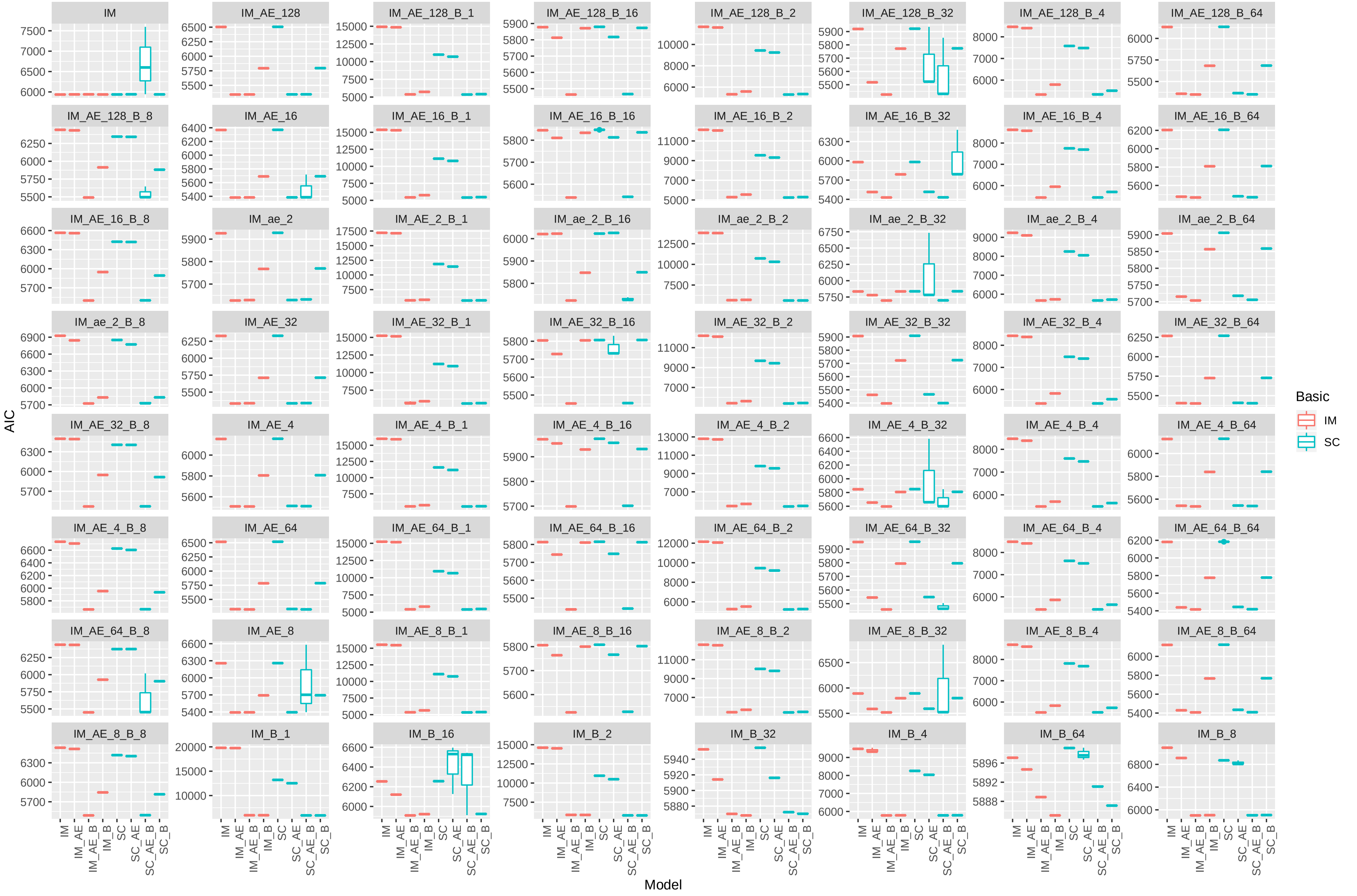


**Supplementary Fig. 6:** Boxplot showing AIC estimates for IM and SC models for the folded jAFS of 64 simulations (1 million loci) under the symmetric IM model. The best three replicates for each model/simulation combination are plotted. Note the very narrow boxplots showing models have converged to the same likelihhods, with a few exception.

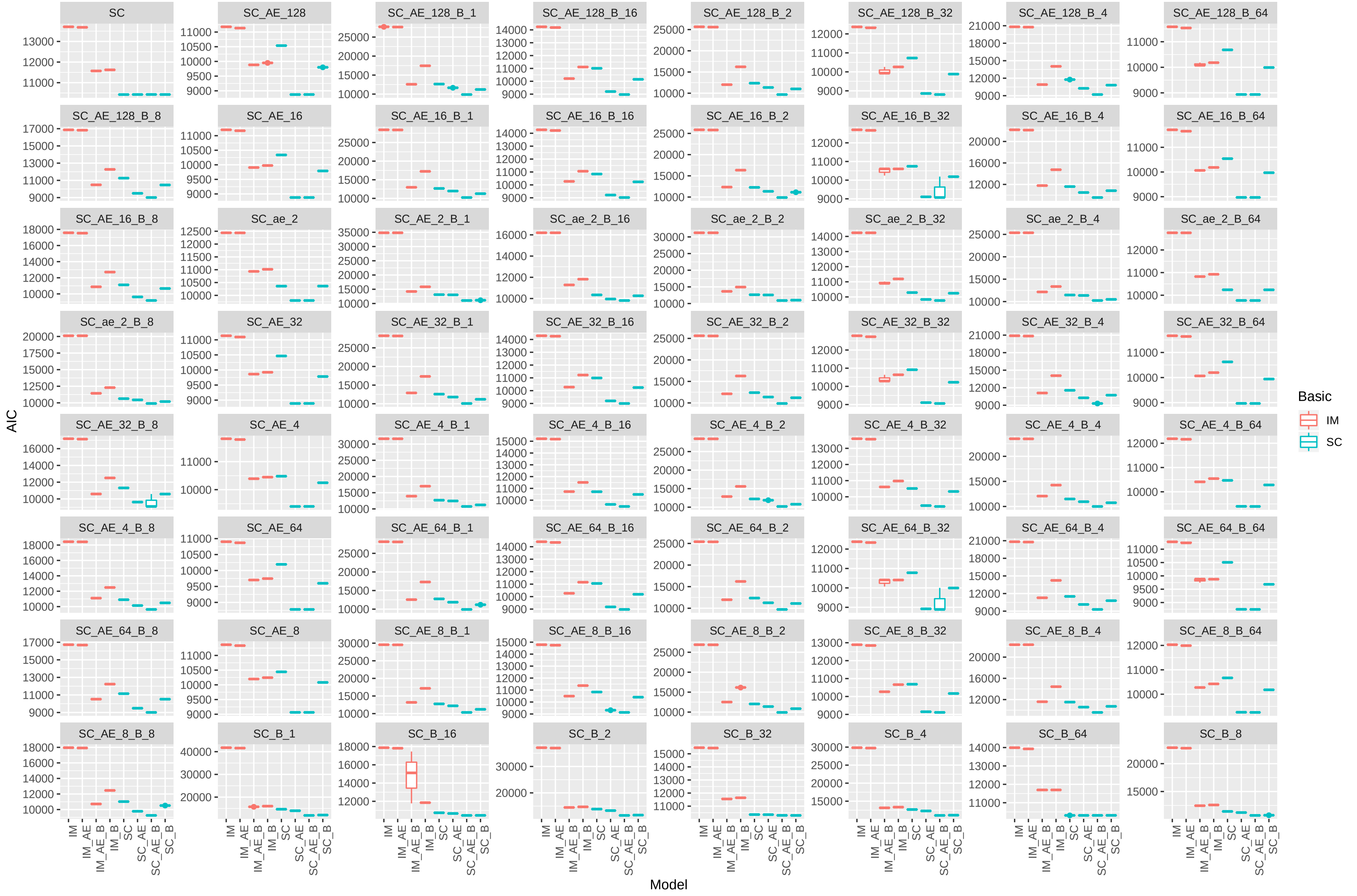

**Supplementary Fig. 7:** Boxplot showing AIC estimates for IM and SC models for the unfolded jAFS of 64 simulations (1 million loci) under the asymmetric SC model. The best three replicates for each model/simulation combination are plotted. Note the very narrow boxplots showing models have converged to the same likelihhods, with a few exception.

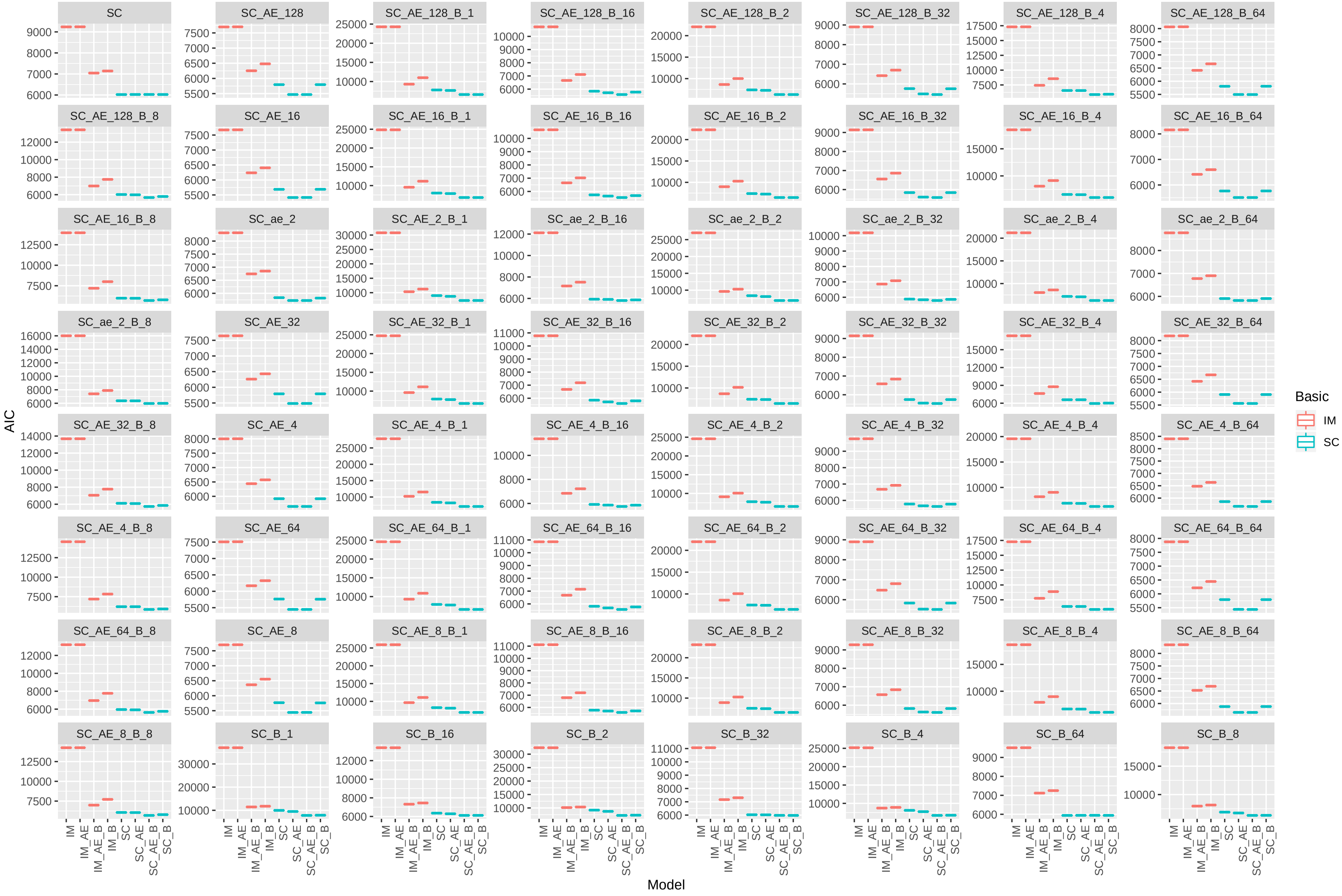


**Supplementary Fig. 8:** Boxplot showing AIC estimates for IM and SC models for the folded jAFS of 64 simulations (1 million loci) under the asymmetric SC model. The best three replicates for each model/simulation combination are plotted. Note the very narrow boxplots showing models have converged to the same likelihoods, with a few exception.

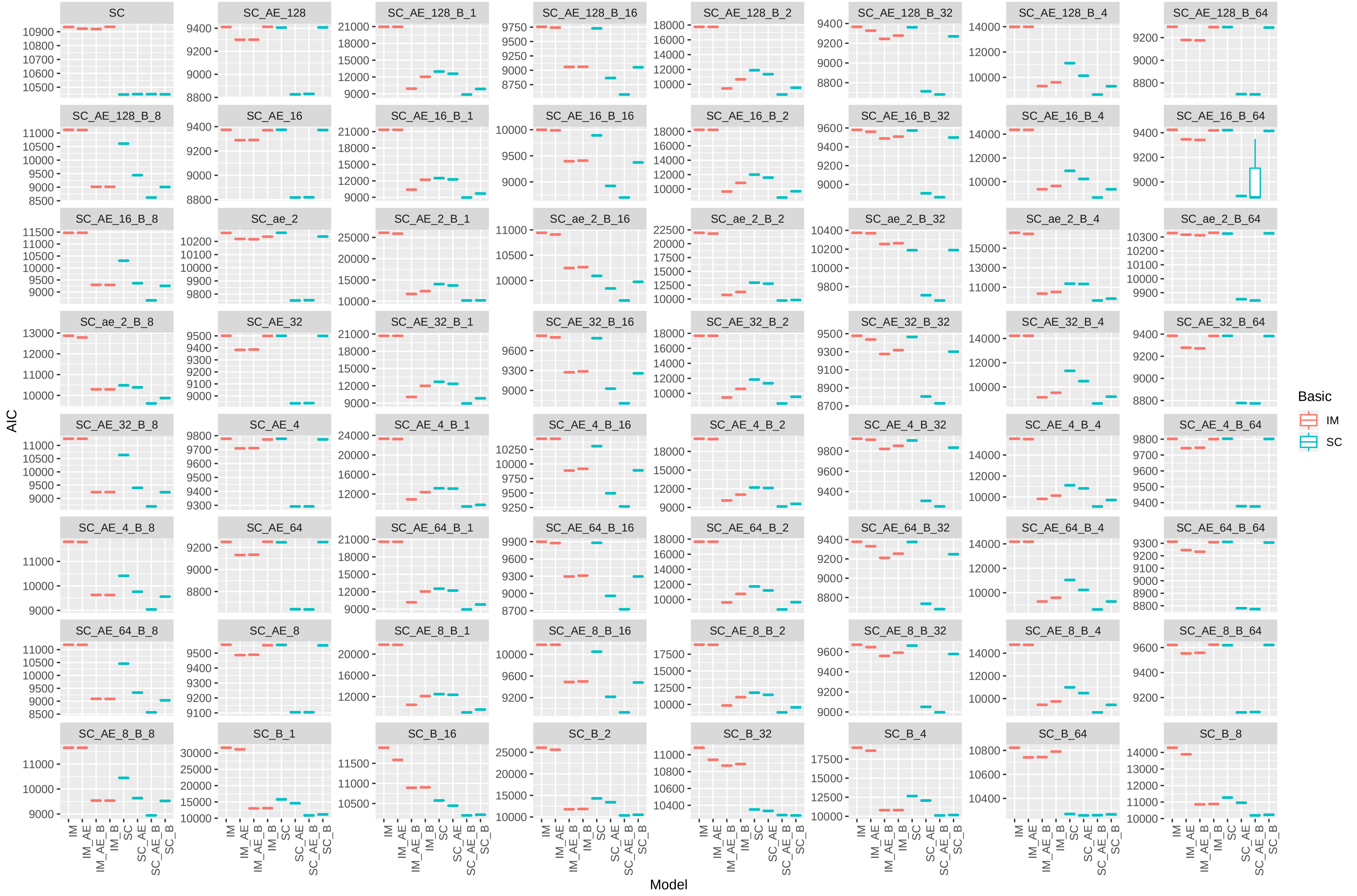


**Supplementary Fig. 9:** Boxplot showing AIC estimates for IM and SC models for the unfolded jAFS of 64 simulations (1 million loci) under the symmetric SC model. The best three replicates for each model/simulation combination are plotted. Note the very narrow boxplots showing models have converged to the same likelihhods, with a few exception.

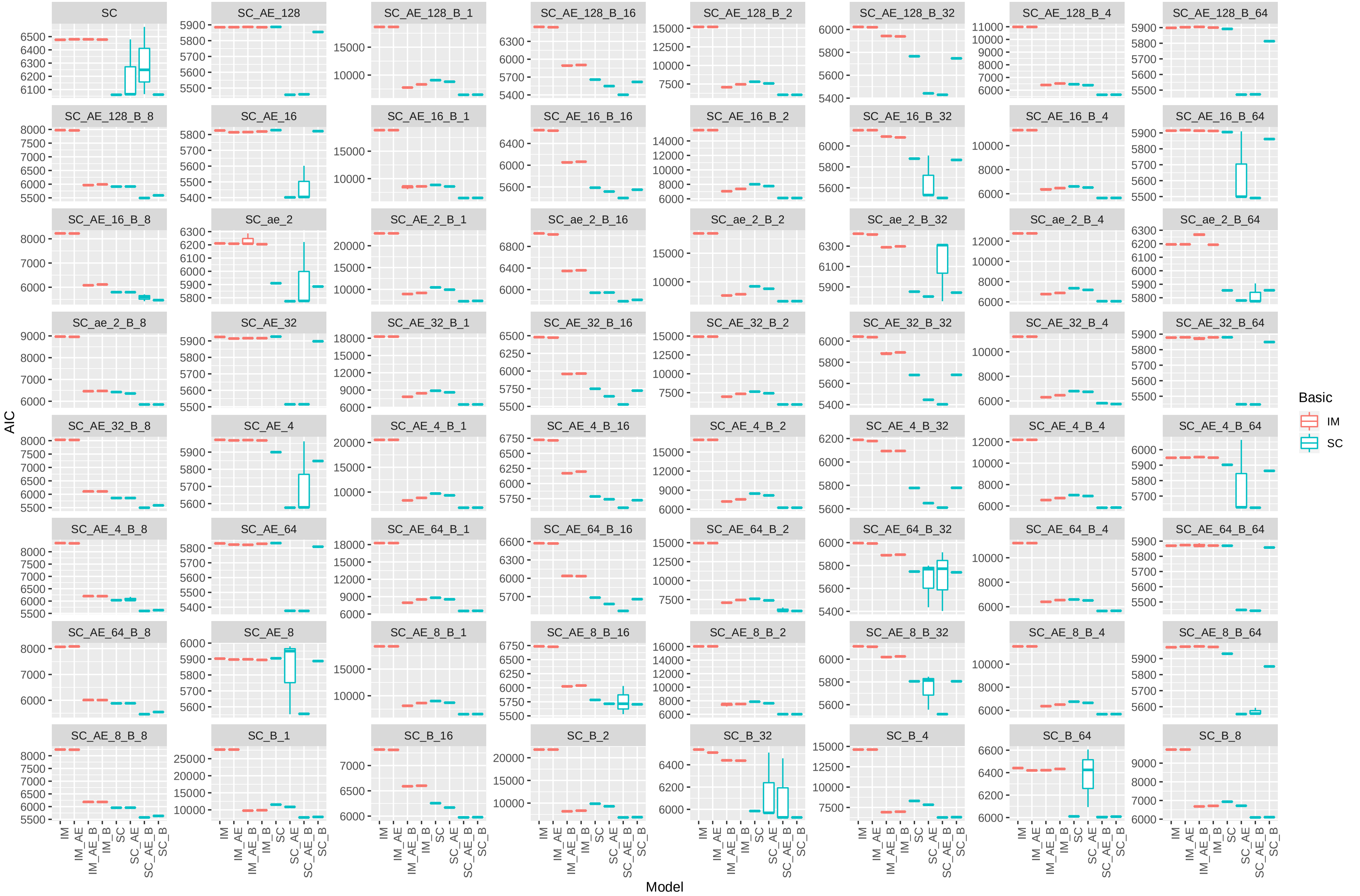


**Supplementary Fig. 10:** Boxplot showing AIC estimates for IM and SC models for the folded jAFS of 64 simulations (1 million loci) under the symmetric SC model. The best three replicates for each model/simulation combination are plotted. Note the very narrow boxplots showing models have converged to the same likelihhods, with a few exception.

**Supplementary Fig. 11:** misestimation of $T_S$. Columns in the grids group by simulation Ancestral Expansion (from 1, i.e. no expansion, to 128 x). The x axes represent the size of population two ($N_2$) at time of split ($T_S$) as a proportion of current $N_2$. Rows of the plot grids group by gene flow scenario of the simulation (IM and SC) and by whether the unfolded or folded spectrum was used for inference. Panels A and B show $T_S$ misestimation when the basic IM and SC models are used for inference, and under a symmetric and asymmetric migration scenario respectively. Panels C and D show ($T_S$) misestimation when the full 8 models (IM,SC, IMAE, $SC_{AE}$, $IM_B$, $SC_B$, $IM_{AEB}$, $SC_{AEB}$ are used. For each model, $T_S$ misestimation was calculated as $T_S^{model}/T_S^{simulation}$.

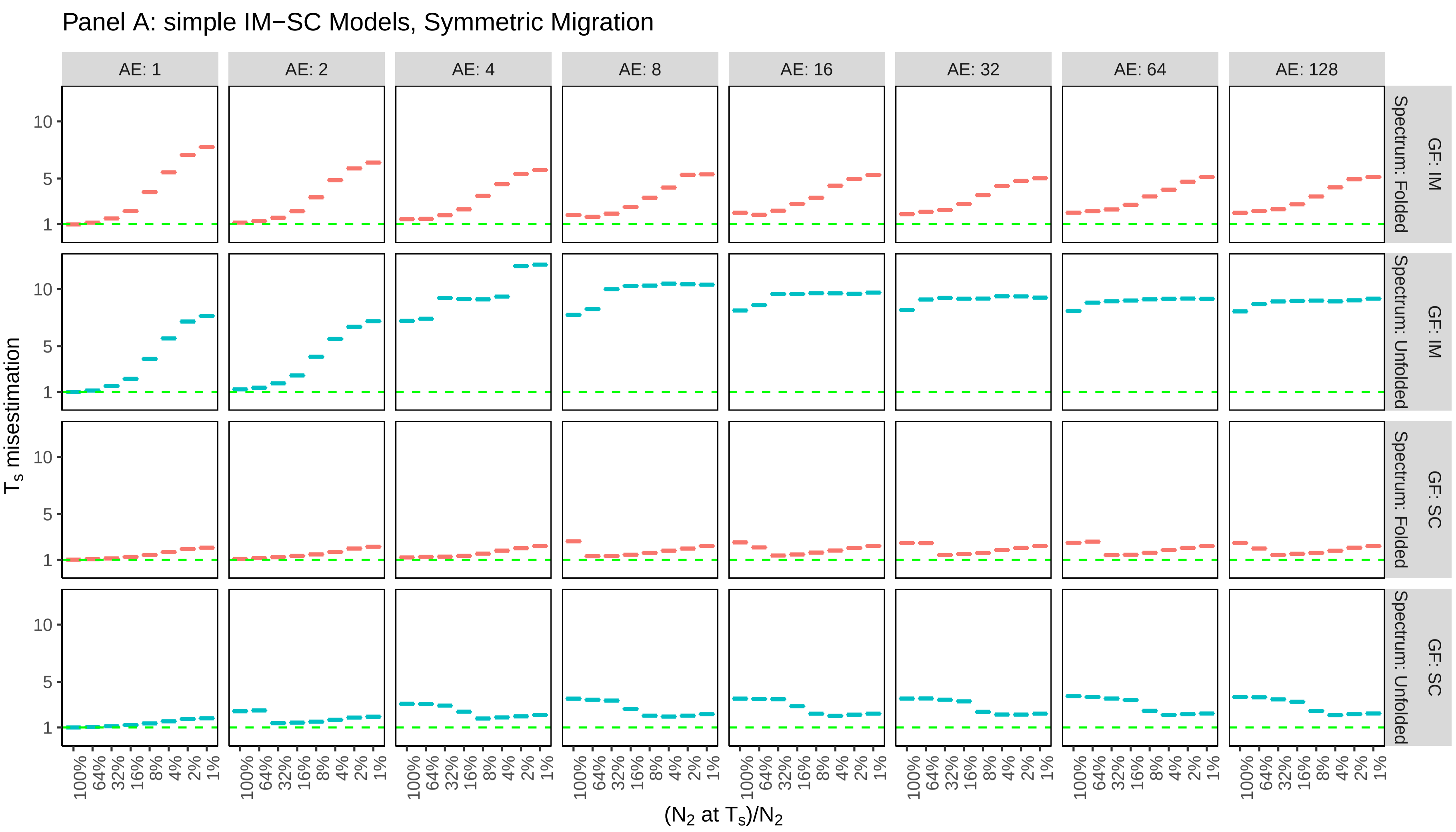

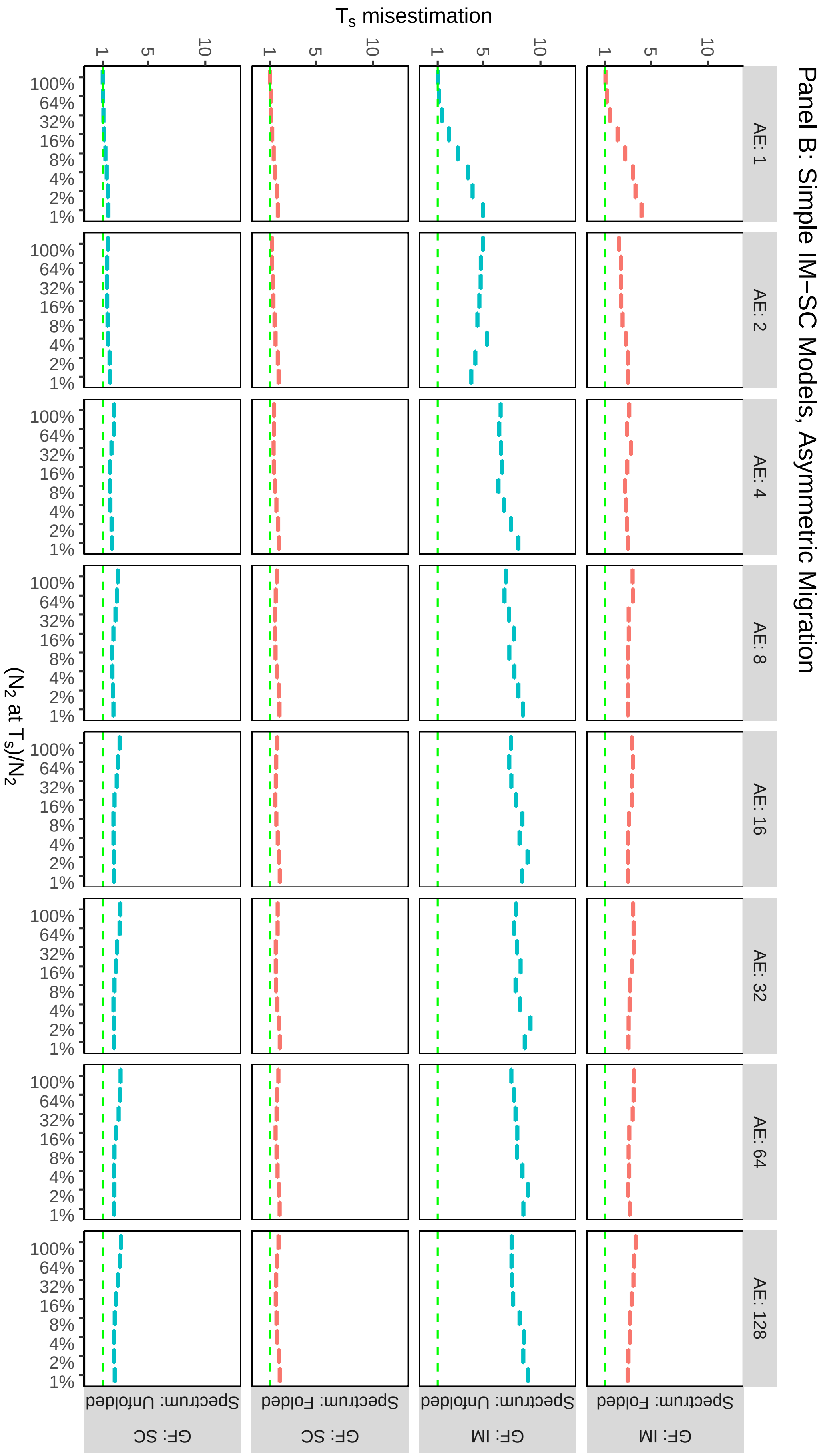

Panel B: Simple IM–SC Models, Asymmetric Migration
$T_s$ misestimation
($N_2$ at $T_s$)/$N_2$
AE: 1
AE: 2
AE: 4
AE: 8
AE: 16
AE: 32
AE: 64
AE: 128
GF: IM
Spectrum: Folded
GF: IM
Spectrum: Unfolded
GF: SC
Spectrum: Folded
GF: SC
Spectrum: Unfolded
100%
64%
32%
16%
8%
4%
2%
1%
1
5
10

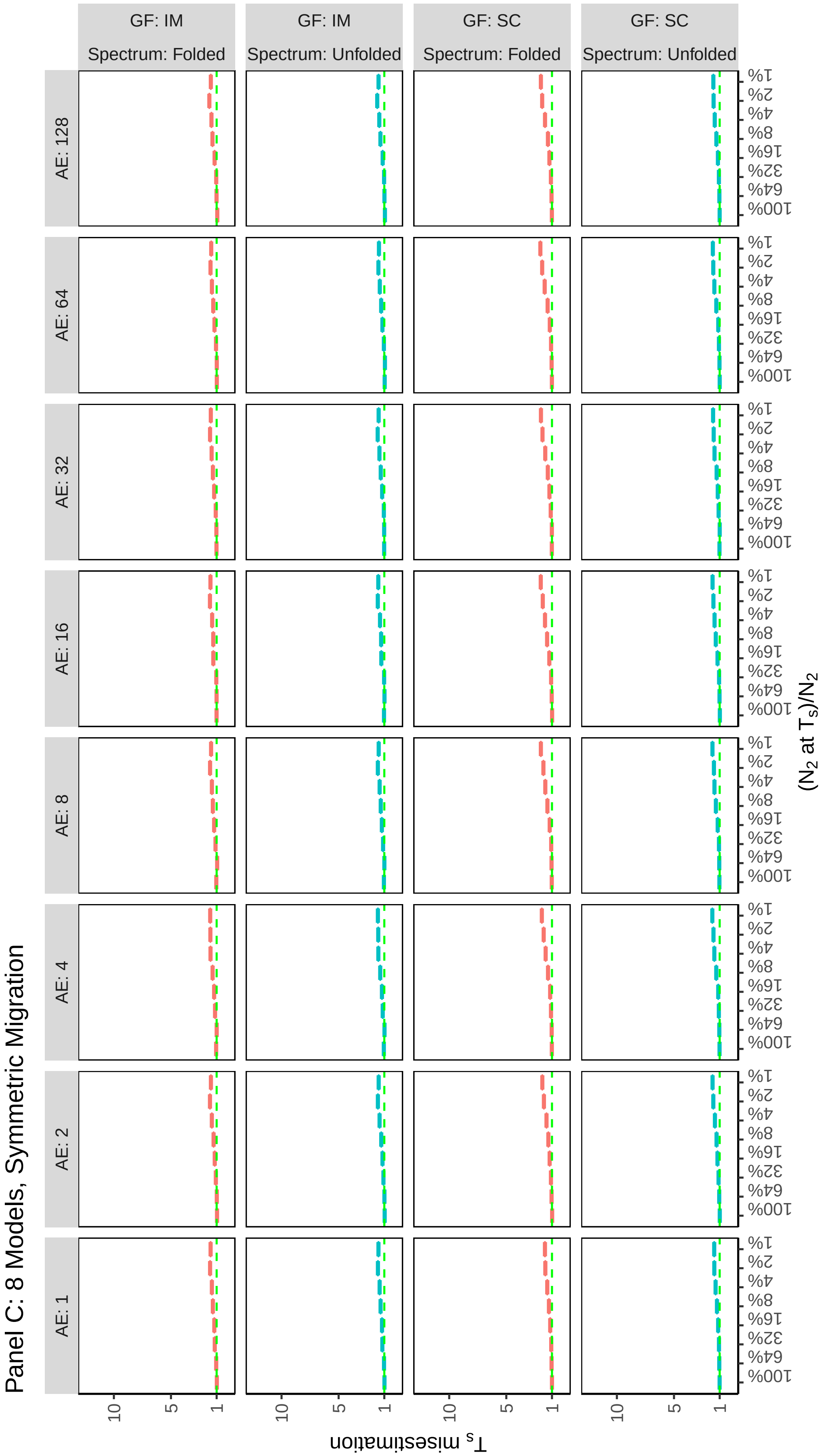
Panel C: 8 Models, Symmetric Migration
GF: IM
Spectrum: Folded
GF: IM
Spectrum: Unfolded
GF: SC
Spectrum: Folded
GF: SC
Spectrum: Unfolded
AE: 128
AE: 64
AE: 32
AE: 16
AE: 8
AE: 4
AE: 2
AE: 1
1%
2%
4%
8%
16%
32%
64%
100%
(N2 at TS)/N2
10
5
1
TS misestimation

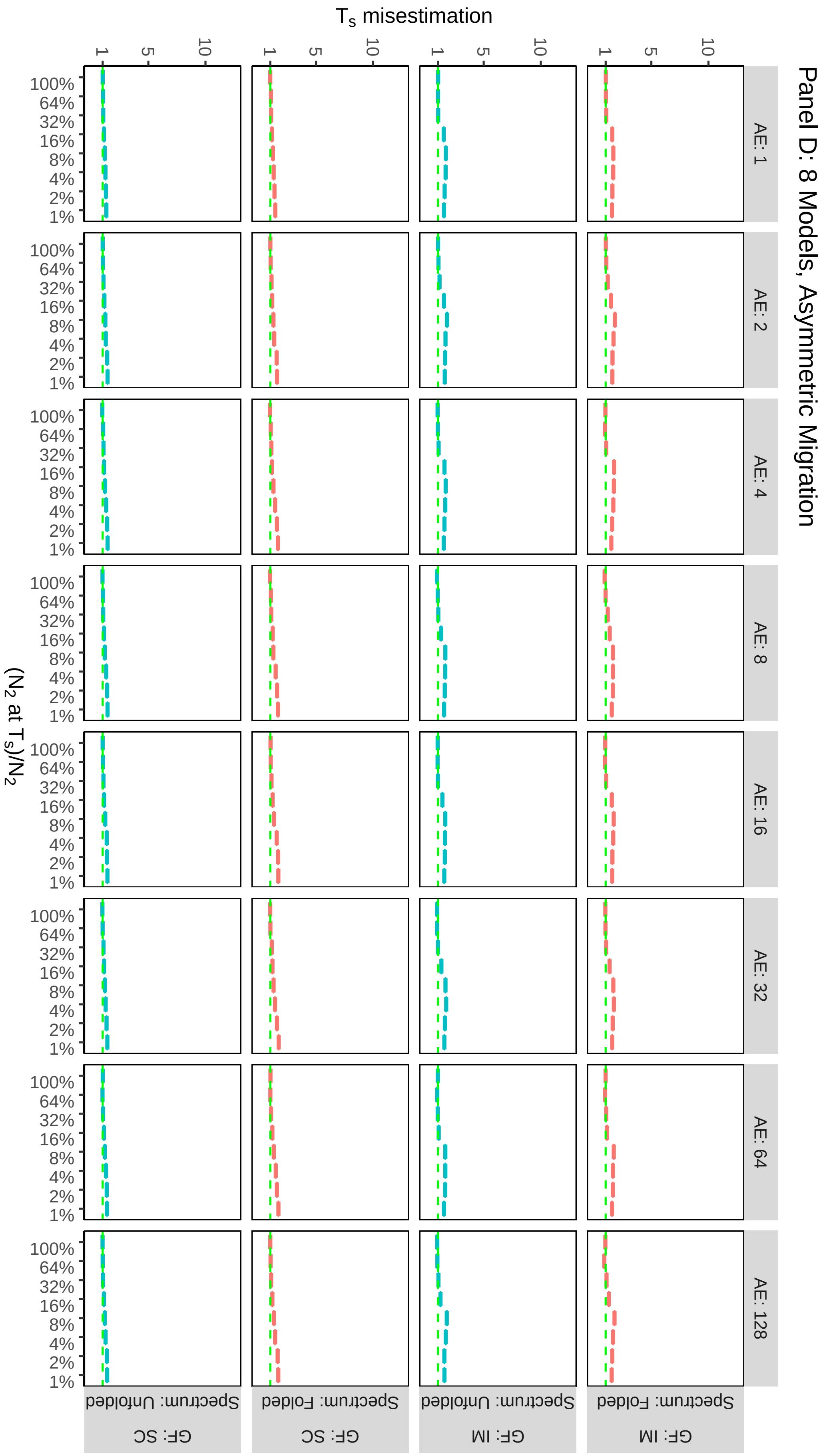
Panel D: 8 Models, Asymmetric Migration
Ts misestimation
(N2 at Ts)/N2
AE: 1
AE: 2
AE: 4
AE: 8
AE: 16
AE: 32
AE: 64
AE: 128
Spectrum: Folded
Spectrum: Unfolded
GF: IM
GF: SC
100%
64%
32%
16%
8%
4%
2%
1%
1
5
10

**Supplementary Fig. 12:** Inferred proportion of SI during divergence by the best model. Columns and rows in the grids group by simulations as for Supplementary Fig. 11. Green dotted lines represent the true proportion of SI for the simulation scenarios. For each simulation the value reported is the value from the model replicate with the lowest AIC.

Panel A: Simple IM−SC Models, Symmetric Migration

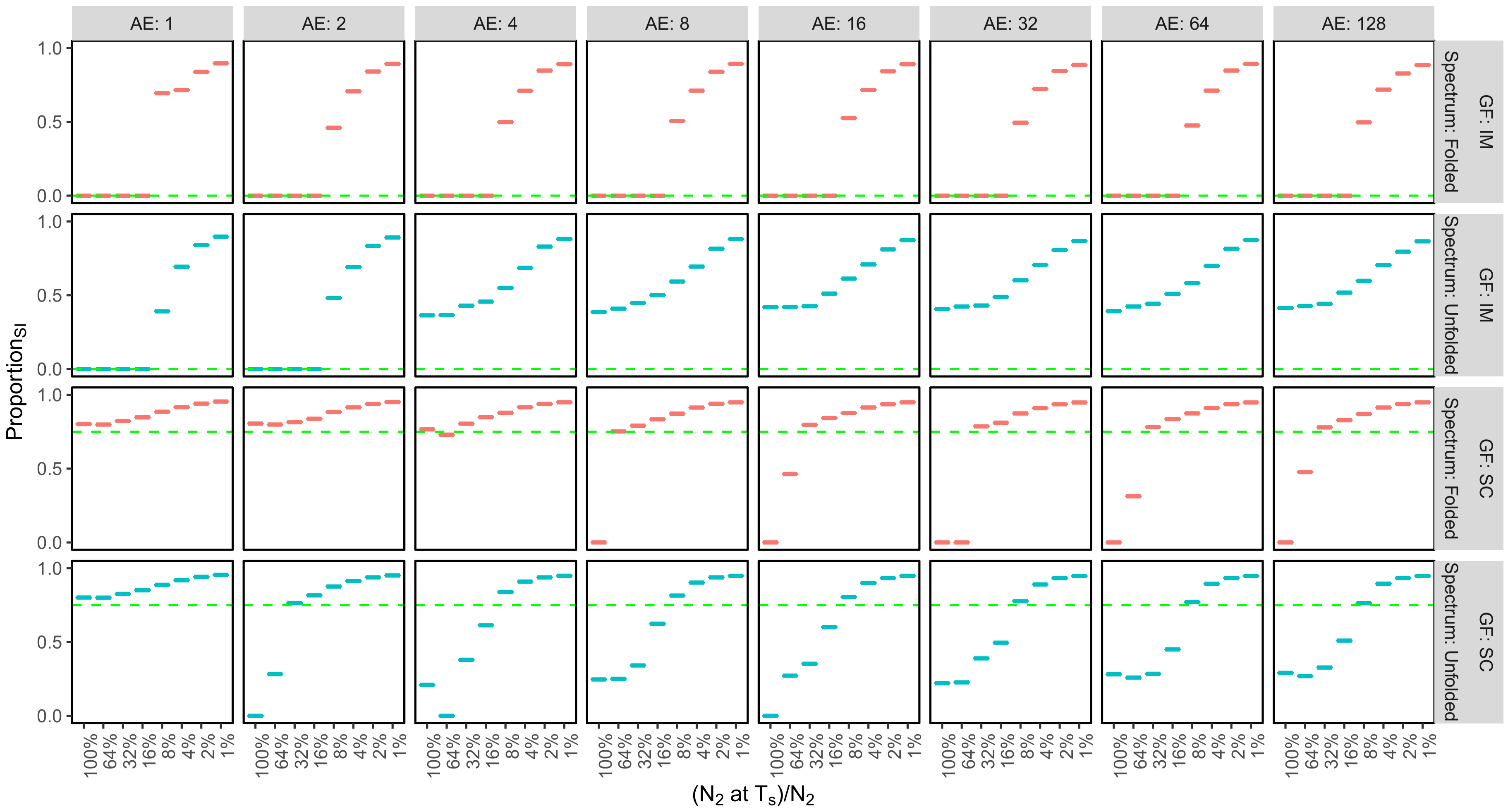

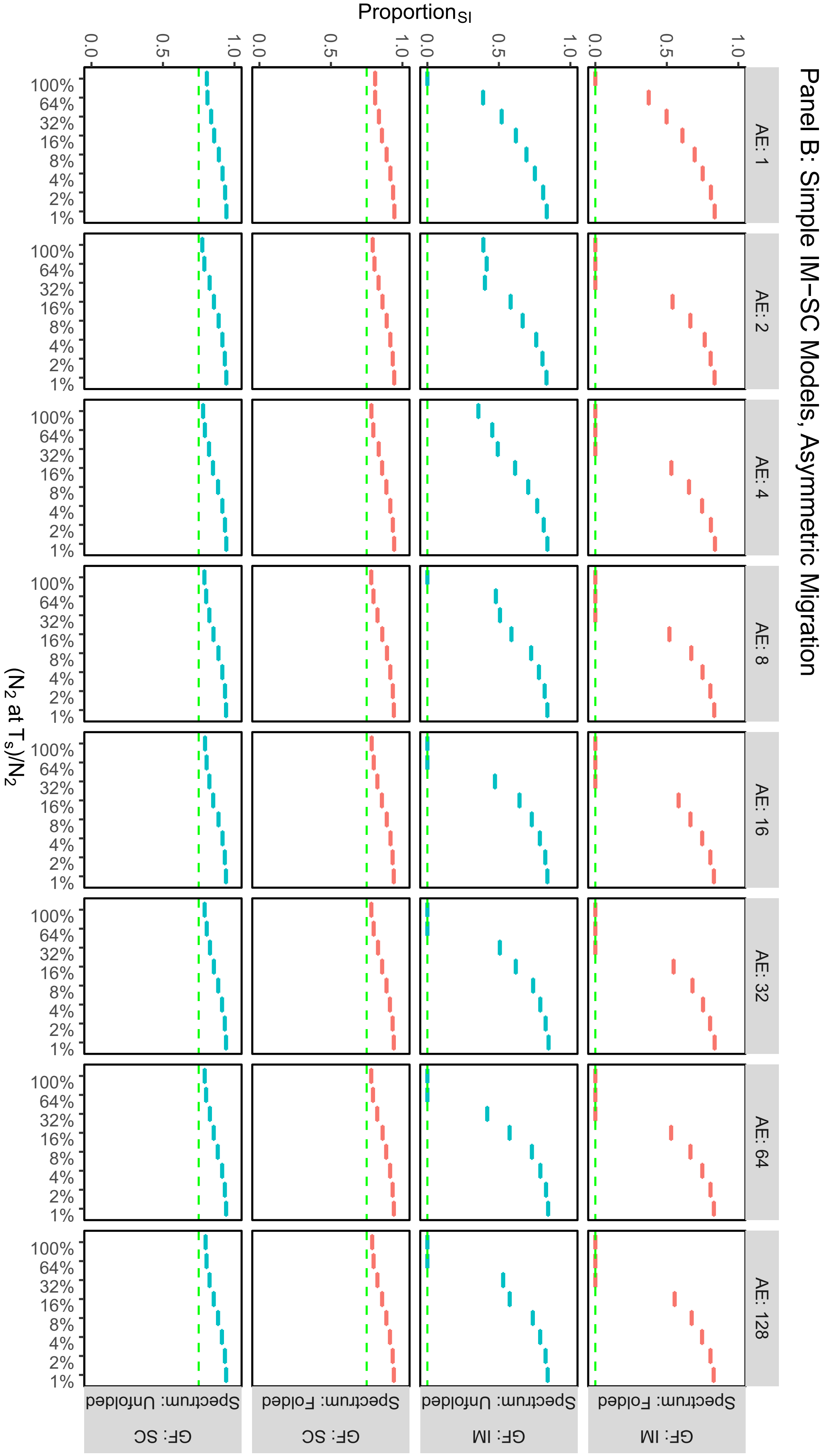

Panel B: Simple IM–SC Models, Asymmetric Migration
Proportion$_{SI}$
0.0
0.5
1.0
AE: 1
AE: 2
AE: 4
AE: 8
AE: 16
AE: 32
AE: 64
AE: 128
100%
64%
32%
16%
8%
4%
2%
1%
($N_2$ at $T_s$)/$N_2$
Spectrum: Folded
Spectrum: Unfolded
GF: IM
GF: SC

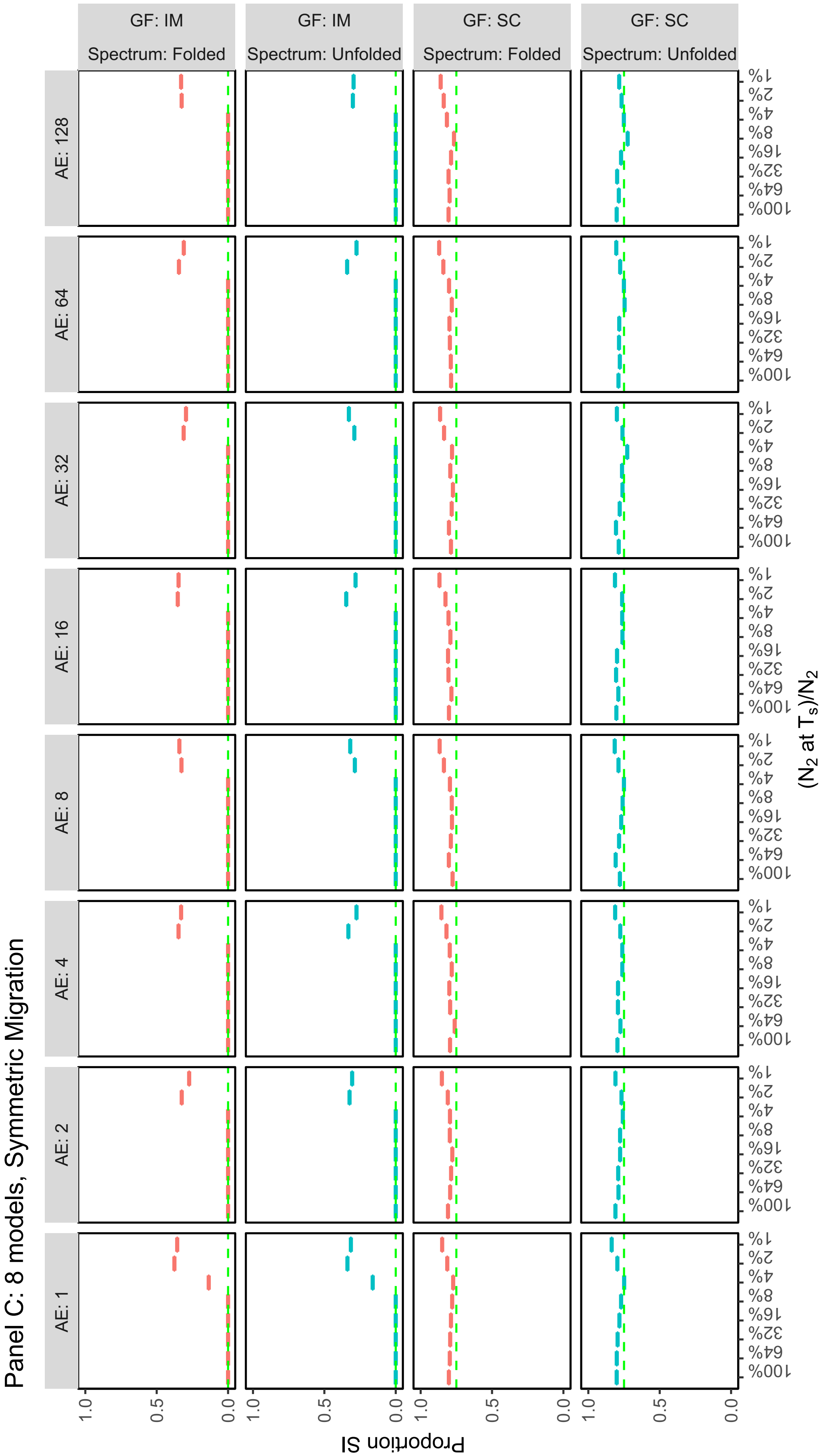

Panel C: 8 models, Symmetric Migration
GF: IM
Spectrum: Folded
GF: IM
Spectrum: Unfolded
GF: SC
Spectrum: Folded
GF: SC
Spectrum: Unfolded
AE: 1
AE: 2
AE: 4
AE: 8
AE: 16
AE: 32
AE: 64
AE: 128
1%
2%
4%
8%
16%
32%
64%
100%
1.0
0.5
0.0
Proportion SI
(N2 at Ts)/N2

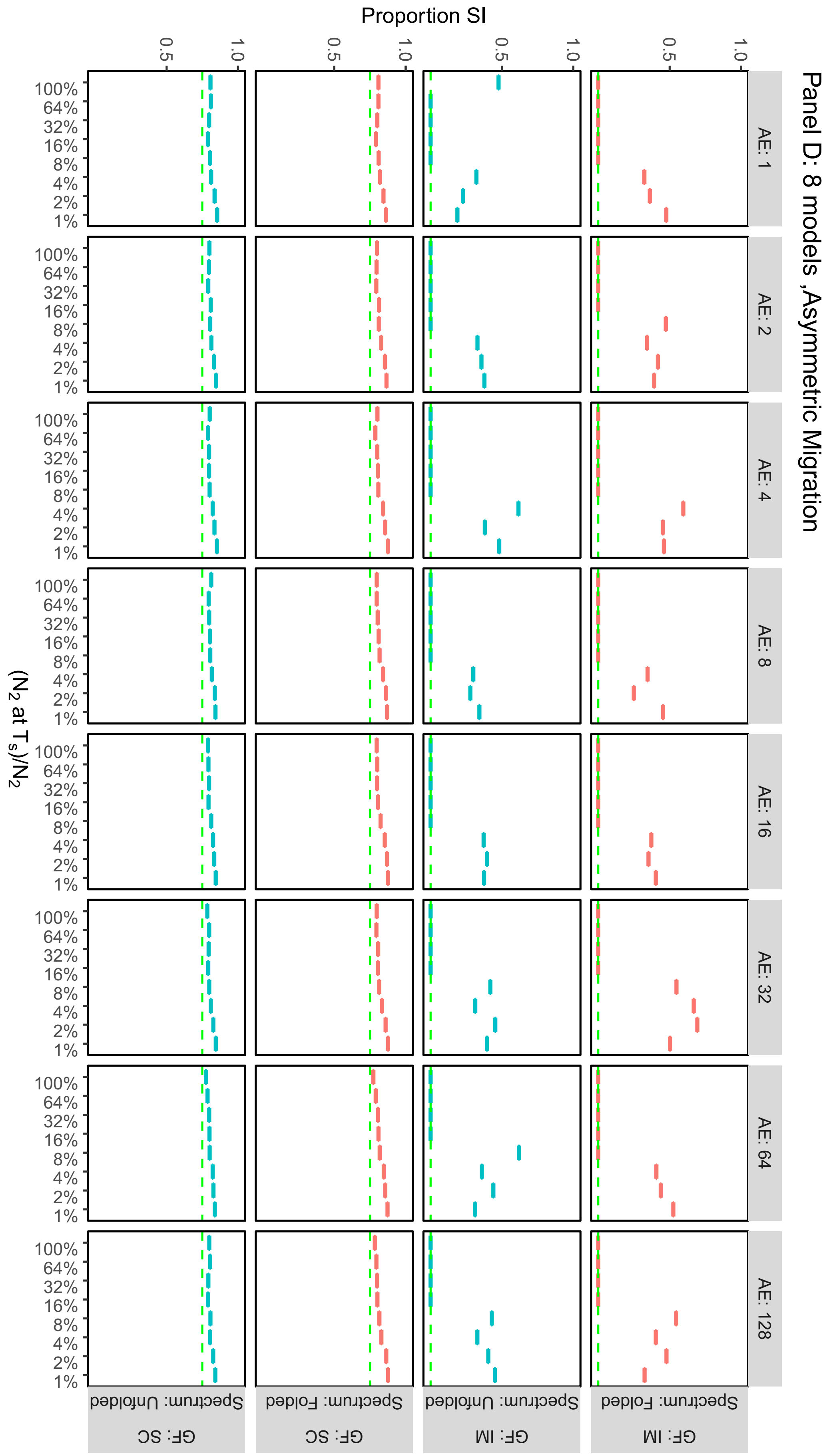
Panel D: 8 models ,Asymmetric Migration
AE: 1
AE: 2
AE: 4
AE: 8
AE: 16
AE: 32
AE: 64
AE: 128
Proportion SI
0.5
1.0
(N2 at Ts)/N2
100%
64%
32%
16%
8%
4%
2%
1%
GF: IM
Spectrum: Folded
GF: IM
Spectrum: Unfolded
GF: SC
Spectrum: Folded
GF: SC
Spectrum: Unfolded

**Supplementary Fig. 13:** misestimation of $N_{ANC}$. Columns in the grids group by simulation Ancestral Expansion (from 1, i.e. no expansion, to 128 x). The x axes represent the size of population 2 ($N_2$) at time of split ($T_S$) as a proportion of current ($N_2$). Rows of the plot grids group by gene flow scenario of the simulation (IM and SC) and by whether the unfolded or folded spectrum was used for inference. Panels **A** and **B** show $N_{ANC}$ misestimation when the basic IM and SC models are used for inference, and under a symmetric and asymmetric migration scenario respectively. Panels **C** and **D** show $N_{ANC}$ misestimation when the full 8 models (IM,SC,$IM_{AE}$, $SC_{AE}$, $IM_B$, $SC_B$, $IM_{AEB}$, $SC_{AEB}$ are used. For each model,$N_{ANC}$ misestimation was calculated as $N_{ANC}$model/ $N_{ANC}$ simulation.

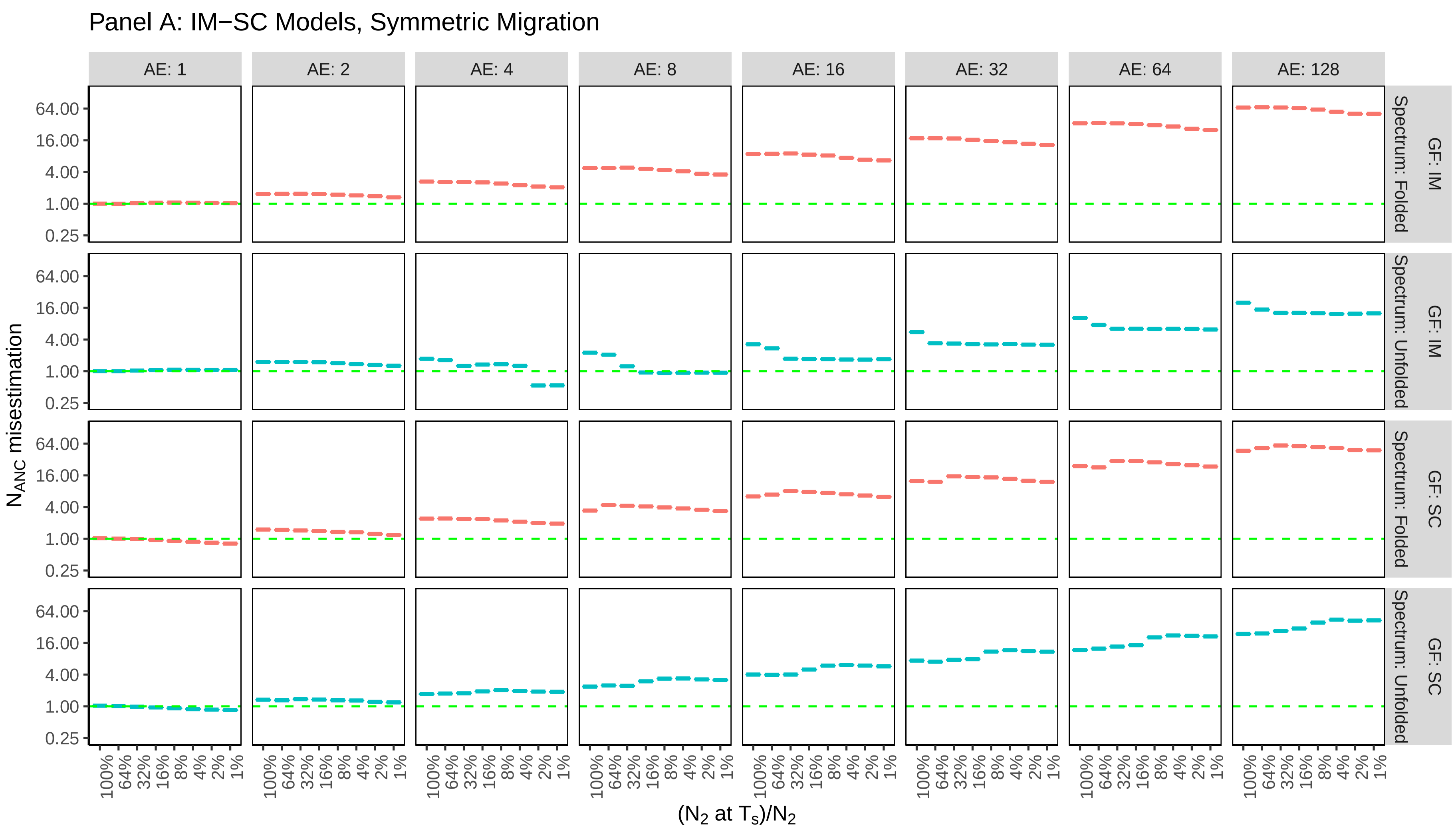

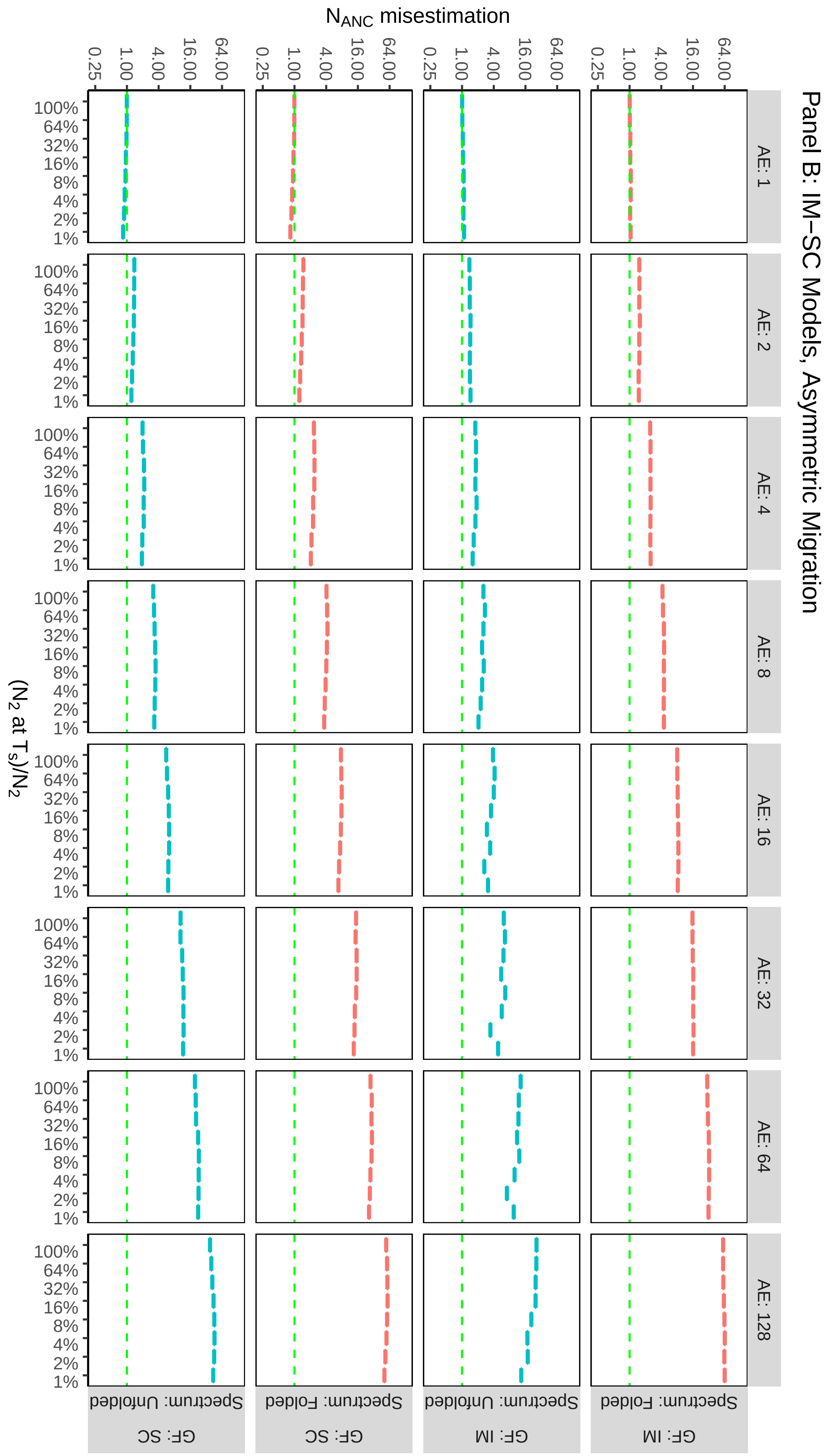
Panel B: IM–SC Models, Asymmetric Migration
N_ANC misestimation
0.25
1.00
4.00
16.00
64.00
(N2 at Ts)/N2
100%
64%
32%
16%
8%
4%
2%
1%
AE: 1
AE: 2
AE: 4
AE: 8
AE: 16
AE: 32
AE: 64
AE: 128
Spectrum: Unfolded
Spectrum: Folded
GF: SC
GF: IM

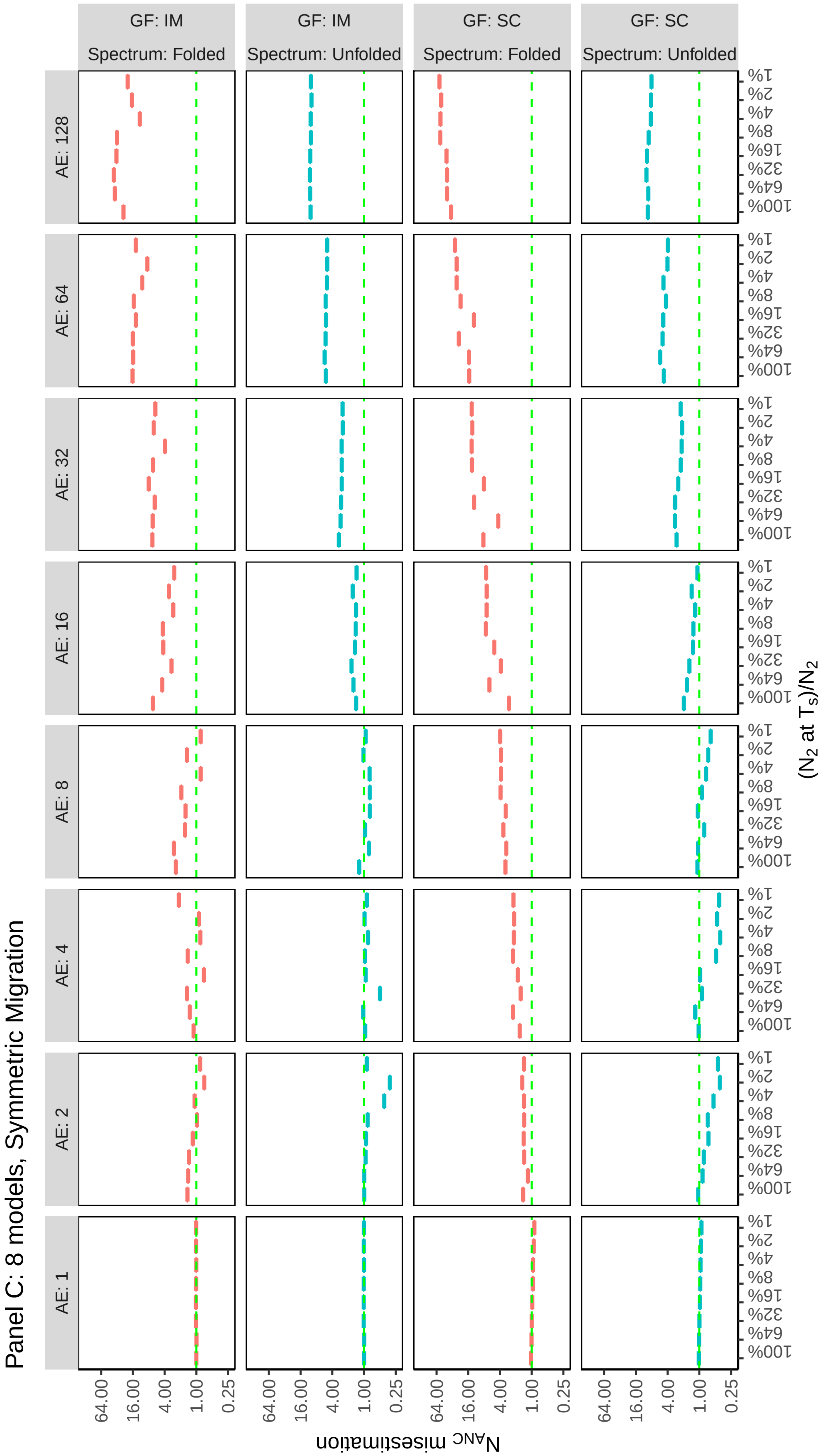
Panel C: 8 models, Symmetric Migration
GF: IM
Spectrum: Folded
GF: IM
Spectrum: Unfolded
GF: SC
Spectrum: Folded
GF: SC
Spectrum: Unfolded
AE: 128
AE: 64
AE: 32
AE: 16
AE: 8
AE: 4
AE: 2
AE: 1
64.00
16.00
4.00
1.00
0.25
$N_{ANC}$ misestimation
1%
2%
4%
8%
16%
32%
64%
100%
($N_2$ at $T_S$)/$N_2$

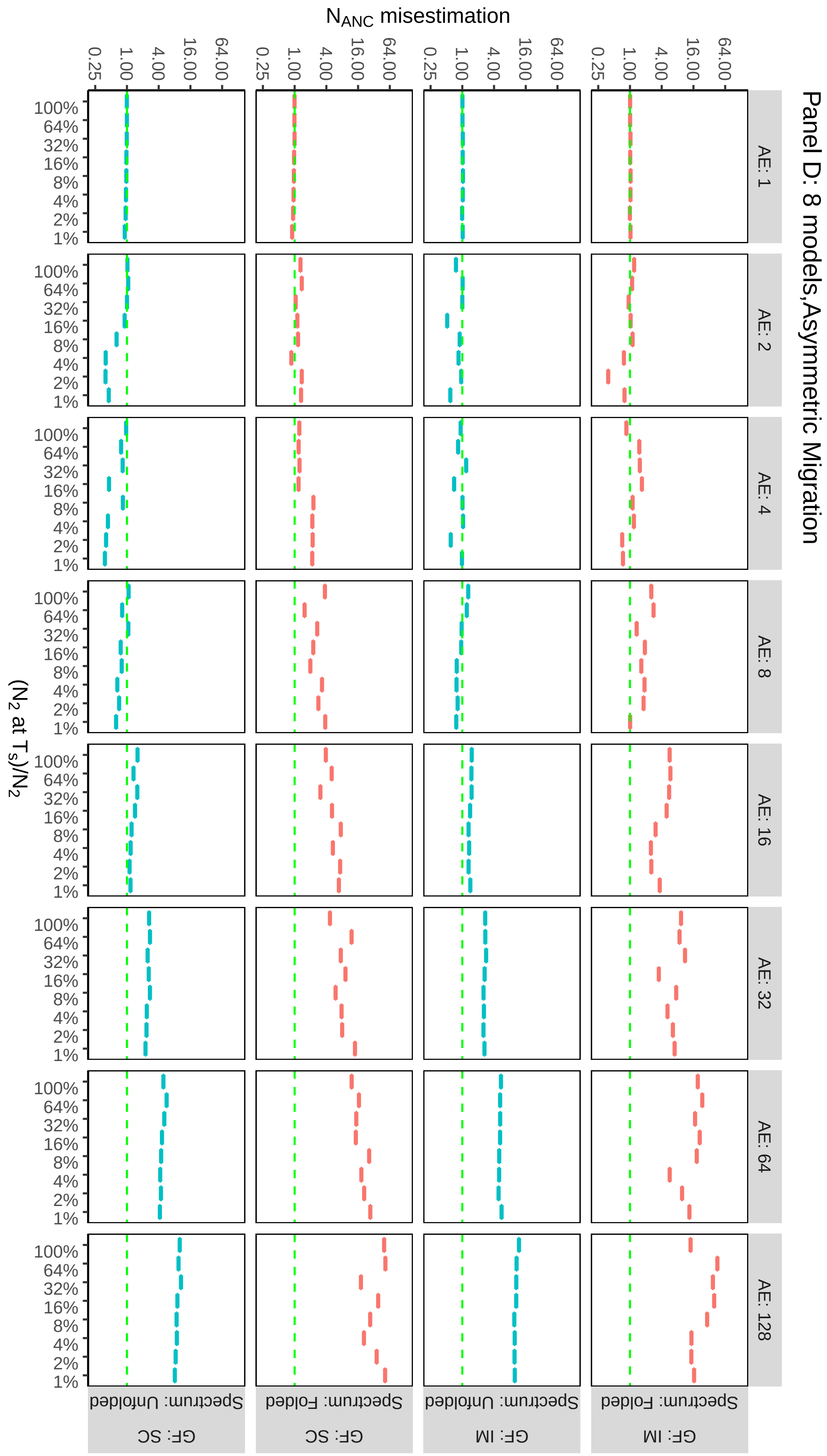

Panel D: 8 models,Asymmetric Migration
$N_{ANC}$ misestimation
0.25
1.00
4.00
16.00
64.00
AE: 1
AE: 2
AE: 4
AE: 8
AE: 16
AE: 32
AE: 64
AE: 128
100%
64%
32%
16%
8%
4%
2%
1%
($N_2$ at $T_S$)/$N_2$
Spectrum: Unfolded
Spectrum: Folded
GF: SC
GF: IM

**Supplementary Fig. 14:** misestimation of $m_{12}$. Columns is the grids group by simulation Ancestral Expansion (from 1, i.e. no expansion, to 128 x). The x axes represent the size of population two ($N_2$) at time of split ($T_S$) as a proportion of current ($N_2$). Rows of the plot grids group by gene flow scenario of the simulation (IM and SC) and by whether the unfolded or folded spectrum was used for inference. Panels **A** and **B** show $m_{12}$ misestimation when the basic IM and SC models are used for inference, and under a symmetric and asymmetric migration scenario respectively. Panels **C** and **D** show $m_{12}$ misestimation when the full 8 models (IM,SC,$IM_{AE}$, $SC_{AE}$, $IM_B$, $SC_B$, $IM_{AEB}$, $SC_{AEB}$ are used. For each model, $m_{12}$ misestimation was calculated as $m_{12}^{model}/m_{12}^{simulation}$

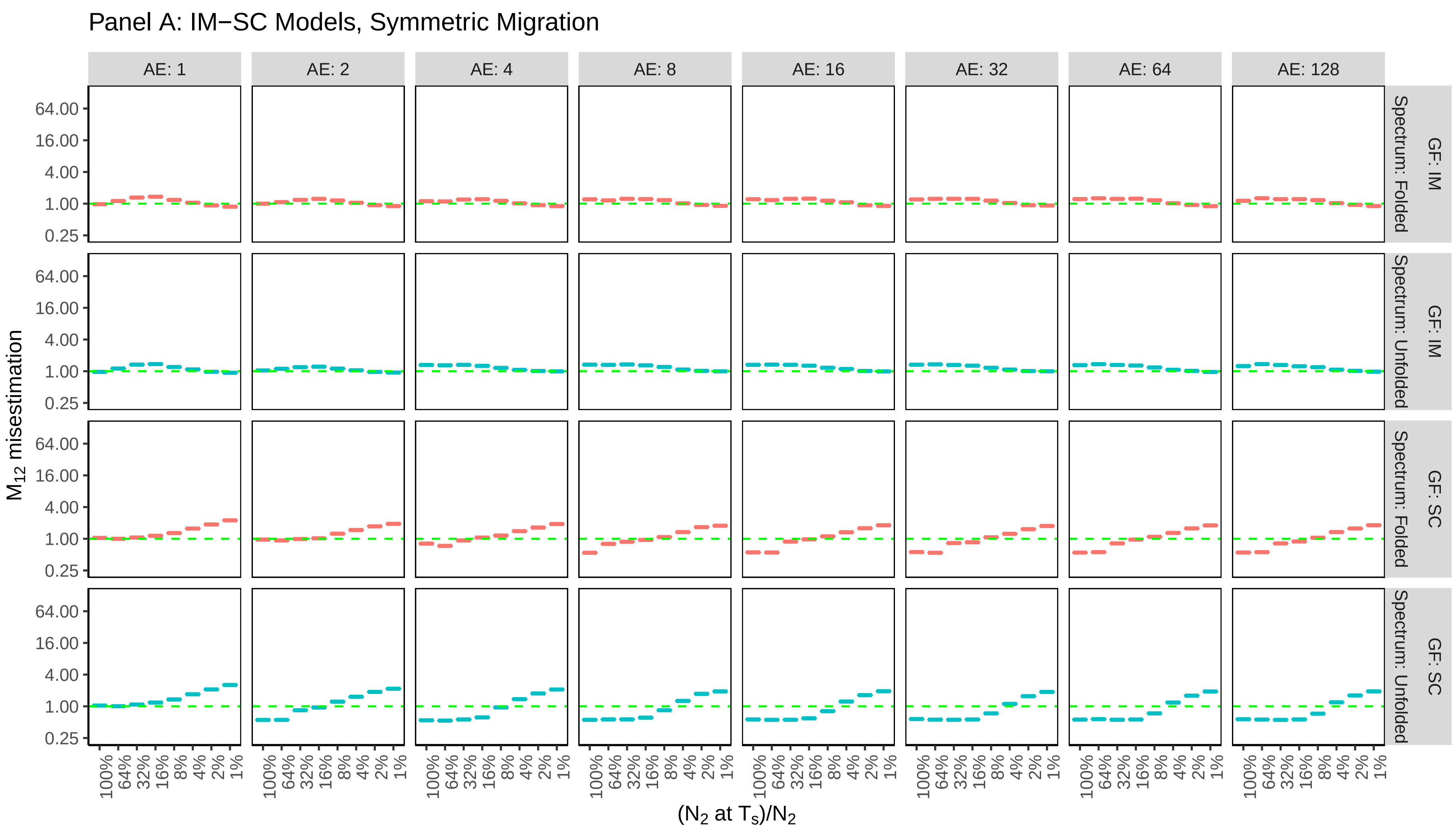

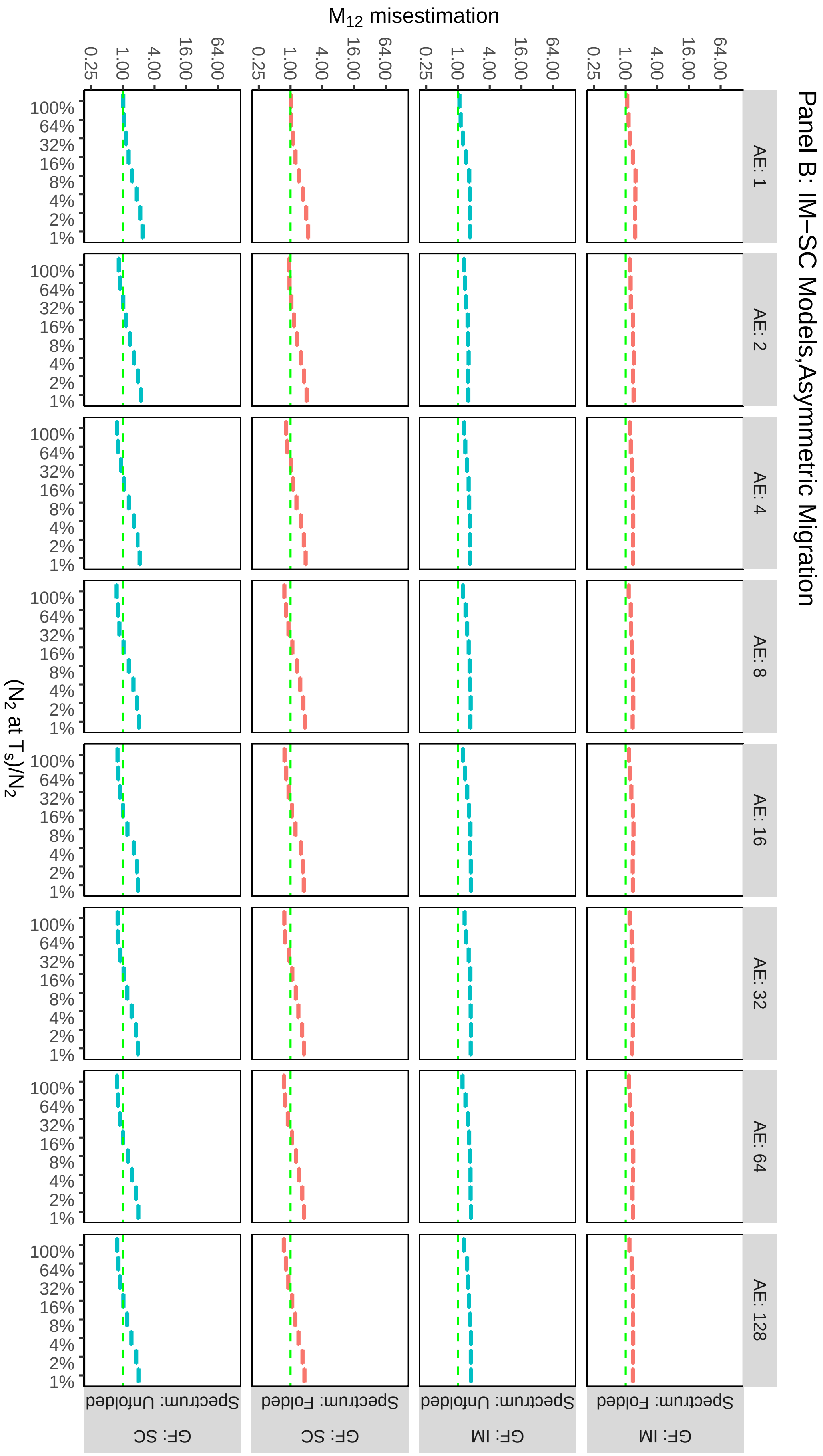

Panel B: IM–SC Models,Asymmetric Migration
M12 misestimation
0.25
1.00
4.00
16.00
64.00
AE: 1
AE: 2
AE: 4
AE: 8
AE: 16
AE: 32
AE: 64
AE: 128
100%
64%
32%
16%
8%
4%
2%
1%
(N2 at Ts)/N2
GF: IM
Spectrum: Folded
GF: IM
Spectrum: Unfolded
GF: SC
Spectrum: Folded
GF: SC
Spectrum: Unfolded

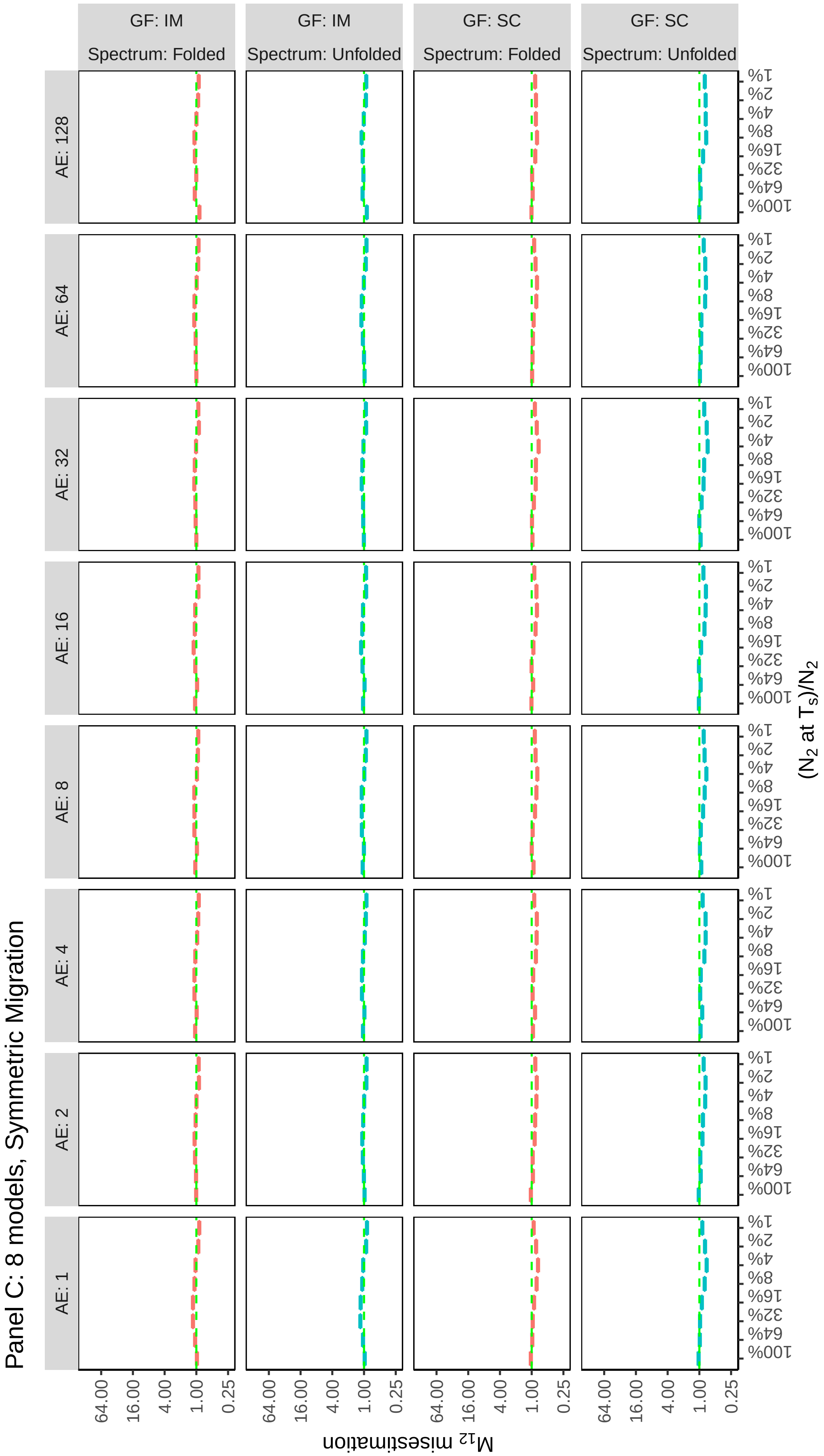
Panel C: 8 models, Symmetric Migration
GF: IM
Spectrum: Folded
GF: IM
Spectrum: Unfolded
GF: SC
Spectrum: Folded
GF: SC
Spectrum: Unfolded
AE: 1
AE: 2
AE: 4
AE: 8
AE: 16
AE: 32
AE: 64
AE: 128
64.00
16.00
4.00
1.00
0.25
$M_{12}$ misestimation
1%
2%
4%
8%
16%
32%
64%
100%
($N_2$ at $T_S$)/$N_2$

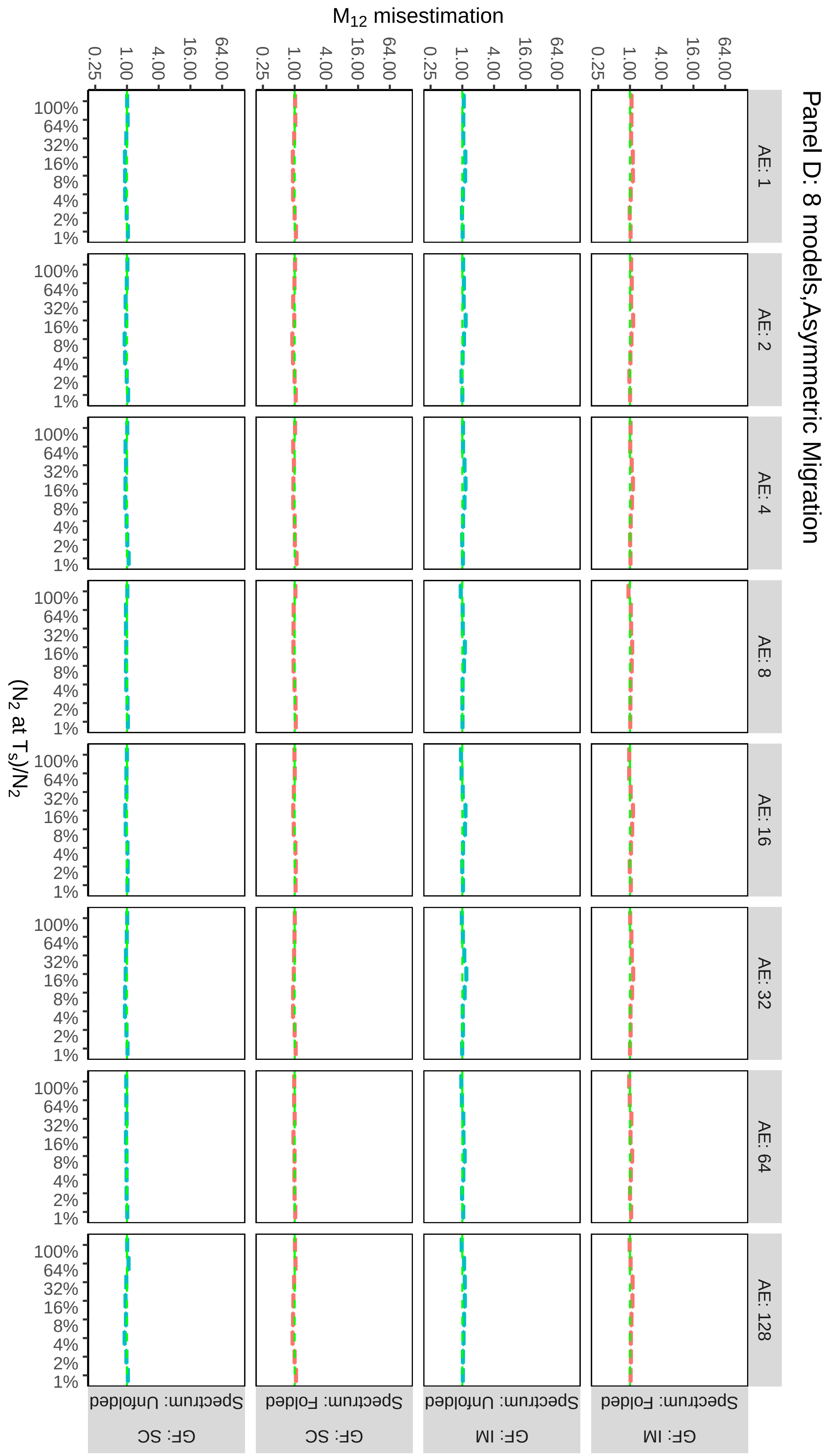
Panel D: 8 models,Asymmetric Migration
M12 misestimation
0.25
1.00
4.00
16.00
64.00
AE: 1
AE: 2
AE: 4
AE: 8
AE: 16
AE: 32
AE: 64
AE: 128
(N2 at Ts)/N2
100%
64%
32%
16%
8%
4%
2%
1%
Spectrum: Unfolded
Spectrum: Folded
GF: SC
GF: IM

**Supplementary Fig. 15:** misestimation of $m_{21}$. Columns is the grids group by simulation Ancestral Expansion (from 1, i.e. no expansion, to 128 x). The x axes represent the size of population two ($N_2$) at time of split ($T_S$) as a proportion of current ($N_2$). Rows of the plot grids group by gene flow scenario of the simulation (IM and SC) and by whether the unfolded or folded spectrum was used for inference. Panels **A** and **B** show $m_{21}$ misestimation when the basic IM and SC models are used for inference, and under a symmetric and asymmetric migration scenario respectively. Panels **C** and **D** show ($N_2$) misestimation when the full 8 models (IM,SC,$IM_{AE}$, $SC_{AE}$, $IM_B$, $SC_B$, $IM_{AEB}$, $SC_{AEB}$ are used. For each model, $m_{21}$ misestimation was calculated as $m_{21}^{model}/m_{21}^{simulation}$

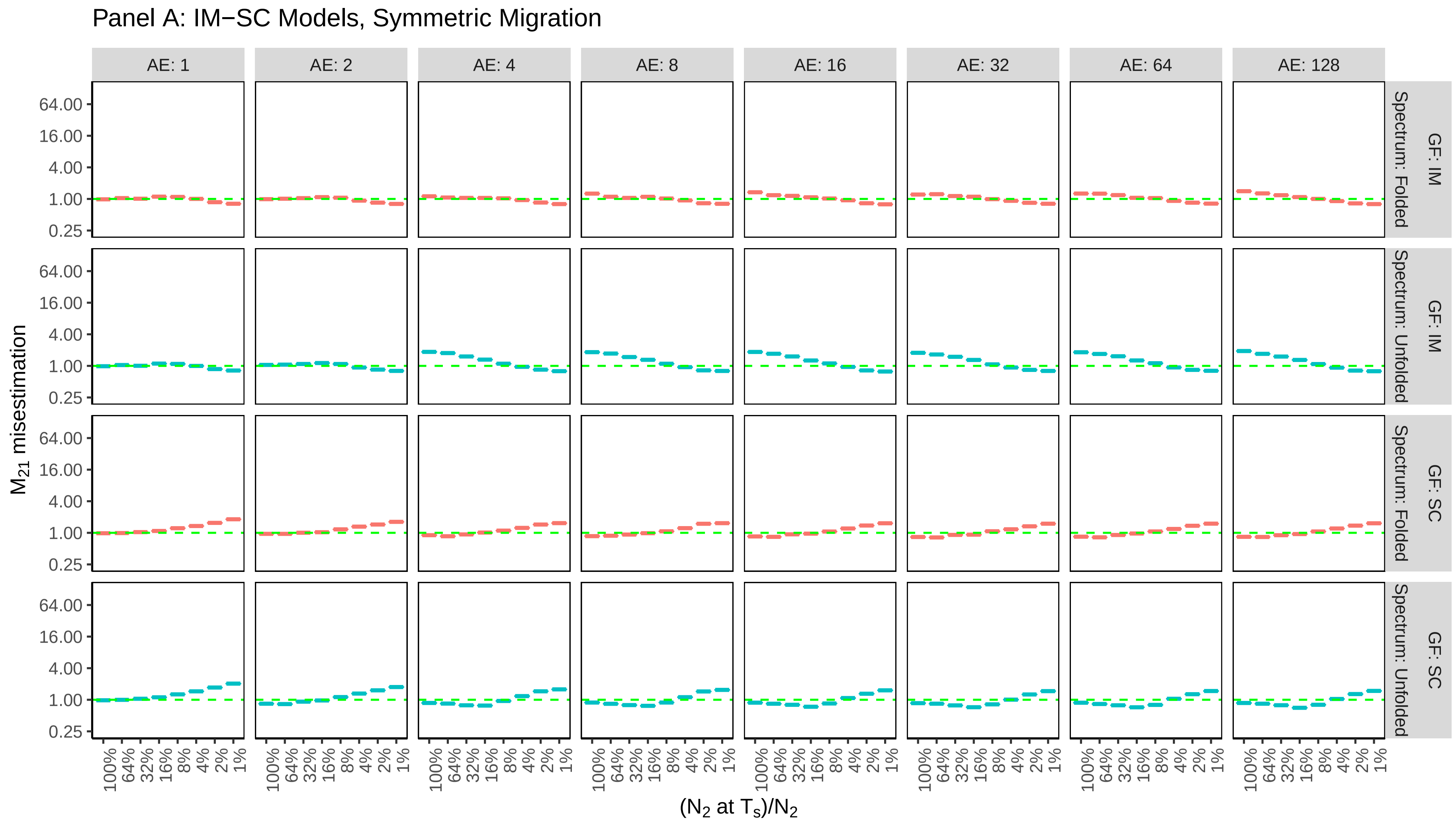

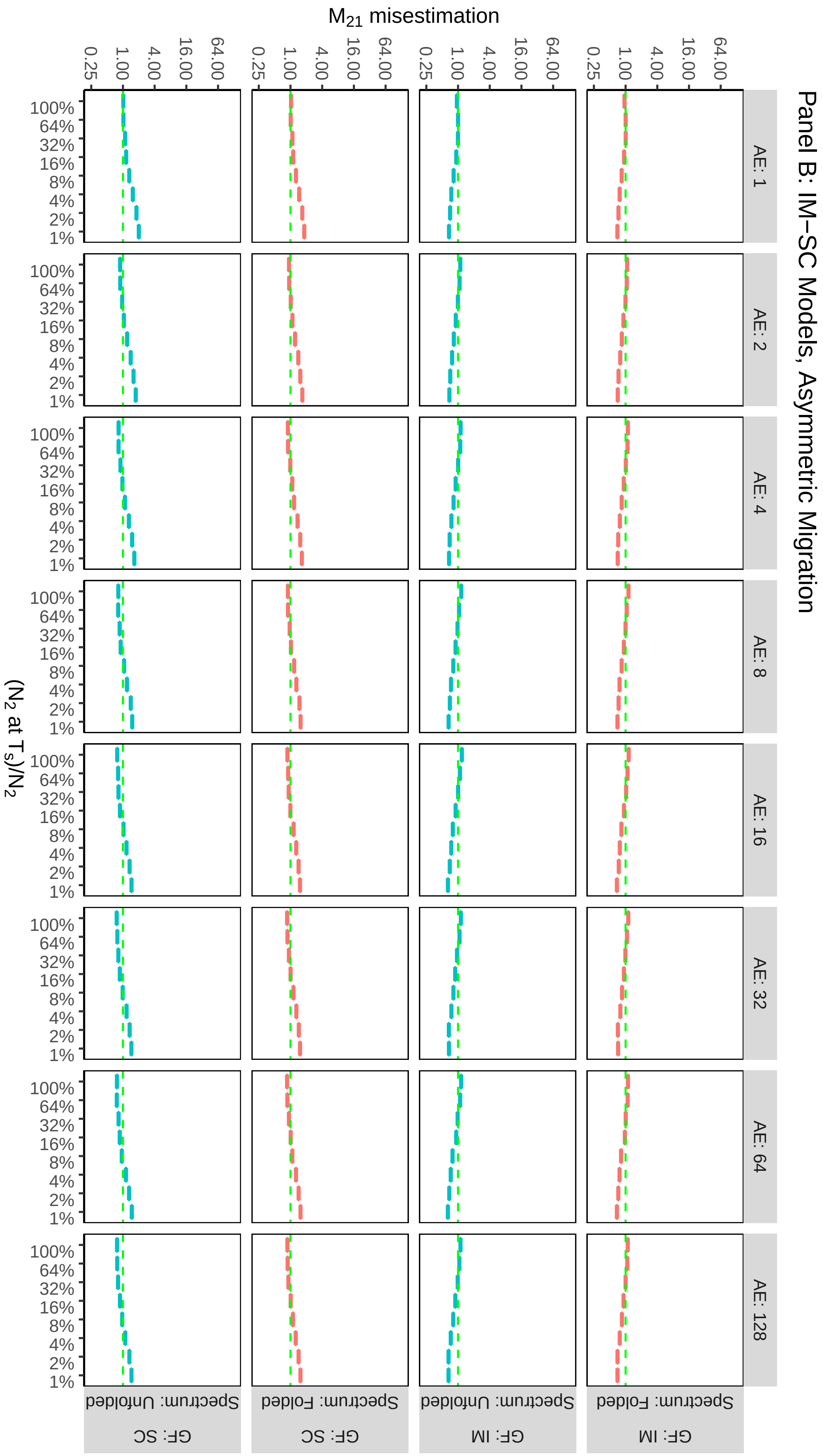
Panel B: IM–SC Models, Asymmetric Migration
M21 misestimation
0.25
1.00
4.00
16.00
64.00
(N2 at Ts)/N2
100%
64%
32%
16%
8%
4%
2%
1%
AE: 1
AE: 2
AE: 4
AE: 8
AE: 16
AE: 32
AE: 64
AE: 128
GF: IM
Spectrum: Folded
GF: IM
Spectrum: Unfolded
GF: SC
Spectrum: Folded
GF: SC
Spectrum: Unfolded

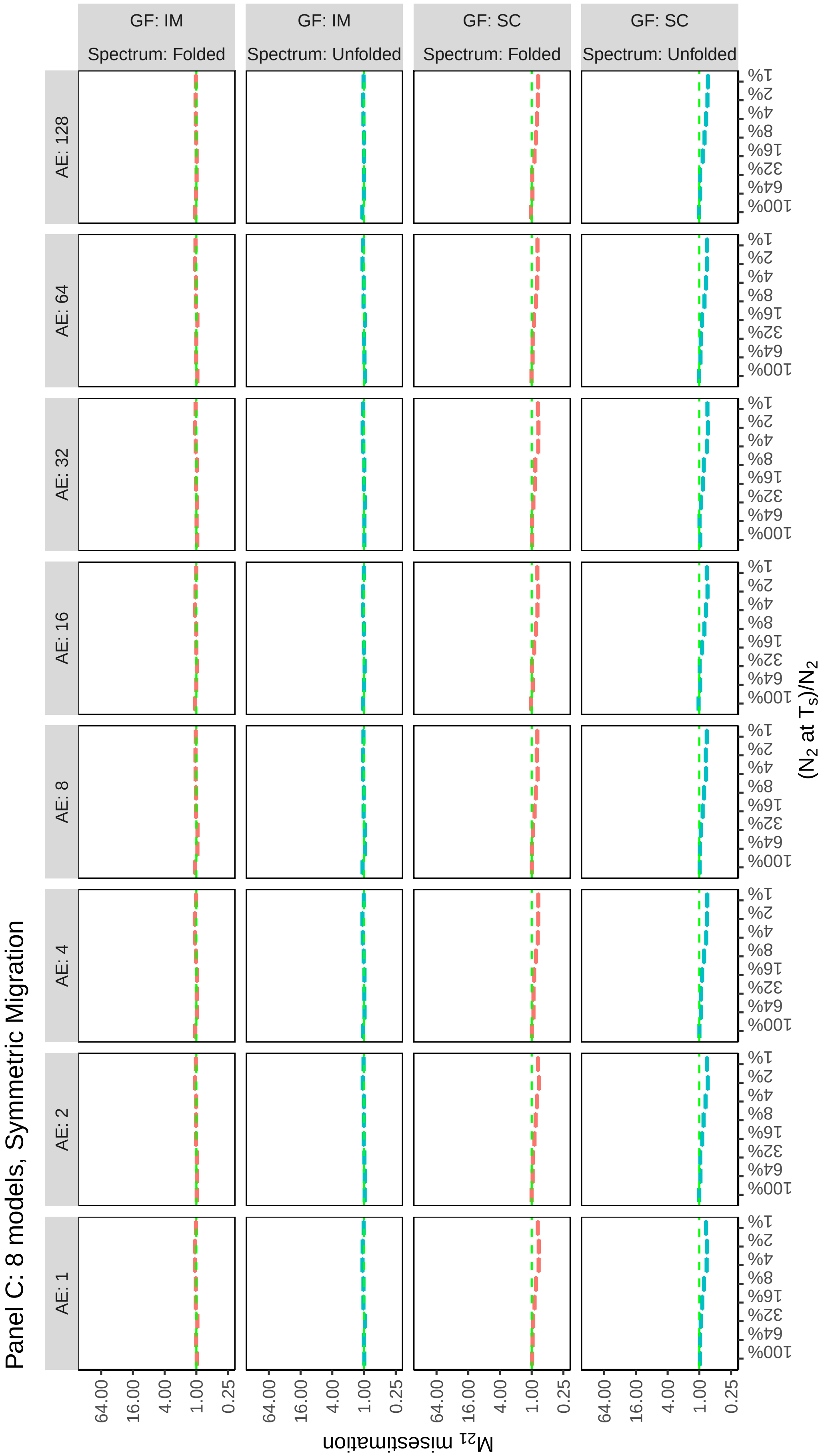
Panel C: 8 models, Symmetric Migration
GF: IM
Spectrum: Folded
GF: IM
Spectrum: Unfolded
GF: SC
Spectrum: Folded
GF: SC
Spectrum: Unfolded
AE: 1
AE: 2
AE: 4
AE: 8
AE: 16
AE: 32
AE: 64
AE: 128
64.00
16.00
4.00
1.00
0.25
$M_{21}$ misestimation
1%
2%
4%
8%
16%
32%
64%
100%
($N_2$ at $T_S$)/$N_2$

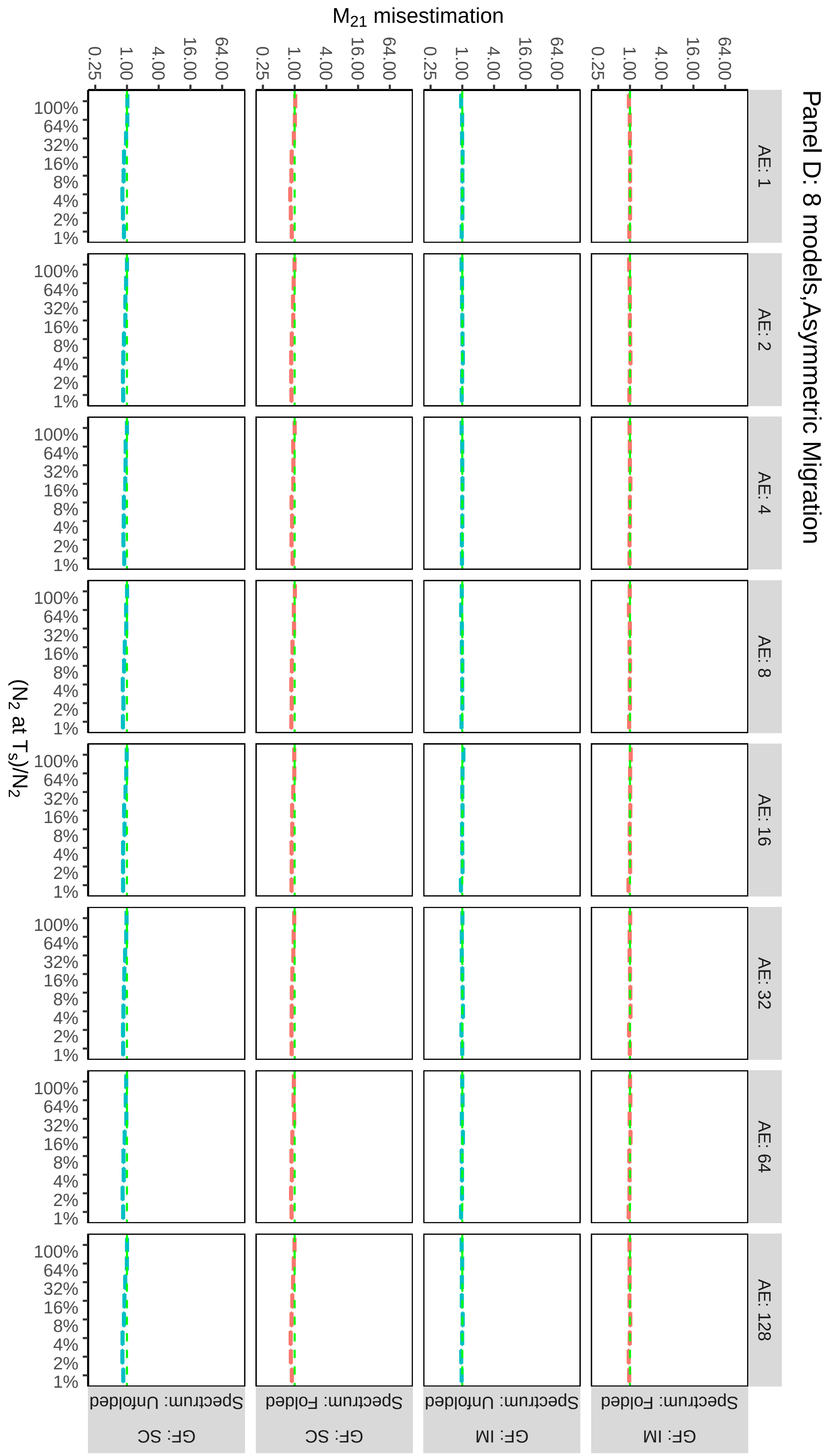
Panel D: 8 models,Asymmetric Migration
M21 misestimation
0.25
1.00
4.00
16.00
64.00
AE: 1
AE: 2
AE: 4
AE: 8
AE: 16
AE: 32
AE: 64
AE: 128
(N2 at Ts)/N2
100%
64%
32%
16%
8%
4%
2%
1%
Spectrum: Unfolded
GF: SC
Spectrum: Folded
GF: SC
Spectrum: Unfolded
GF: IM
Spectrum: Folded
GF: IM

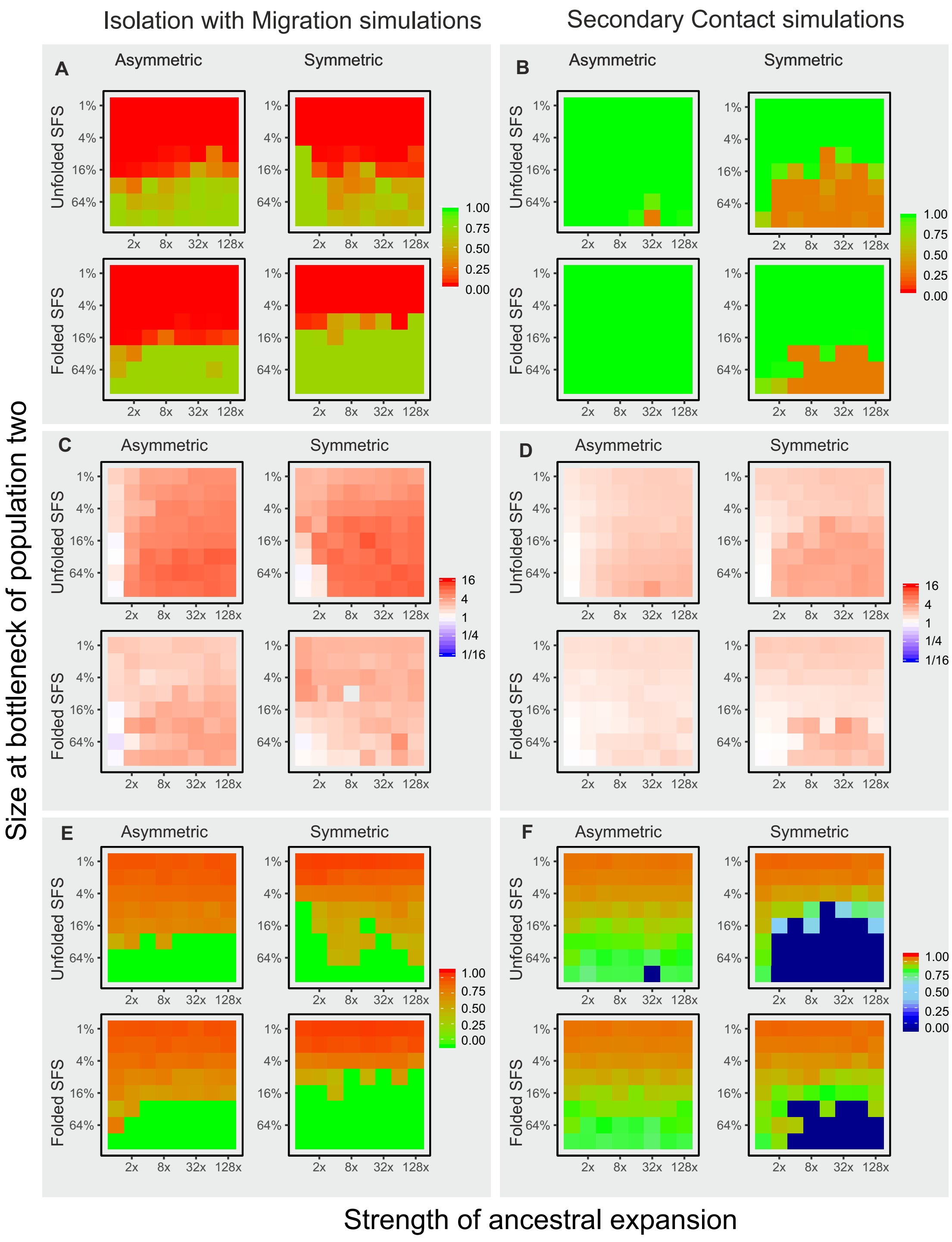


**Supplementary Fig. 16:** Model choice and parameter misestimation for the simple IM-SC models for all simulations of the smaller data sets (100 000 loci). Left panels (**A, C, E**) show results from simulations with constant migration (IM), right panels (**B, D, F**) show results from simulation with a period of strict isolation (SC). Within each panel, results are shown for simulations with symmetric and asymmetric migration, and for model estimates using the folded or unfolded jAFS. Within each panel, each graph represents the values for all 64 simulations as per Fig. 1. Panel **A** & **B** show weight of evidence for the correct gene flow scenario (0-1). Panel **C** & **D** show misestimation of the parameter $T_S$ as $T_S$ model / $T_S$ simulation. Panels **E** and **F** show the estimated proportion of the divergence time for which the model inferred strict isolation (green represents the correct time, i.e. 0 for IM model and 0.75 for SC models). This figure is the equivalent of Fig. 2 in the main manuscript, but representing values for the smaller simulated datatsets.

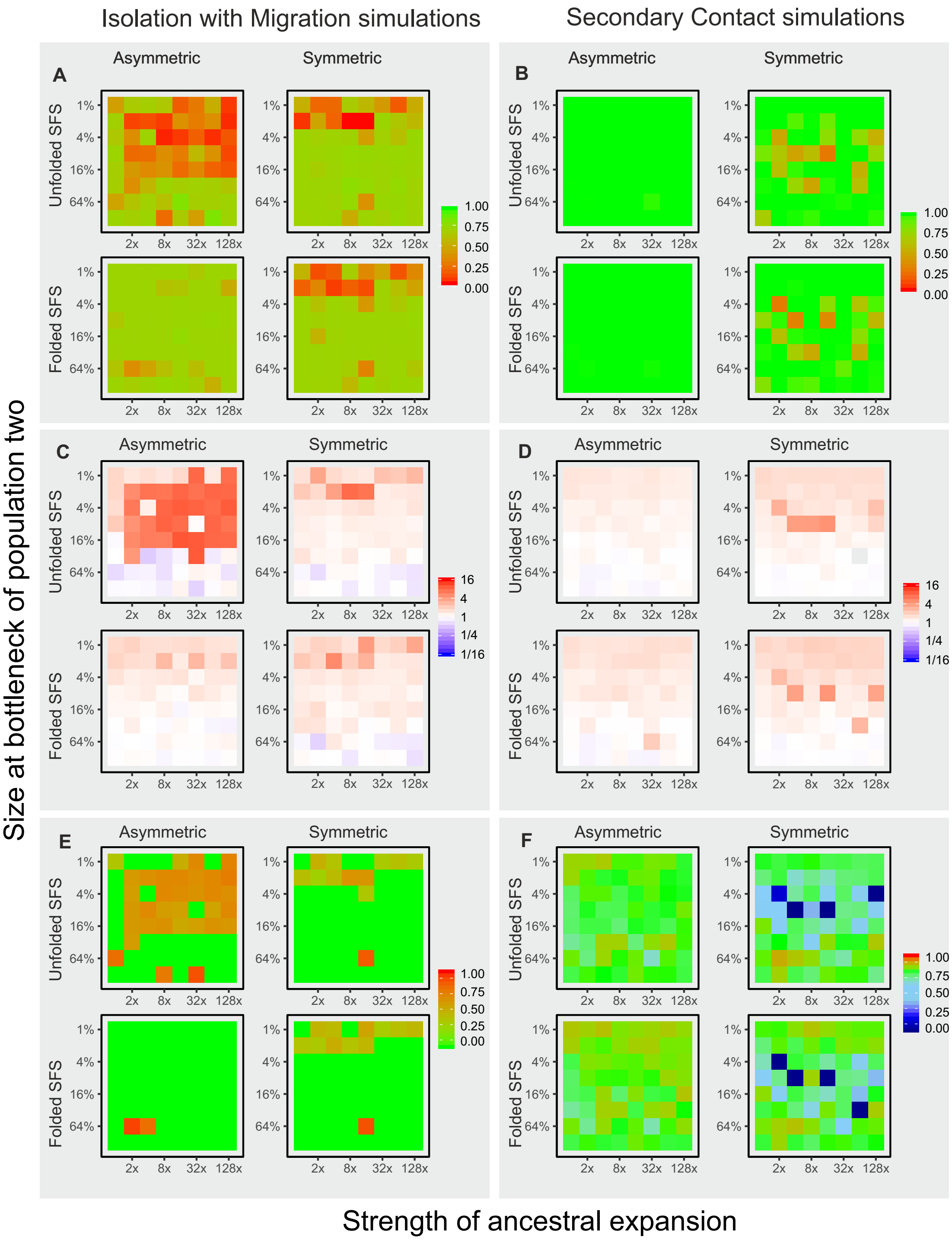


**Supplementary Fig. 17:** Model choice and parameter misestimation for all 8 divergence models for all simulations of the smaller data sets (100 000 loci). Left panels (**A, C, E**) show results from simulations with constant migration (IM), left panels (**B, D, F**) show results from simulation with a period of strict isolation (SC). Within each panel, results are shown for simulations with symmetric and asymmetric migration, and for model estimates using the folded or unfolded jAFS. Within each panel, each graph represents the values for all 64 simulations as per Fig. 1. Panel **A** & **B** show weight of evidence for the correct gene flow scenario (0-1). Panel **C** & **D** show misestimation of the parameter $T_S$ as $T_S$ model / $T_S$ simulation). Panels **E** and **F** show the estimated proportion of the divergence time for which the model inferred strict isolation (green represents the correct time, i.e. 0 for IM model and 0.75 for SC models). This figure is the equivalent of Supplementary Fig. 2, but representing values for the smaller simulated data sets.

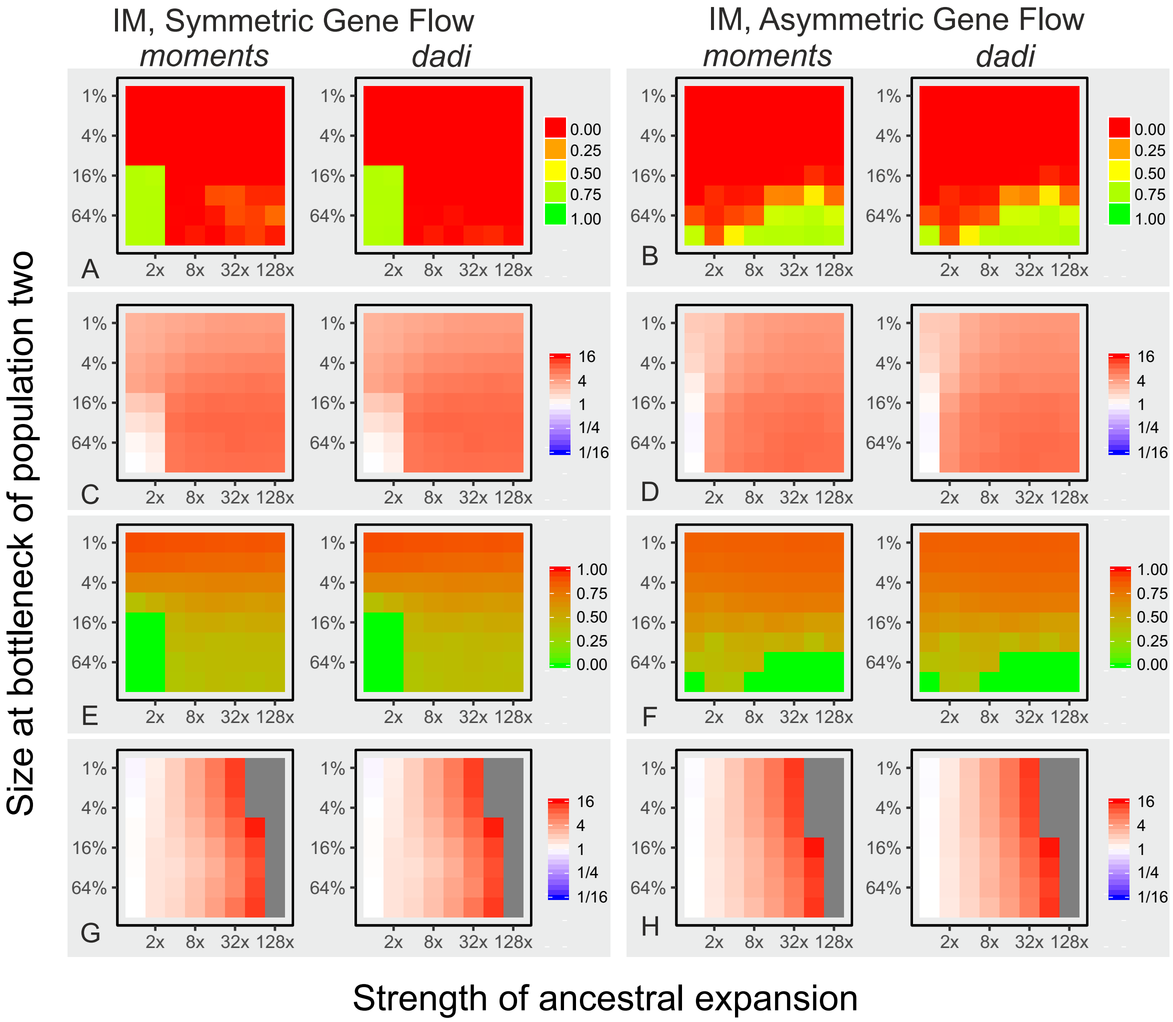


**Supplementary Fig. 18:** Comparison of model choice and parameter estimates of the IM and SC simple models in *moments* and *dadi* for IM simulations (1 million loci) under a scenario of symmetric (left panels: **A**, **C**, **E**, **G**) and asymmetric (right panels: **B**,**D**,**F**,**H**) gene flow. Panel **A** and **B** show weight of evidence for the correct gene flow scenario (0-1). Panel **C** and **D** show misesistemation of the parameter $T_S$ (calculated as $T_S$ model / $T_S$ simulation) for the best fitting model. Panels **E** and **F** show the estimated proportion of the divergence time for which the best fitting model inferred strict isolation (green represent the correct time, i.e. 0 for IM model). Panels **G** and **H** show the misestimaton of the ancestral population size $N_{ANC}$ ($N_{ANC}$ model / $N_{ANC}$ simulations). Notice that *moments* and *dadi* gave nearly identical results. Grey squares represent off-scale values.

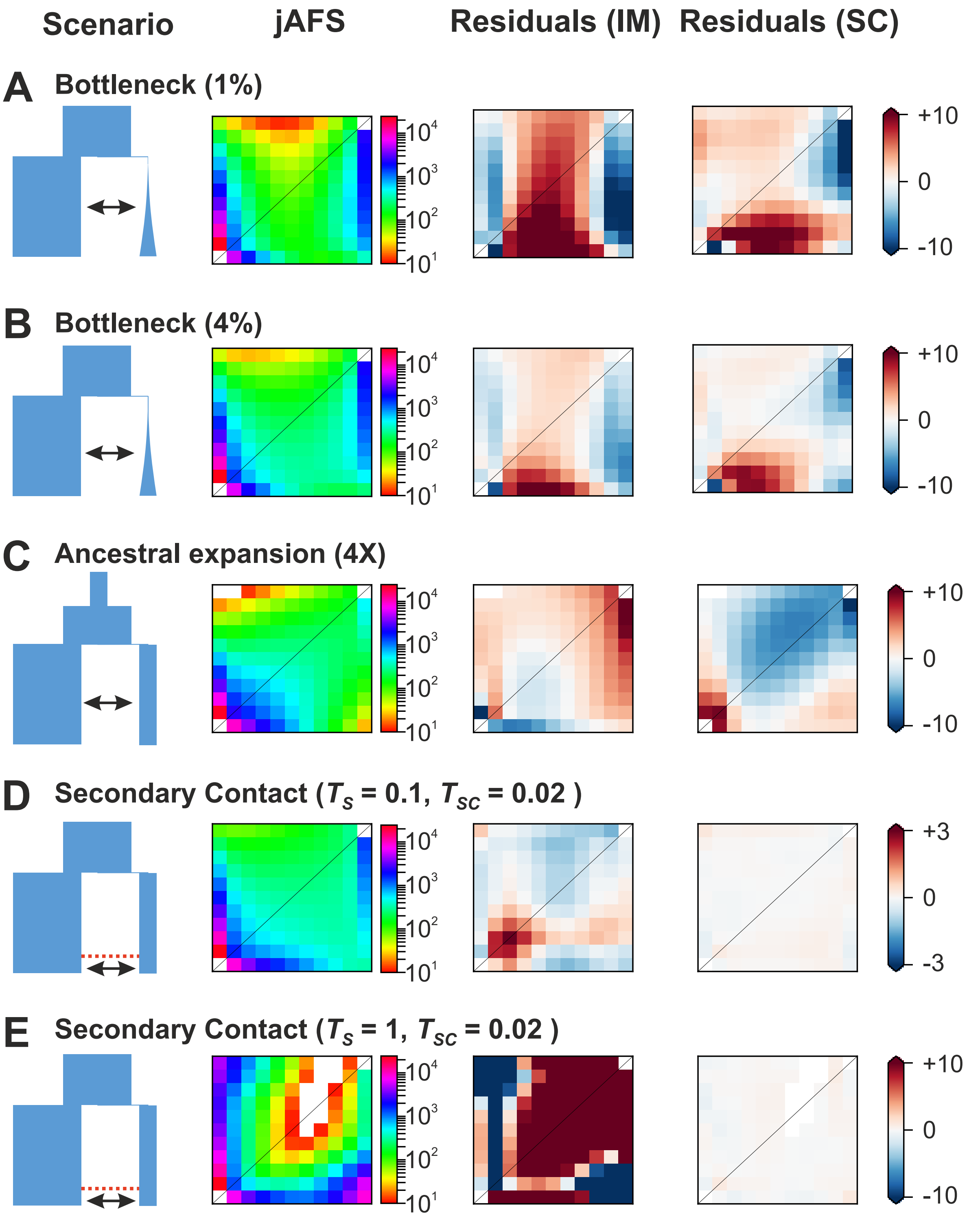


**Supplementary Fig. 19:** Examples of residual plots from optimized IM and SC models under a range of different demographic scenarios. All scenarios, with the exception of panel **D**, come from the recent divergence scenarios depicted in Fig. 1B. $T_S = 0.05$ for IM scenarios (**A,B,C**) and $T_S = 0.1$ for the SC scenario in **C**. In panel **D** we show an example of a SC scenario with a long period of strict isolation ($T_S = 1$), all other parameters are as per other recent divergence scenario. For each panel we show the demographic scenario (left, population sizes are in scale) and the observed jAFS from the simulations (jAFS), projected down to 10 alleles for ease of visualization. We then show the residual plots from the optimized IM and SC models. In all scenarios, SC showed strong statistical support. Indeed, SC models fit the data considerably better, not only when the data were simulated under a SC scenario (**D & E**), but only when there was a strong **A** or mild **B** unaccounted bottleneck, and to a lesser degree when there was an unaccounted ancestral expansion **C**.

**Supplementary Fig. 20:** Genomic landscape of differentiation between North Sea and Baltic Sea turbots across all chromosomes. The lines represent patterns of genetic diversity calculated in non-overlapping windows of 250 kb. The red line represents $\pi$ in the Baltic Sea, the blue line $\pi$ in the North Sea, and the black line represents $d_{xy}$. Dots along the chromosomes represent q values (FDR) on a negative log scales from fastPCA genome scan for selection for each individual SNP, the dotted line remarks the 0.1 FDR and green dots represents significant outliers according to the HHM test.

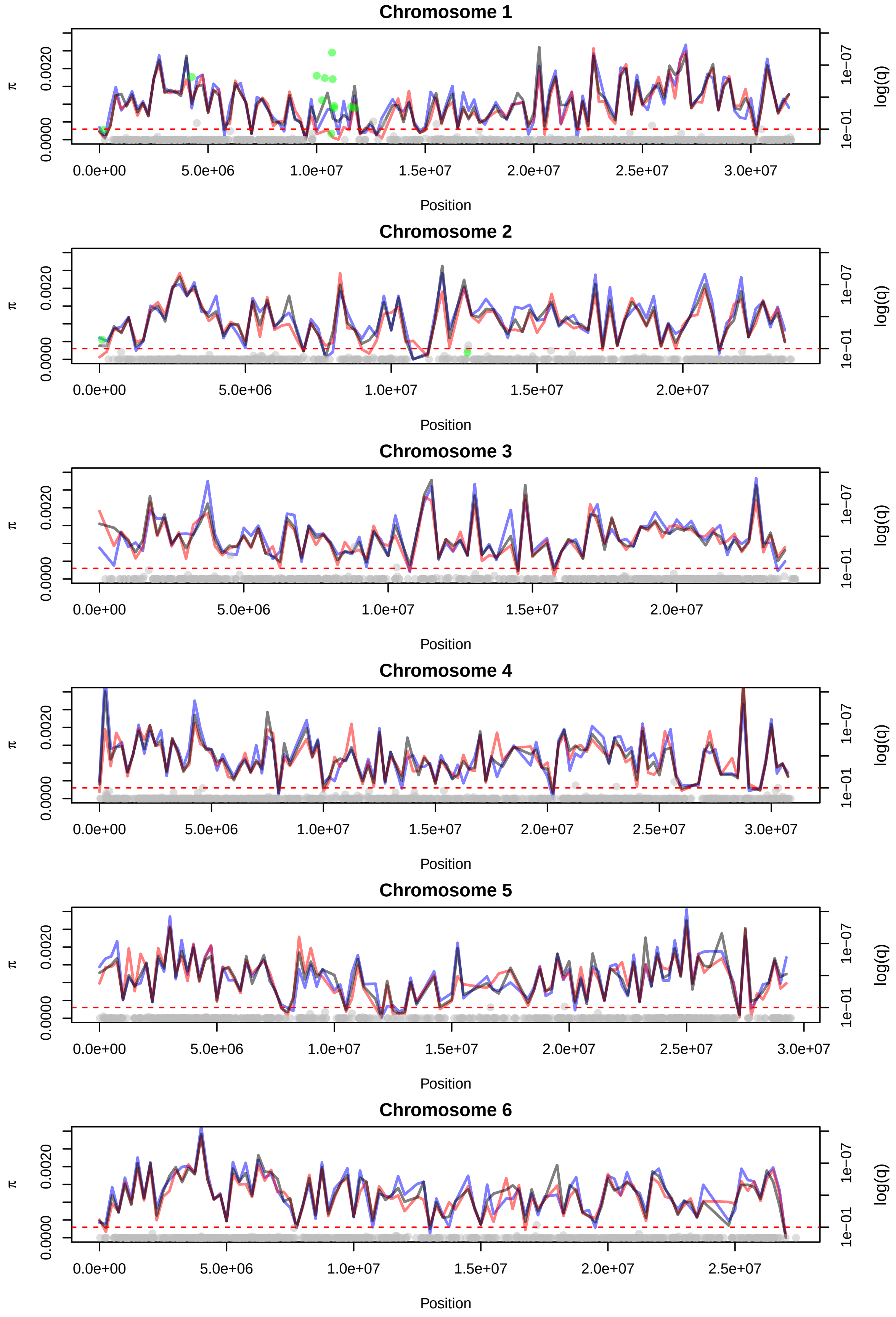

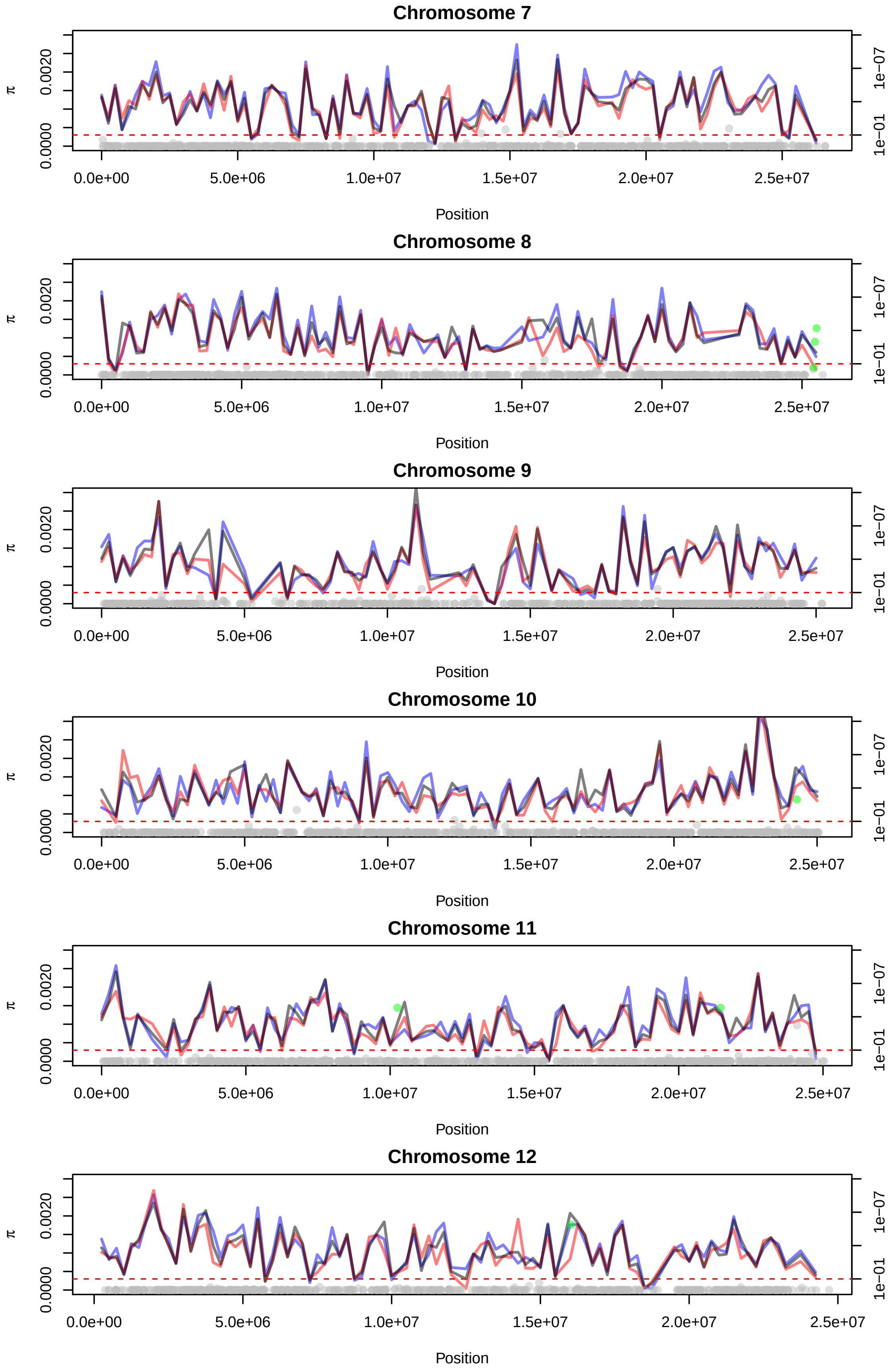
Chromosome 7
π
0.0000
0.0020
1e−01
1e−07
log(q)
0.0e+00
5.0e+06
1.0e+07
1.5e+07
2.0e+07
2.5e+07
Position
Chromosome 8
π
0.0000
0.0020
1e−01
1e−07
log(q)
0.0e+00
5.0e+06
1.0e+07
1.5e+07
2.0e+07
2.5e+07
Position
Chromosome 9
π
0.0000
0.0020
1e−01
1e−07
log(q)
0.0e+00
5.0e+06
1.0e+07
1.5e+07
2.0e+07
2.5e+07
Position
Chromosome 10
π
0.0000
0.0020
1e−01
1e−07
log(q)
0.0e+00
5.0e+06
1.0e+07
1.5e+07
2.0e+07
2.5e+07
Position
Chromosome 11
π
0.0000
0.0020
1e−01
1e−07
log(q)
0.0e+00
5.0e+06
1.0e+07
1.5e+07
2.0e+07
2.5e+07
Position
Chromosome 12
π
0.0000
0.0020
1e−01
1e−07
log(q)
0.0e+00
5.0e+06
1.0e+07
1.5e+07
2.0e+07
2.5e+07
Position

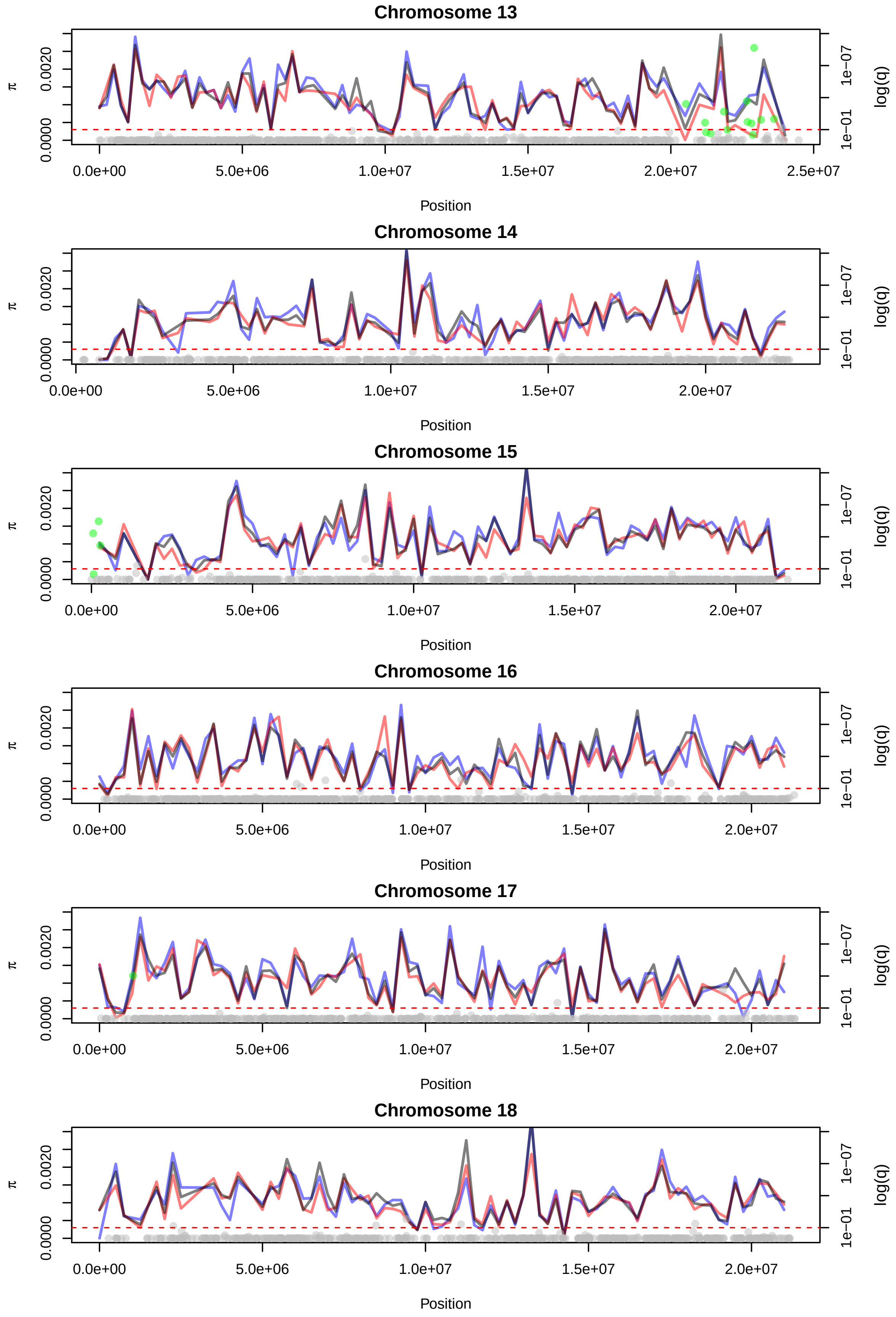
Chromosome 13
π
0.0000
0.0020
0.0e+00
5.0e+06
1.0e+07
1.5e+07
2.0e+07
2.5e+07
Position
1e−01
1e−07
log(q)
Chromosome 14
π
0.0000
0.0020
0.0e+00
5.0e+06
1.0e+07
1.5e+07
2.0e+07
Position
1e−01
1e−07
log(q)
Chromosome 15
π
0.0000
0.0020
0.0e+00
5.0e+06
1.0e+07
1.5e+07
2.0e+07
Position
1e−01
1e−07
log(q)
Chromosome 16
π
0.0000
0.0020
0.0e+00
5.0e+06
1.0e+07
1.5e+07
2.0e+07
Position
1e−01
1e−07
log(q)
Chromosome 17
π
0.0000
0.0020
0.0e+00
5.0e+06
1.0e+07
1.5e+07
2.0e+07
Position
1e−01
1e−07
log(q)
Chromosome 18
π
0.0000
0.0020
0.0e+00
5.0e+06
1.0e+07
1.5e+07
2.0e+07
Position
1e−01
1e−07
log(q)

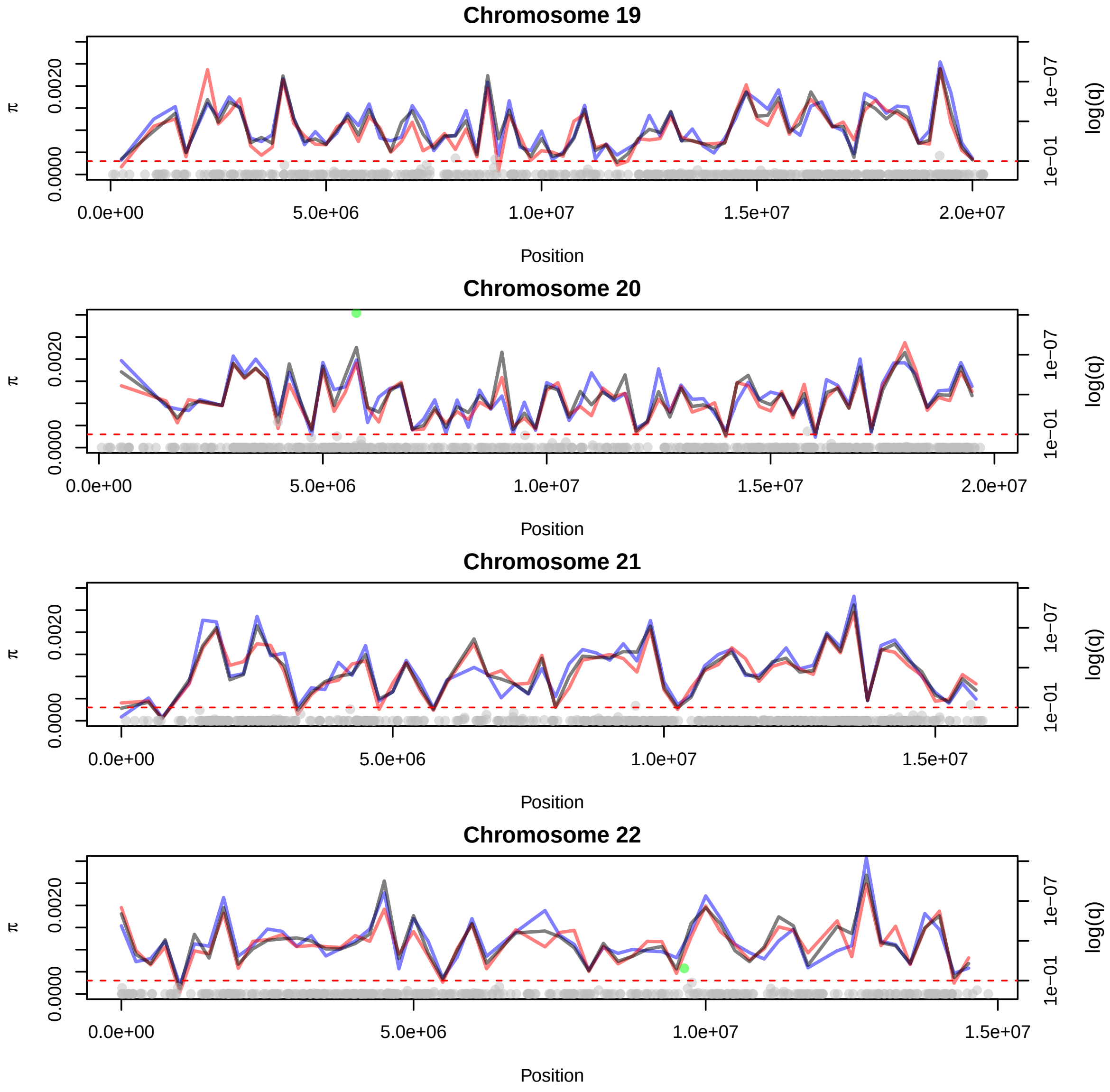
Chromosome 19
π
0.0000
0.0020
log(q)
1e−01
1e−07
0.0e+00
5.0e+06
1.0e+07
1.5e+07
2.0e+07
Position
Chromosome 20
π
0.0000
0.0020
log(q)
1e−01
1e−07
0.0e+00
5.0e+06
1.0e+07
1.5e+07
2.0e+07
Position
Chromosome 21
π
0.0000
0.0020
log(q)
1e−01
1e−07
0.0e+00
5.0e+06
1.0e+07
1.5e+07
Position
Chromosome 22
π
0.0000
0.0020
log(q)
1e−01
1e−07
0.0e+00
5.0e+06
1.0e+07
1.5e+07
Position
